# Dualsteric and dual-acting modulation of muscarinic receptors by antagonist KH-5

**DOI:** 10.64898/2026.01.15.699667

**Authors:** Alena Janoušková-Randáková, Eva Dolejší, Nikolai Chetverikov, Jan Jakubík

## Abstract

**Background and purpose:** Muscarinic acetylcholine receptors are key therapeutic targets, and ligands engaging both orthosteric and allosteric sites may offer improved selectivity and efficacy. The muscarinic antagonist KH-5 displays functional antagonistic potency exceeding its binding affinity, suggesting a non-classical mechanism of action. Here, we investigated whether KH-5 acts as a dualsteric antagonist and defined its mode of interaction with muscarinic receptors.

**Experimental approach:** Functional responses at human M_1_ and M_2_ receptors expressed in CHO cells were assessed using inositol phosphate accumulation and [^35^S]GTPγS binding, respectively. Radioligand binding studies employed orthosteric antagonists and agonists in combination with KH-5 and classical allosteric modulators. Data were analysed using competitive, allosteric, and dualsteric binding and operational models. Molecular docking, molecular dynamics simulations, and site-directed mutagenesis were used to identify structural determinants of KH-5 binding.

**Key results:** KH-5 antagonised responses to multiple agonists in a saturable and probe-dependent manner consistent with an allosteric interaction. However, KH-5 did not decrease maximal response to agonists, contradicting simple allosteric antagonism. At M_2_ receptors, antagonism was largely competitive. Binding studies revealed transient enhancement of agonist binding at M_1_ receptors at nanomolar concentrations of KH-5, best described by a dualsteric binding model involving independent orthosteric and ectopic site interactions. KH-5 did not bind to the classical muscarinic allosteric site at the second extracellular loop but interacted with an extracellular vestibule site, supported by molecular modelling and mutation of key residues.

**Conclusions and implications:** At M_1_ receptors, the most parsimonious model among those tested combines orthosteric competition with an ectopic/allosteric component. At M_2_ receptors, KH-5 behaves predominantly as an orthosteric antagonist under the present conditions, although a weak or probe-specific allosteric component cannot be excluded.

**Summary:** *What is already known:* Muscarinic receptors are therapeutic targets with conserved orthosteric binding sites. Allosteric or dualsteric ligands may improve receptor-subtype selectivity.

*What this study adds:* KH-5 shows dualsteric, dual-acting modulation at M_1_ receptors. At M_2_ receptors, KH-5 behaves mainly as a competitive antagonist.

*Clinical significance:* Dualsteric muscarinic antagonists may enable subtype-selective anticholinergic drug development. KH-5 provides a framework for designing mixed orthosteric/allosteric ligands.

## Introduction

Muscarinic acetylcholine receptors represent a family of five metabotropic G-protein-coupled receptors (M_1_–M_5_) that mediate slow modulatory responses to acetylcholine throughout the central and peripheral nervous systems (Bonner et al., 1987). These receptors exhibit distinct tissue distributions and G-protein coupling selectivity: the M_1_, M_3_, and M_5_ subtypes preferentially activate phospholipase C and mobilise intracellular calcium via G_q/11_ proteins, whereas M_2_ and M_4_ receptors inhibit adenylyl cyclase and modulate ion channel conductance through G_i/o_-coupled pathways. Muscarinic receptors are central to several of the most fundamental physiological functions, including learning and memory consolidation, motor control, nociception, regulation of sleep-wake cycles, and cardiovascular homeostasis (Eglen, 2012).

The pathophysiological role of dysfunction in muscarinic cholinergic transmission has motivated considerable therapeutic interest in selective modulation of muscarinic receptor subtypes. Muscarinic antagonists, traditionally used as broad-spectrum anticholinergics, are currently being refined into subtype-selective agents to treat various disorders while minimising the side effects associated with non-selective blockade.

Muscarinic antagonists play a critical role in managing seizures induced by organophosphate exposure, which causes muscarinic hyperstimulation leading to status epilepticus (SE) (Miller et al., 2017). The M_1_ receptor subtype is vital for the early phases of seizure generation. Pretreatment with the selective M_1_ antagonist VU0255035 significantly suppresses seizure severity and prevents the development of SE in rats (Miller et al., 2017). There is growing interest in using anticholinergics to prevent epilepsy following traumatic brain injury (Sanabria et al., 2022). In animal models, agents like scopolamine and biperiden administered during the acute phase after injury have been shown to decrease the frequency and severity of spontaneous recurrent seizures. Biperiden, a relatively specific M_1_ antagonist, is currently the only anticholinergic being investigated in clinical trials of post-traumatic epilepsy (Sanabria et al., 2022).

The advantage of muscarinic antagonists, like scopolamine, in treating depression is defined by their rapid onset of action, contrasting with the weeks required for typical antidepressants, including those with treatment-resistant depression. Preclinical data suggest that the action of scopolamine is mediated by the medial prefrontal cortex and specifically involves the M_1_ receptor (Navarria et al., 2015). However, while scopolamine showed promise in mixed cohorts, a recent randomised double-blind placebo-controlled trial in a bipolar disorder cohort found it inefficient (Miravalles et al., 2022).

Anticholinergics were among the first treatments for Parkinson’s disease (PD) and remain useful for treating resting tremors and rigidity (Xiang et al., 2012; LeWitt et al., 2024). Standard treatments like trihexyphenidyl and benztropine are non-selective, leading to dose-limiting side effects like cognitive impairment and blurred vision. Several lines of evidence suggest that the M_4_ receptor is the primary subtype responsible for the antiparkinsonian and locomotor-stimulating effects of scopolamine. Selective M_4_ antagonist VU6021625 has shown antiparkinsonian and antidystonic efficacy in rodent models (Moehle and Conn, 2019). In genetic models of dystonia, excessive cholinergic transmission overstimulates M_1_ receptors, leading to impaired synaptic plasticity. Selective M_1_ antagonists have been shown to rescue these plasticity deficits (Eskow Jaunarajs et al., 2015).

Muscarinic receptors, particularly M_4_ and M_5_, are key modulators of the mesolimbic dopaminergic reward circuitry involved in addiction (Gunter et al., 2018; Walker et al., 2020; Garrison et al., 2022). Deletion of the M_5_ receptor reduces the reinforcing effects and relative strength of cocaine in animal models (Thomsen et al., 2005). The M_5_-selective antagonist VU6019650 has been shown to inhibit oxycodone self-administration in rats at doses that do not impair motor output (Garrison et al., 2022). M_5_ receptor inhibition also reduces alcohol self-administration (Walker et al., 2021).

Overactive bladder (OAB) results from pathological detrusor hyperactivity and altered afferent signalling that can be partially reversed by M₃ receptor antagonism (Kwon et al., 2024). Chronic obstructive pulmonary disease (COPD) and asthma, characterised by airway smooth muscle contraction and bronchoconstriction driven by overexcitation of M₃ receptors, represent the primary clinical indication for long-acting muscarinic antagonists (Moulton and Fryer, 2011; Gosens and Gross, 2018). Thus, the pharmacotherapeutic potential of muscarinic antagonists is broad.

In our previous work, we synthesised the muscarinic antagonist KH-5 (1-{2-[4-(hexyloxy)benzoyloxy]ethyl}-1-methyl-1,2,3,6-tetrahydropyridin-1-ium iodide) (Boulos et al., 2018) (Figure 1), which exhibits sustained antagonistic properties. Notably, the potency of KH-5 to antagonise functional response to carbachol is greater than its binding affinity (Randáková et al., 2018). This discrepancy indicates that the high potency of KH-5 may result from positive cooperativity with an agonist bound to the receptor, meaning it has a greater affinity for the receptor-agonist complex than for the free receptor itself. Therefore, we hypothesised that KH-5 acts as a positive allosteric modulator (PAM) antagonist (Kenakin and Strachan, 2018). The functional antagonism observed correlates with the affinity for the receptor-agonist complex, whereas the binding affinity measured reflects interaction with the unoccupied receptor. Since binding of KH-5 is mutually exclusive with the orthosteric antagonist N-methylscopolamine (NMS), it may be considered a dualsteric antagonist. Dualsteric ligands are small molecules capable of binding to both the orthosteric and allosteric sites of a receptor, unlike ligands that target only one site (Antony et al., 2009). Binding to the conserved orthosteric site ensures good efficacy and signalling specificity, while interaction with the less conserved allosteric site confers subtype selectivity. Classical bitopic dualsteric ligands employ two different pharmacophores, usually connected by a linker. One of the pharmacophores binds to the orthosteric site, the other to the allosteric site. Bitopic ligands may bind to both sites concurrently. In contrast, KH-5, as a single pharmacophore ligand, is hypothesised to bind these sites independently (Figure 1). Thus, KH-5 can bind to the orthosteric binding site and competitively antagonise functional response to agonists. It can also bind to the allosteric site, exerting positive binding cooperativity with an agonist, while antagonising functional response with higher potency.

**Figure 1.**
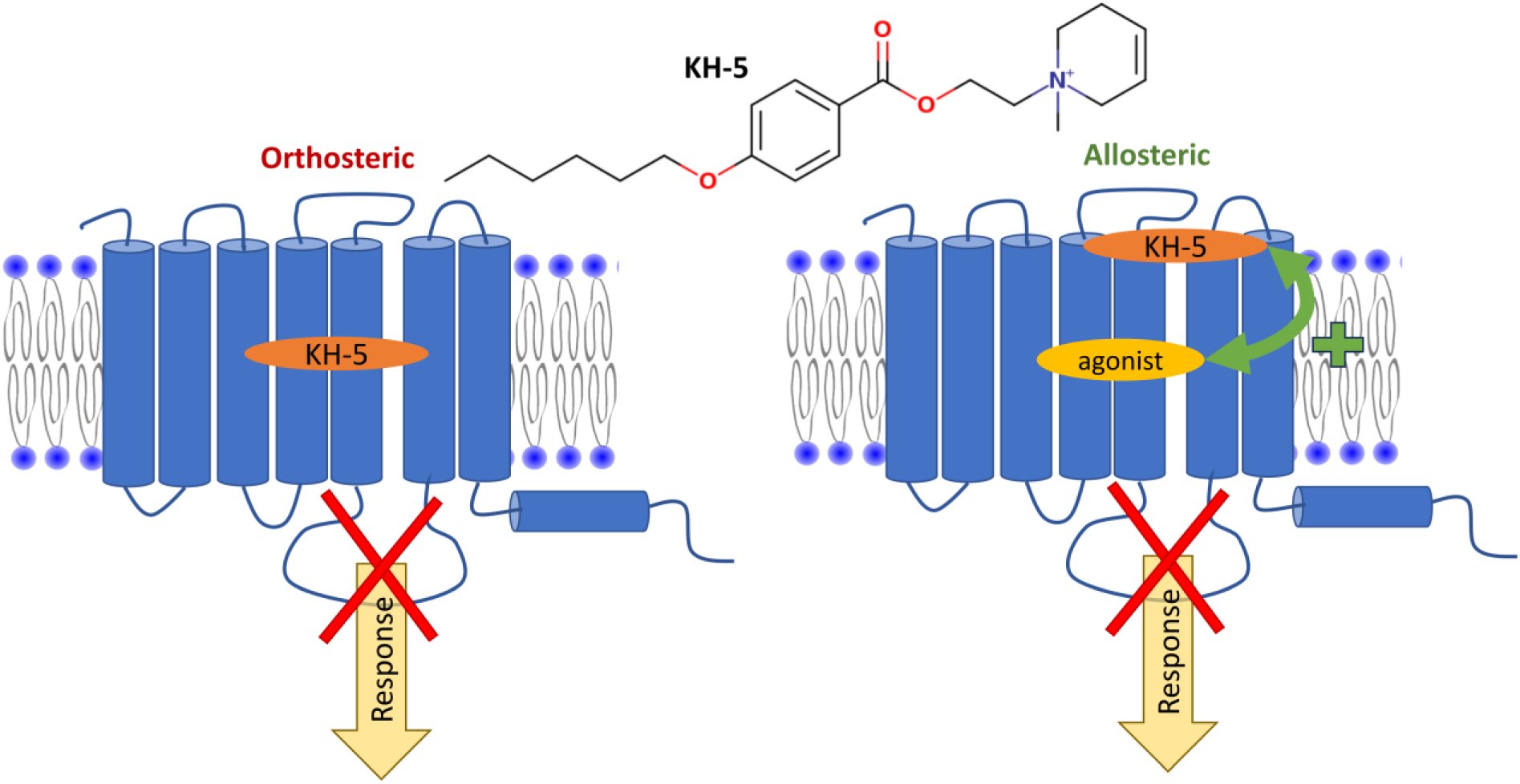
Scheme of KH-5 and the putative mechanism of KH-5 antagonism. KH-5 can bind both to the orthosteric site (left) and allosteric site (right). By blocking the orthosteric site, KH-5 competitively antagonises functional response to agonists. From the allosteric site, KH-5 exerts positive binding cooperativity with an agonist while antagonising functional response.

In the presented work, we investigate the functional antagonism of KH-5 with various agonists, examine its binding interactions with orthosteric and allosteric ligands across different experimental setups, and analyse the impact of receptor mutations on KH-5 binding.

## Methods

### Materials

Common chemicals were purchased from Sigma (Prague, CZ) in the highest available purity. Radiolabelled [^35^S]GTPγS was Perkin Elmer NEG030H, and [^3^H]acetylcholine, [^3^H]oxotremorine-M, [^3^H]myo-inositol, and [^3^H]N-methyl scopolamine ([^3^H]NMS) were American Radiochemical Company ART-0317, ART-1805, ART-0116 and ART-1214, respectively, purchased from Lacomed (Kralupy nad Vltavou, CZ). Solid propylbenzilylcholine mustard was NLP-014 Lot 2389-088 from NEN Products DuPont (Boston, MA).

### Cell culture and membrane preparation

The Chinese hamster ovary (CHO) cells stably expressing individual human variants (M_1_ to M_5_) of muscarinic receptors (RRID: CVCL_F2AT, CVCL_F2AY, CVCL_F2AZ, CVCL_F4BI, CVCL_F4BJ) (Missouri S&T cDNA Resource Center, Rolla, MO, USA) were grown to confluence in 75 cm^2^ flasks in Dulbecco’s modified Eagle’s medium supplemented with 10% fetal bovine serum and 2 million cells were subcultured in 100 mm Petri dishes. The medium was supplemented with 5 mM butyrate for the last 24 hours of cultivation to increase receptor expression. Cells were detached by mild trypsinisation on day 5 after subculture. For the cultivation of transiently transfected cells, see “Construction of mutant receptors” below. Detached cells were washed twice in 50 ml of phosphate-buffered saline and centrifuged for 3 min at 250 x g. Harvested cells were stored at -20°C overnight. Thawed cells were suspended in 20 ml of ice-cold incubation medium (100 mM NaCl, 20 mM Na-HEPES, 10 mM -MgCl_2_, pH 7.4) supplemented with 10 mM EDTA and homogenised on ice by two 30-s strokes using a Polytron homogeniser (Ultra-Turrax; Janke & Kunkel GmbH & Co. KG, IKA-Labortechnik, Staufen, Germany) with a 30-s pause between strokes. Cell homogenates were centrifuged for 5 min at 1000×g to remove whole cells and cell nuclei. The resulting supernatants were centrifuged for 30 min at 30,000×g. Pellets were suspended in a fresh incubation medium, incubated on ice for 30 min, and centrifuged again. The resulting membrane pellets were kept at -80 °C until assayed within 10 weeks.

### Protein content determination

Proteins were determined using Peterson’s modification (Peterson, 1977) of Lowry’s method (Lowry et al., 1951).

### Construction of mutant receptors

Plasmids coding muscarinic receptors were obtained from the Missouri S&T cDNA Resource Center (Rolla, MO, USA) and used to generate sequences with the desired amino acid substitution using Q5 Site-Directed Mutagenesis Kit (NEB, via BioTek, Praha, Czech Republic). Primers were designed using NEBaseChanger (https://nebasechanger.neb.com/). CHO-K1 cells (RRID: CVCL_0214) were transfected with desired plasmids using linear polyethylenimine (PEI) (Polysciences, Germany). Approximately 2 million cells were seeded in 10 ml of a 1:1 mixture of Dulbecco’s modified Eagle’s and Ham’s F-12 with L-glutamine in a 100 mm Petri dish and incubated for 24 hours at 37°C in a humidified 5% CO_2_ atmosphere. DNA and PEI were mixed in a 1:3 ratio (12 μg DNA per 1 ml PBS: 36 μg PEI per 1 ml PBS) and incubated at room temperature for 30 min. Then 1 ml of transfection mix was added to a dish and incubated for 16 hours at 37°C in a humidified 5% CO_2_ atmosphere. Then DMEM was removed, and 8 ml of fresh DMEM was added, and the cells were incubated for an additional 24 hours.

### Accumulation of inositol phosphates (IP_X_)

Cells were seeded in 96-well plates, 20 thousand cells per well in 100 µl of DMEM. The next day, DMEM was removed, and 50 µl of DMEM supplemented with 30 nM [^3^H]myo-inositol was added for 12 hours. Then, the cells were washed with Krebs-HEPES buffer (KHB; 138 mM NaCl; 4 mM KCl; 1.3 mM CaCl_2_; 1 mM MgCl_2_; 1.2 mM NaH_2_PO_4_; 10 mM glucose; 20 mM Na-HEPES; pH = 7.4) supplemented with 10 mM LiCl. Cells were preincubated at 37◦C for 30 minutes with the tested compound at the desired concentration, and then the tested agonist was added for an additional 30 minutes. Then KHB was removed, and accumulation of inositol phosphates was stopped by the addition of 50 µl of 20 % trichloracetic acid (TCA). Plates were put to 4◦C for 1 hour, then 40 µl of TCA extract was transferred to another 96-well plate, mixed with 200 µl of Rotiszint scintillation cocktail and counted in Wallac Microbeta^2^. The rest of the TCA extract was discarded, individual wells were washed with 50 µl of 20 % TCA, 50 µl of 1 M NaOH was added to each well and plates were shaken at room temperature for 15 min. Then 40 µl of NaOH lysate was transferred to another 96-well plate, mixed with 200 µl of Rotiszint scintillation cocktail and counted in Wallac Microbeta2. Levels of inositol phosphates were calculated as a fraction of soluble (TCA extract) to total (TCA extract plus NaOH lysate) radioactivity.

### Chemical inactivation

Alternatively, to reduce the receptor reserve, cells were treated with propylbenzilylcholine mustard (2-[2-chloroethyl(methyl)amino]ethyl 2-hydroxy-2,2-diphenylacetate, PBCM) immediately before measurement of accumulation of IP_X_. PBCM was diluted to the final concentration of 10 mM in methanol and kept at -20°C. It was activated by 60-min incubation in a water solution (0 .1 mM) at room temperature and then added to the cells to a final 5 nM concentration in KHB and incubated at room temperature for 2 or 5 min. The reaction was stopped by adding sodium thiosulphate to a final concentration of 1 mM, KHB containing PBCM was discarded, and cells were quickly washed twice with 100 μl of KHB.

### [^35^S]GTPγS binding

Measurement of [^35^S]GTPγS binding was carried out on membranes in 96-well plates at 30°C in the incubation medium described above supplemented with freshly prepared dithiothreitol at a final concentration of 1 mM, essentially as described previously Randáková et al., 2018). Membranes at concentrations of 10 μg of protein per well were used. The final volume was 200 μl, the final concentration of [^35^S]GTPγS was 500 pM, and the final concentration of GDP was 50 μM. Non-specific binding was determined in the presence of 1 μM unlabelled GTPγS. Total GTP binding capacity was determined in the absence of GDP. Agonist was added 15 min before [^35^S]GTPγS. Incubation with [^35^S]GTPγS was carried out for 20 min, and the free ligand was removed by filtration through GF/C glass-fibre filtration plates (Unifilter) using a Brandel cell harvester (Brandel, Gaithersburg, MD, USA). Filtration and washing with ice-cold water lasted for 9 seconds. Filters were dried, and then 40 µl of liquid scintillator Rotiszint was added to each well. The filtration plates were counted in a Wallac-Microbeta scintillation counter.

### Saturation binding

Membranes (20 -50 μg of membrane proteins per sample) were incubated in 96-well plates at 30°C for 5 h in 800 μl ([^3^H]NMS) or 1 h in 200 μl ([^3^H]acetylcholine, [^3^H]oxotremorine-M) of incubation medium at eight concentrations of radioligand 3/4 dilutions. Nonspecific binding was determined in the presence of 1 μM atropine. Incubations were terminated by filtration through GF/C glass-fibre filtration plates (Unifilter) using a Brandel cell harvester (Brandel, Gaithersburg, MD, USA). Filtration and washing with ice-cold water lasted for 6 ([^3^H]NMS) or 3 ([^3^H]acetylcholine, [^3^H]oxotremorine-M) seconds. Filters were dried, and then 40 µl of liquid scintillator Rotiszint was added to each well. The filtration plates were counted in a Wallac-Microbeta scintillation counter.

### Competition binding

Membranes (20 -50 μg of membrane proteins per sample) were incubated in 96-well plates at 30°C for 1 h in 400 μl ([^3^H]NMS) or in 200 μl ([^3^H]acetylcholine, [^3^H]oxotremorine-M) of incubation medium. In competition experiments with KH-5 only, KH-5 was added, and incubation continued for an additional 15 hours. When interaction between KH-5 and alcuronium was investigated, after one hour with [^3^H]NMS, KH-5 was added for 3 hours, and then alcuronium was added for the final 15 hours of incubation. Then Nonspecific binding was determined in the presence of 1 μM atropine. Incubations were terminated by filtration through GF/C glass-fibre filtration plates (Unifilter) using a Brandel cell harvester (Brandel, Gaithersburg, MD, USA). Filtration and washing with ice-cold water lasted for 6 ([^3^H]NMS) or 3 ([^3^H]acetylcholine, [^3^H]oxotremorine-M) seconds. Filters were dried, and then 40 µl of liquid scintillator Rotiszint was added to each well. The filtration plates were counted in a Wallac-Microbeta scintillation counter.

### Molecular modelling

#### Preparation of receptor structure

The structures of the muscarinic acetylcholine receptors were downloaded from the RCSB Protein Data Bank (https://www.rcsb.org/) using YASARA. Non-protein and nanobody molecules were deleted.

#### Ligand docking

Structures of KH-5 stereoisomers were sketched in ChemAxon MarvinSketch and imported into YASARA, where they were prepared for docking (Krieger et al., 2002). Antagonist KH-5 was docked to receptors in inactive conformations (5CXV of M_1_, 3UON of M_2_, 4DAJ of M_3_, 5DSG of M_4_ and 6OL9 of M_5_) and active conformations (6OIJ for M_1_, 4MQT for M_2_ and 8FX5 of M_4_) (Haga et al., 2012; Kruse et al., 2012, 2013; Thal et al., 2016; Maeda et al., 2019; Vuckovic et al., 2019; Burger et al., 2023). The ectopic binding site was defined as a 10 Å extended cuboid around xanomeline bound to the allosteric site at M_4_ receptor (8FX5). Other receptor subtypes were structurally aligned with 8FX5 in YASARA by MUSTANGPP (Konagurthu et al., 2006), and the ectopic site was mapped onto them. The ligands were docked to the orthosteric or ectopic binding site using the YASARA implementation of the AutoDock local search procedure for 888 poses (Trott and Olson, 2010). All poses were rescored in YASARA using AutoDock VINA’s local search, confined closely to the original ligand pose. The pose with the highest rescore value was selected for further work. The docking energies are scoring estimates used only for relative pose prioritisation; they are not to be treated as experimentally determined free energies.

#### Simulation of molecular dynamics

To evaluate KH-5 binding to the receptor and quantify its interactions with the receptor, conventional molecular dynamics (cMD) was simulated using Desmond ver. 7.6 (Shaw, 2005). First, the ligand-receptor complexes were processed in Maestro using the Protein Preparation Wizard according to Sastry et al. guidelines (Sastry et al., 2013). The simulated (OPLS force field) system, consisting of the receptor-ligand complex in 1-palmitoyl-2-oleoyl-sn-glycero-3-phosphocholine (POPC) membrane set to an OPM-oriented receptor in water and 0.15M NaCl, was constructed in Maestro using System Builder. The system was first relaxed by the standard Desmond protocol for membrane proteins, and then 120 ns γNPT (Noose-Hover chain thermostat at 300 K, Martyna-Tobias-Klein barostat at 1.01325 bar, isotropic coupling, Coulombic cutoff at 0.9 nm) molecular dynamics without restraints was simulated. MD was run three times with random initial velocities (3 independent replicas). The quality of the molecular dynamics simulation was assessed using Maestro’s simulation quality analysis tools and analysed using the Simulation Event Analysis tool. Ligand-receptor interactions were identified using the Simulation Interaction Diagram tool.

### Experimental data and statistical analysis

The data and statistical analysis comply with the recommendations of the British Journal of Pharmacology on experimental design and analysis in pharmacology (Curtis et al., 2025). Experiments were independent, using different seedings and transfection, if applicable, of CHO cells, followed by membrane preparation, if applicable. Binding experiments as well as functional assays were carried out in five independent experiments with samples in quadruplicate. Experimenters were blind to the chemical structures of the tested compounds. After subtraction of non-specific binding (binding experiments) or background/blank values (functional experiments), data were normalised to control values determined in each experiment. IC_50_ and EC_50_ values and parameters derived from them (K_I_ and K_A_) were analysed after conversion to their logarithms. All data were included in the analysis, no outliers were excluded, and the normality of distribution was checked. In statistical analysis, a value of P < 0.05 was taken as significant for all data. When testing a single concentration, the one-sample t-test was used. In multiple comparison tests, ANOVA with P < .05 was followed by Tukey’s post-test (P < 0.05). When fitting several models to the data, the goodness of fit was evaluated by the Runs test. The nested models (e.g., Equation 2 vs. Equation 3) were compared by the Extra Sum-of-Squares F-Test. The non-nested models (Equation 5 to Equation 8) were compared by Akaike Information Criterion corrected (AICc) (Sutherland et al., 2023). Statistical analysis and fitting of equations were carried out using in-house Python scripts using scipy and statmodels libraries. Sniplets of code are in the Supplementary Information. Full Python code can be downloaded from GitHub: Pharmacological analysis https://github.com/janjakubik-sr/jackonda and statistical analysis https://github.com/janjakubik-sr/pysa.

#### Saturation experiments

After subtraction of non-specific binding, data were converted to substance amounts and concentrations. Equation 1 was fitted to data from saturation experiments.

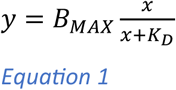

Where y is the specific radioligand binding at a concentration of the radioligand x, B_MAX_ is the maximal binding capacity, and K_D_ is the equilibrium dissociation constant of the radioligand.

#### Competition experiments

After subtraction of non-specific binding, data were normalised to the radioligand binding in the absence of competitor(s) and expressed as percentages. The following equations were fitted to the data as needed.

For simple competition, the inhibition constant (K_I_) was calculated according to Equation 2.

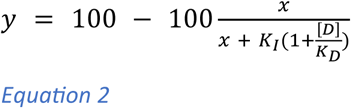

where y is specific radioligand binding at concentration x of competitor expressed as a percentage of binding in the absence of a competitor, [D] is the concentration of radioligand used, and K_D_ is its equilibrium dissociation constant determined in saturation experiments.

For competition for two sites, inhibition constants K_I1_ and K_I2_ were calculated according to Equation 3

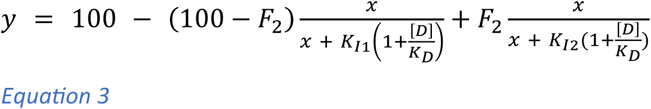

where F_2_ is the percentage of sites of K_I2_.

In case of allosteric interaction, the equilibrium dissociation constant of allosteric modulator A (K_A_) and the factor of binding cooperativity α between allosteric modulator and radiolabelled tracer were calculated according to Equation 4 (Ehlert, 1988).

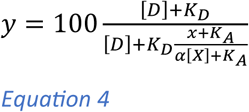

where y is specific radioligand binding at concentration x of allosteric modulator, expressed as a percentage of binding in the absence of an allosteric modulator.

In case of allosteric interaction of radiolabelled tracer with two allosteric modulators A and B competing for the same allosteric site, their respective equilibrium dissociation constants K_A_ and K_B_ and the factor of binding cooperativity between tracer and allosteric modulator B were calculated according to Equation 5 (Jakubík et al., 2019).

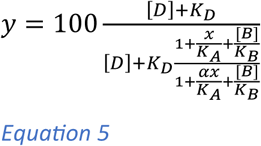

where y is specific radioligand binding at concentration x of allosteric modulator A, expressed as a percentage of binding in the absence of allosteric modulators A and B.

In case of interaction of radiolabelled tracer with allosteric modulator A and orthosteric competitor B, their respective equilibrium dissociation constants K_A_ and K_B_ and factor of binding cooperativity β between allosteric modulator A and orthosteric competitor B were calculated according to Equation 6 (Jakubík et al., 1997).

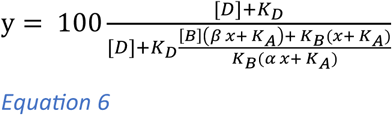

where y is specific radioligand binding at concentration x of allosteric modulator A, expressed as a percentage of binding in the absence of an allosteric modulator A and orthosteric competitor B.

For the competition of three ligands, radiolabelled tracer and two orthosteric competitors A and B, their respective equilibrium dissociation constants K_A_ and K_B_ were calculated according to Equation 7 (Jakubík et al., 2019a).

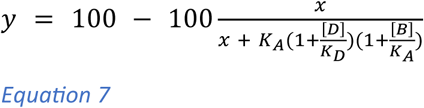

where y is specific radioligand binding at concentration x of competitor A, expressed as a percentage of binding in the absence of competitors A and B.

For dualsteric interaction between the orthosteric tracer and dualsteric ligand binding independently to both orthosteric and allosteric sites, equilibrium dissociation constants for the allosteric site K_A_ and the orthosteric site K_B_ and factors of cooperativity between tracer and dualsteric ligand α and two bound molecules of dualsteric ligand β were calculated according to Equation 8 (For derivation see Supplementary information).

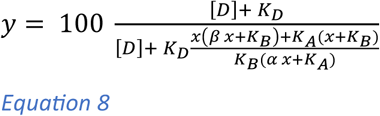

where y is the specific radioligand binding at concentration x of dualsteric ligand expressed as a percentage of binding in its absence. For derivations, see Supplementary Information.

#### Functional response

Experiments were independent. Functional assays were carried out in quadruplicate. After subtraction of non-specific binding (GTP binding assay), data were normalised to folds over basal.

Parameters of a functional response were determined by fitting Equation 9 to experimental data.

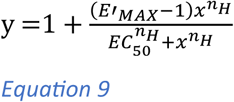

Where y is the functional response at a concentration of tested compound x, E’_MAX_ is the observed maximal response to the tested compound, EC_50_ is the half-efficient concentration, and n_H_ is the slope factor (Hill coefficient). EC_50_ values were calculated as logarithms.

To analyse functional response to an agonist according to the operational model of agonism (OMA) (Black and Leff, 1983): First, system E_MAX_ was determined as described previously (Jakubík et al., 2019b). Then, functional responses to agonists were expressed as a fraction of E_MAX_. Then Equation 10 was fitted to the data.

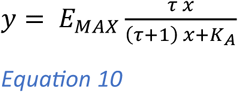

Where E_MAX_ is the maximal response of the system, K_A_ is the equilibrium dissociation constant of the agonist, and τ is its operational efficacy.

Functional response to agonist A in the presence of a competitive (orthosteric) antagonist B is described by Equation 11 (For derivation see Supplementary Information).

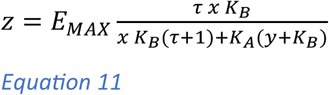

Where K_B_ is the equilibrium dissociation constant of an antagonist and y its concentration.

Functional response to agonist A in the presence of allosteric antagonist B is described by Equation 12 (Jakubík et al., 2020).

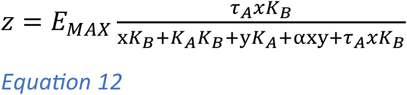

Where K_B_ is the equilibrium dissociation constant of an allosteric modulator, y is its concentration, and α is a factor of binding cooperativity between agonist and allosteric modulator according to Equation 4.

Functional response to agonist A in the presence of dualsteric ligand B (that is an orthosteric antagonist and allosteric modulator) is described by Equation 13 (For derivations see Supplementary Information).

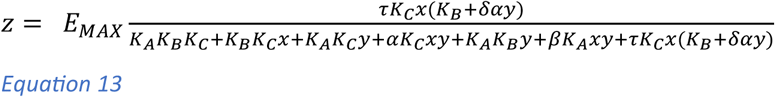

Where y is the concentration of a dualsteric ligand, K_B_ and K_C_ are equilibrium dissociation constants of dualsteric ligand for the allosteric and orthosteric binding sites, respectively, α and β are factors of binding coopertivity between molecule of dualsteric ligand bound to the allosteric binding site and molecule of agonist and dualsteric ligand bound to the orthosteric binding site, respectively, and δ is the cooperativity factor of operational efficacy. For dulasteric ligand B, which is an orthosteric and allosteric antagonist, δ = 0, and Equation 13 simplifies to Equation 14.

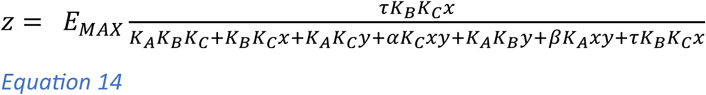

#### Schild analysis

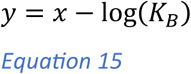

Where x is log[B], decadic logarithm of concentration of an antagonist, y is log(DR-1), decadic logarithm of dose ratio minus 1, and K_B_ is an inhibition constant of an antagonist.

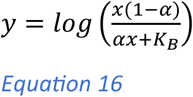

Where x is log[B], decadic logarithm of concentration of an allosteric antagonist, y is log(DR-1), decadic logarithm of dose ratio minus 1, K_B_ is an inhibition constant of an allosteric antagonist, and α is a factor of binding cooperativity between allosteric antagonist and agonist.

### Nomenclature of targets and ligands

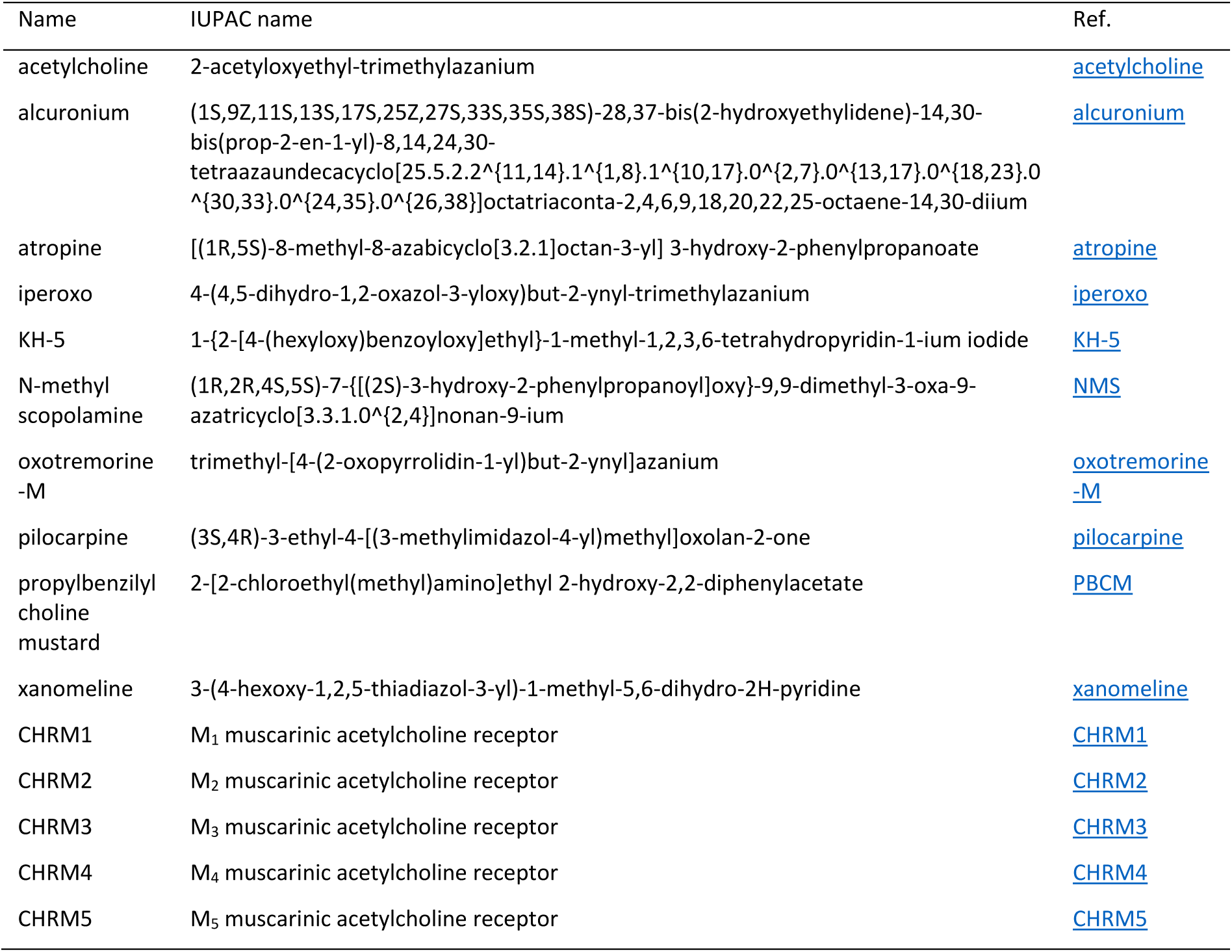

## Results

Our hypothesis posits that KH-5 functions as a dualsteric antagonist at the M_1_ muscarinic receptor. This premise is based on observations that KH-5’s ability to inhibit receptor activity surpasses what would be anticipated solely from its binding affinity, indicating a potentially complex mechanism of antagonism. To investigate this, we conducted a series of experiments assessing the functional antagonism of KH-5 with various agonists, examined its binding interactions with orthosteric and allosteric ligands across different experimental setups, and analysed the impact of receptor mutations on KH-5 binding. The results obtained were mixed; some findings aligned with the characteristics of competitive antagonism, while others suggested the involvement of non-classical, possibly allosteric, mechanisms. These outcomes highlight the complexity of KH-5’s interaction with the M_1_ receptor and suggest that its effects involve multiple modes of action.

## Functional antagonism

Firstly, we focused on understanding how KH-5 interacts with various agonists (acetylcholine, iperoxo, oxotremorine-M, pilocarpine and xanomeline) and whether its mode of action involves allosteric modulation. To this end, we measured the functional response of the receptor to different agonists by quantifying inositol phosphate accumulation (Figure 2). KH-5 was tested across a concentration range from 31.6 nM to 10 μM. Results demonstrated that KH-5 caused a dose-dependent rightward shift in the concentration-response curves for all tested agonists, indicating antagonistic activity. Notably, except for xanomeline, the antagonistic effect saturated at micromolar concentrations of KH-5, which is characteristic of allosteric interactions. Despite this, the maximum response (E_MAX_) remained unchanged (Supplementary Information Table S3), a finding that complicates the interpretation, as it contradicts allosteric antagonism, where E_MAX_ is decreased.

**Figure 2.**
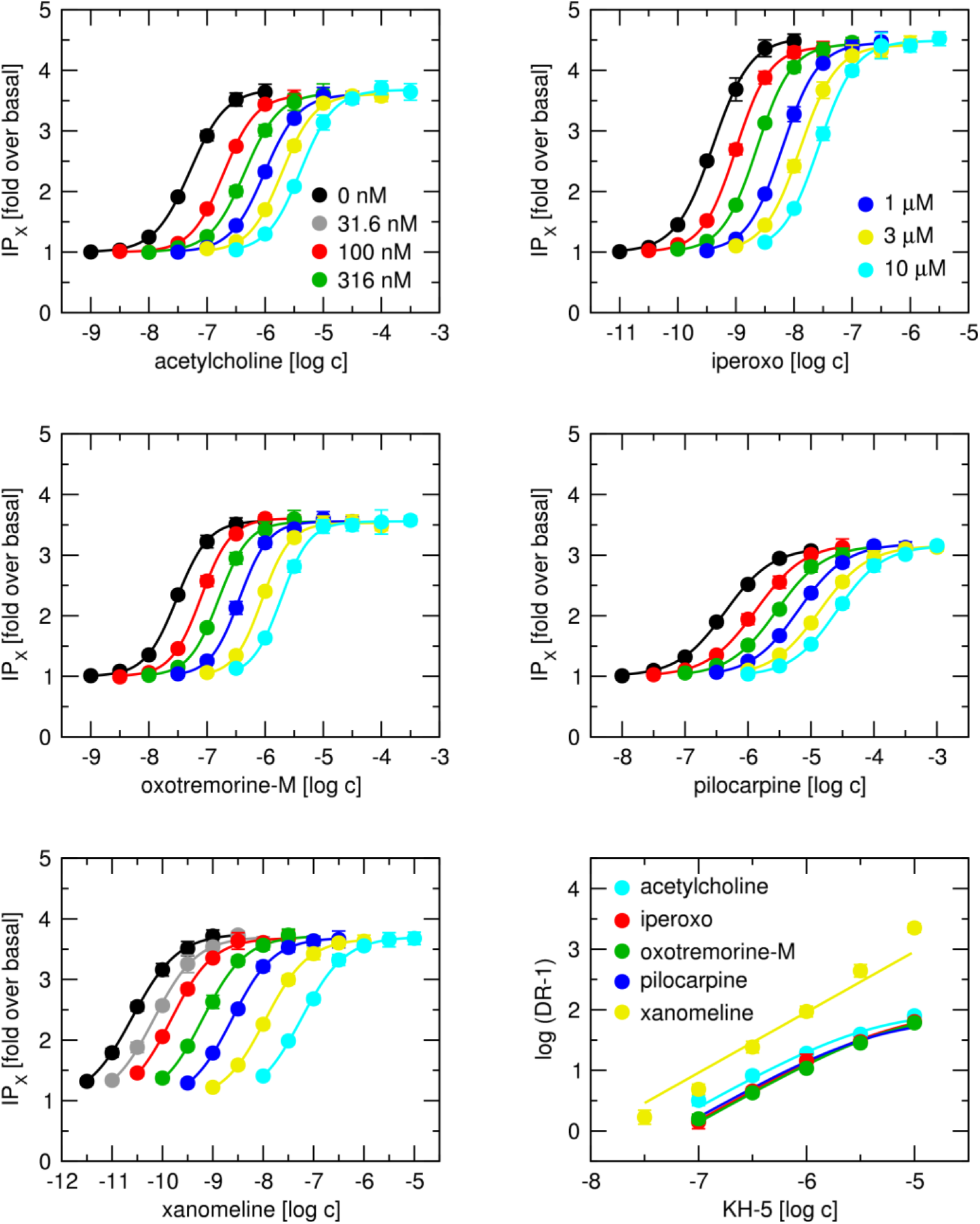
Functional responses of M_1_ receptors to agonists. Functional response of M_1_ receptors to agonists (acetylcholine, iperoxo, oxotremorine-M, pilocarpine and xanomeline) in the absence or presence of antagonist KH-5 at concentrations indicated in the legends was measured as accumulation of inositol phosphates (IP_X_) in CHO cells. Abscissa, concentration of agonist expressed as a decadic logarithm of molar concentration. Ordinate, level of IP_X_ expressed as fold over basal. Curves and their EC_50_ values were obtained by fitting Equation 9 to the data. **Bottom right**, Schild plots of KH-5 effects on functional response to individual agonists are indicated in the legend. Abscissa, concentration of KH-5 expressed as a decadic logarithm of molar concentration. Ordinate, a decadic logarithm of DR-1, where DR is the ratio of agonist EC_50_ in the presence of KH-5 to EC_50_ in the absence of KH-5. Data are means ± SD from 5 independent experiments performed in quadruplicate.

Further analysis using the Schild model (Figure 2, bottom right) revealed that the data fit better with an allosteric antagonism equation (Equation 16) than with a purely competitive model (Equation 15) (Table 1). This suggests that KH-5 interacts with the receptor in a manner that involves allosteric modulation. In the case of xanomeline, the data curve turns upward, indicating positive operational cooperativity. Additionally, the inhibition constants (K_B_) varied among the tested agonists (Supplementary Information Figure S1 and Table S4), demonstrating probe dependence—a hallmark of allosteric interactions. These findings collectively support the hypothesis that KH-5 exerts mixed antagonistic effects through the orthosteric and allosteric binding sites, thereby modulating receptor activity in a complex manner.

**Table 1.** Inhibition constants of functional responses to agonists. Inhibition constants K_B_ and factors of cooperativity were obtained by fitting Equation 15 or Equation 16 to the data from functional response assays in the presence of KH-5. Values of K_B_ are expressed as negative decadic logarithms of molar concentration. Data are means ± SD from 5 independent experiments performed in quadruplicate.

| | | $M_1$ | | $M_2$ | |
| --- | --- | --- | --- | --- | --- |
| | $pK_B$ | $\alpha$ | | $pK_B$ | |
| acetylcholine | 7.41 ± 0.07 | 0.0108 | ± 0.0012 | 5.61 | ± 0.06 |
| iperoxo | 7.21 ± 0.06 | 0.0105 | ± 0.0009 | 5.61 | ± 0.10 |
| oxotremorine-M | 7.16 ± 0.06 | 0.0106 | ± 0.0006 | 5.60 | ± 0.06 |
| pilocarpine | 7.30 ± 0.05 | 0.0134 | ± 0.0016 | 5.55 | ± 0.12 |
| xanomeline | 7.96 ± 0.09 | | | 5.81 | ± 0.08 |

In our analysis of the M_2_ receptor, we measured an increase in [^35^S]GTPγS binding to membrane preparations as an indicator of a functional response of the receptor to an agonist (Figure 3, Supplementary Information Table S5). We tested KH-5 across a concentration range from 3.16 μM to 100 μM. The results showed that KH-5 caused a concentration-dependent rightward shift in the functional-response curves without reaching saturation, while the maximum response (E_MAX_) remained unchanged. This pattern suggests a competitive interaction between KH-5 and the receptor. Furthermore, in Schild analysis (Figure 3, bottom right), the data fitted better to the competitive model (Equation 15) than the allosteric model (Equation 16) (Table 1), indicating competitive interaction. In contrast to the M_1_ receptor, Schild analysis indicated that KH-5 exhibits higher potency in antagonising the functional response only to xanomeline compared to other agonists (Supplementary Information Figure S2 and Table S6). This finding highlights probe dependence and suggests that KH-5 interacts with the receptor also through an allosteric mechanism (Figure 3, bottom right). Thus, similarly to the M_1_ receptor, at least for xanomeline, KH-5 exerts mixed antagonistic effects through the orthosteric and allosteric binding sites also at the M_2_ receptor.

**Figure 3.**
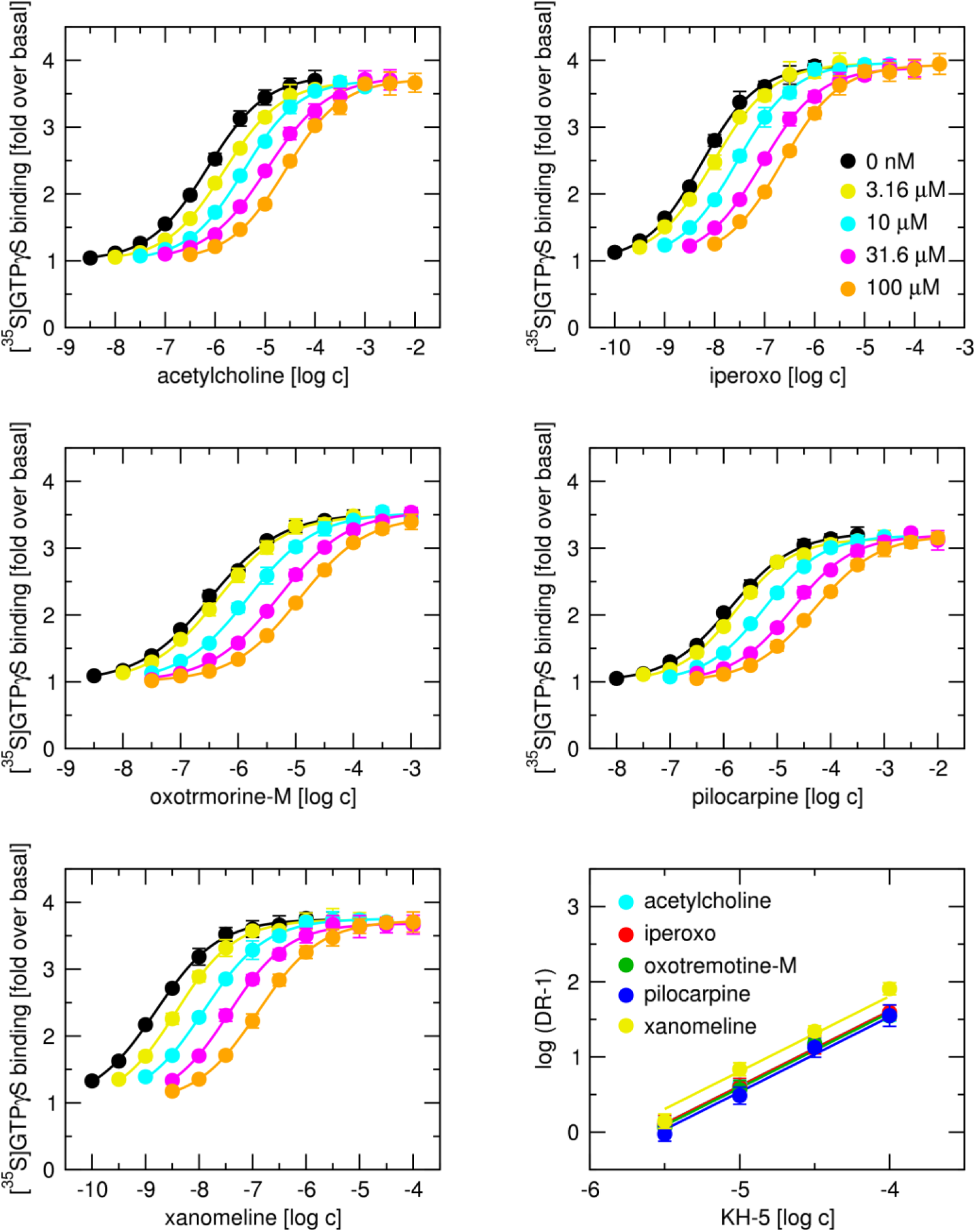
Functional responses of M_2_ receptors to agonists. Functional response of M_2_ receptors to agonists (acetylcholine, iperoxo, oxotremorine-M, pilocarpine and xanomeline) in the absence or presence of antagonist KH-5 at concentrations indicated in the legends was measured as [^35^S]GTPγS binding to membranes. Abscissa, concentration of agonist expressed as a decadic logarithm of molar concentration. Ordinate, level of [^35^S]GTPγS binding as fold over basal. Curves and their EC_50_ values were obtained by fitting Equation 9 to the data. **Bottom right**, Schild plots of KH-5 effects on functional response to individual agonists are indicated in the legend. Abscissa, concentration of KH-5 expressed as a decadic logarithm of molar concentration. Ordinate, a decadic logarithm of DR-1, where DR is the ratio of agonist EC_50_ in the presence of KH-5 to EC_50_ in the absence of KH-5. Data are means ± SD from 5 independent experiments performed in quadruplicate.

The distinguishing property of allosteric antagonism is its unsurmountable nature that manifests itself as a decrease in the observed maximum of functional response to an agonist in the presence of an allosteric antagonist. This decrease may be hard to spot for full agonists due to receptor reserve. Therefore, we measured the functional responses of the M_1_ receptor to acetylcholine and oxotremorine-M, respectively, in cells treated with an irreversible (covalent) antagonist, propyl benzilyl choline mustard (PBCM), that permanently blocks receptors, decreasing the receptor reserve. Treatment of CHO cells by PBCM brought a robust time-dependent decrease in the observed maximal response to agonists (E’_MAX_) and potency (increased EC_50_) of both agonists (Figure 4 and Supplementary Information Table S7). These data are in accordance with the irreversible blockade of receptors and a decrease in receptor reserve. However, KH-5 produced, at most, modest decreases in E′_MAX_ after receptor depletion. These changes were small relative to the PBCM-induced loss of receptor reserve and did not reveal the marked unsurmountable antagonism expected for a simple allosteric antagonist (Supplementary Information Table S7).

**Figure 4.**
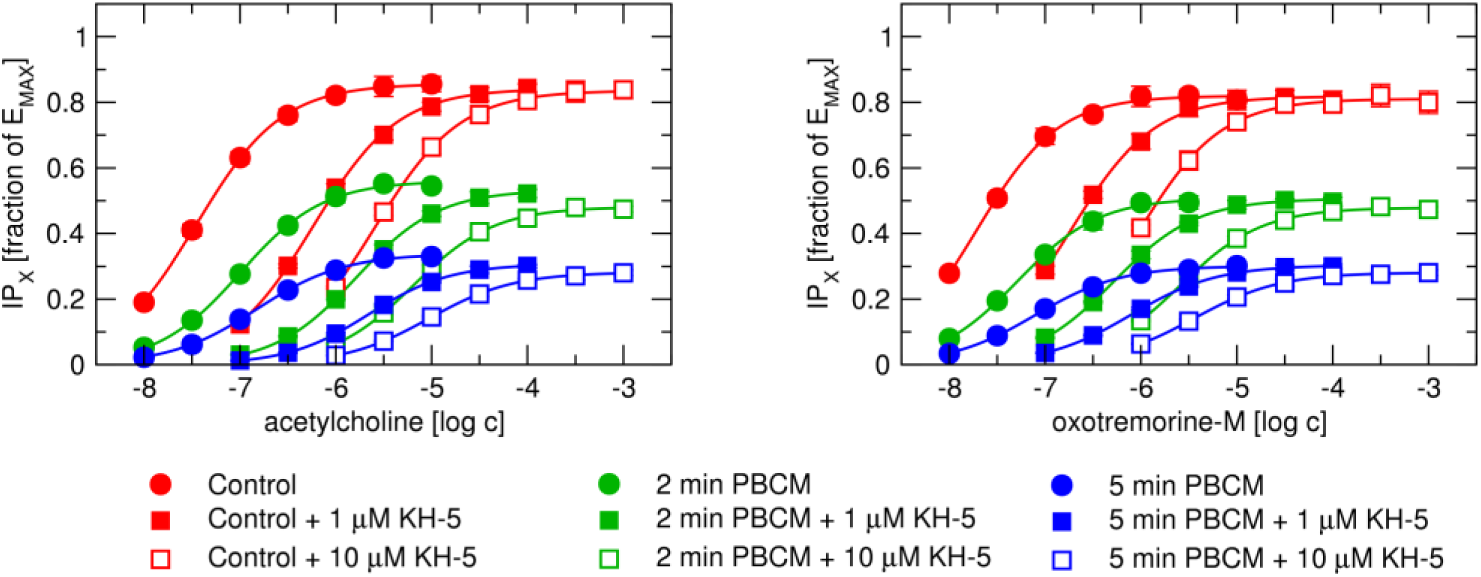
Effects of PBCM treatment on functional responses of M_1_ receptors to agonists. Functional response of M_1_ receptors to agonists acetylcholine (left) and oxotremorine-M (right) in the absence (circles) or presence of 1 μM (closed squares) or 10 μM (open squares) antagonist KH-5 at concentrations indicated in the legends was measured as accumulation of inositol phosphates (IP_X_) in control (red) and PBCM-treated (green and blue) CHO cells. Abscissa, concentration of agonist expressed as a decadic logarithm of molar concentration. Ordinate, level of IP_X_ expressed as a fraction of E_MAX_. Curves and their EC_50_ values were obtained by fitting Equation 9 to the data. Data are means ± SD from 5 independent experiments.

## Binding experiments

To investigate the interaction mechanism between an agonist and the compound KH-5, we conducted competition binding experiments involving KH-5 and the radiolabeled ligand [^3^H]NMS, in the presence of a fixed concentration of various agonists, as illustrated in Figure 5. Initially, we determined the dissociation constant (K_D_) of [^3^H]NMS through saturation binding experiments, fitting Equation 1 to the data, with detailed results provided in Supplementary Information Table S1. Subsequently, we calculated the inhibition constants (K_I_) of KH-5 from binding assays performed without any agonist, fitting Equation 2 to the data, as shown in Figure 5 (black data points Additionally, we obtained the inhibition constants (K_I1_ and K_I2_) for the tested agonists—namely acetylcholine, oxotremorine-M, and xanomeline—by fitting Equation 3 to the competition data, with results summarized in Supplementary Information Table S2. Then, the agonists were used at concentrations that reduced the control binding of [^3^H]NMS by approximately 25% to 40% (Figure 5). We applied two models: Equation 6, which describes the interaction among the labelled orthosteric tracer (NMS), the orthosteric ligand (agonist), and the potential allosteric modulator (KH-5); and Equation 7, which accounts for competition among three orthosteric ligands. Analysis showed that Equation 7 provided a better fit than Equation 6, indicating that the interaction between the tested agonists and KH-5 is likely competitive in nature.

**Figure 5.**
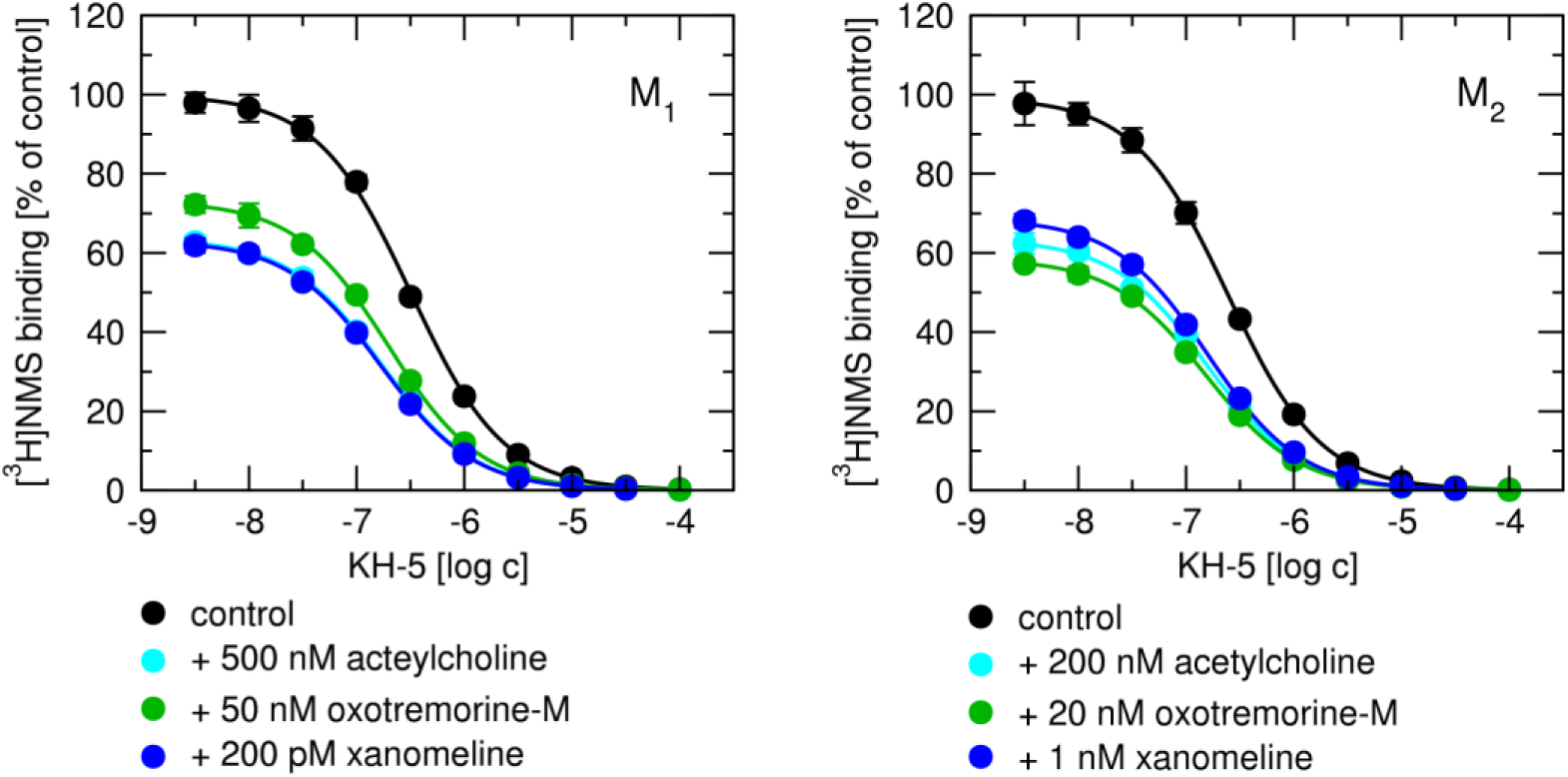
Competition of KH-5 with agonists for binding to M_1_ and M_2_ receptors. Effects of increasing concentrations of KH-5 on the binding of [^3^H]NMS in the absence (black) or presence of acetylcholine (cyan), oxotremorine-M (green) or xanomeline (blue) at fixed concentration indicated in the legends to membranes from CHO cells expressing M_1_ (left) and M_2_ (right) receptors. Abscissa, concentration of KH-5 is expressed as a decadic logarithm of molar concentration. Ordinate, specific [^3^H]NMS binding is expressed as a percentage of control in the absence of KH-5. Data are means ± SD from 5 independent experiments performed in quadruplicate.

To rule out the potential effects of NMS binding on the interaction between various agonists and the compound KH-5, we measured the direct competition between radiolabeled agonists, specifically [^3^H]acetylcholine and [^3^H]oxotremorine-M, and KH-5, as illustrated in Figure 6. We first determined the dissociation constants (K_D_) for these agonists through saturation binding experiments by fitting Equation 1 to the saturation binding data, with results summarised in Supplementary Information Table S1. At the M_1_ receptor, KH-5 at 3.16 to 31.6 nM concentration slightly increased binding of [^3^H] acetylcholine (by about 20 %) and at 10 nM concentration slightly increased binding of [^3^H]oxotremorine-M (by about 10 %) in each of five independent experiments. Binding curves from individual experiments are shown in Supplementary Information Figure S3. This transient increase was followed by complete inhibition at a 31.6 μM concentration of KH-5. In contrast to the M_1_ receptor, KH-5 only inhibited binding of tracers at the M_2_ receptor (Figure 6, right).

**Figure 6.**
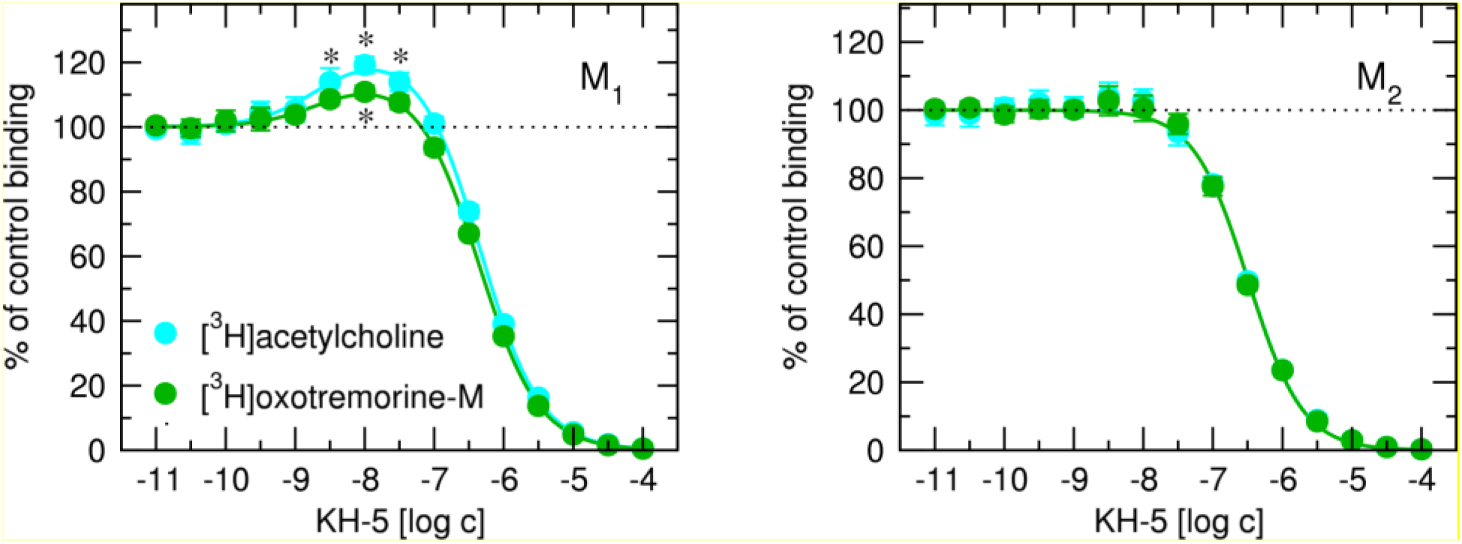
Interaction between KH-5 and tritiated agonists at M_1_ and M_2_ receptors. Effects of increasing concentrations of KH-5 on the binding of [^3^H]acetylcholine (cyan) and [^3^H]oxotremorine-M (green) to membranes from CHO cells expressing M_1_ (left) and M_2_ (right) receptors. Abscissa, concentration of KH-5 is expressed as a decadic logarithm of molar concentration. Ordinate, specific radioligand binding is expressed as a percentage of control in the absence of KH-5. Data are means ± SD from 5 independent experiments performed in quadruplicate. *, different from control (P < 0.05 according to one-sample t-test)

Subsequently, we applied different binding models to analyse the interactions. For competitive binding, we used Equation 2; for allosteric interactions, Equation 4; for interaction between labelled orthosteric ligand (agonist) and allosteric modulator (KH-5) binding to two allosteric sites, Equation 5; and for interaction between labelled orthosteric ligand (agonist) and dualsteric ligand (KH-5), Equation ^8^. Our analysis revealed that, although the increase in agonist binding at the M_1_ receptor was small, Equation 8 provided the best model for the M_1_-receptor binding data. (Equation 8 was the only one that passed the Runs-test.) This suggests a dualsteric binding mode (positive cooperativity from the allosteric site and competition for the orthosteric site) for KH-5 at the M_1_ receptor. The discrepancy between [^3^H]NMS data (Figure 5) versus [^3^H]acetylcholine and [^3^H]oxotremorine data may be explained by tracer bias and/or receptor activation state dependence. In contrast, based on AICc, Equation 2 provided the best model for the M_2_-receptor binding data. Meaning that M_2_ receptor data are best described by a purely orthosteric competitive binding model. However, an ectopic allosteric site with neutral cooperativity (α = 1) cannot be distinguished from orthosteric binding alone using current data.

From fitting Equation 8 to the M_1_-receptor binding data, acetylcholine pK_A_ was 8.57 ± 0.13, with an α value of 1.54 ± 0.07, and pK_B_ of 6.68 ± 0.03; and oxotremorine-M pK_A_ was 8.61 ± 0.25, with an α of 1.30 ± 0.03, and a pK_B_ of 6.64 ± 0.02. These values were based on five independent experiments. According to AICc, in each experiment, Equation 8 fitted better than Equation 4 orEquation 5. From fitting Equation 2 to the M_2_-receptor binding data, pK_I_ values for acetylcholine and oxotremorine-M were 6.71 ± 0.01 and 6.66 ± 0.01, respectively, also derived from five independent experiments.

Further, we aimed to narrow down the specific location of KH-5 binding within the receptor. To achieve this, we examined how KH-5 influenced the allosteric interaction between the orthosteric antagonist [^3^H]NMS and the classical allosteric modulator, alcuronium. We determined the equilibrium dissociation constant (K_D_) of [^3^H]NMS through saturation binding experiments, fitting Equation 1 to the saturation binding data (Supplementary Information Table S1). The inhibition constants (K_I_) of KH-5 were derived from competition binding experiments, where Equation 2 was fitted to the competition data (Supplementary Information Table S11).

Furthermore, we obtained the equilibrium dissociation constant of the allosteric modulator alcuronium (K_A_) and the binding cooperativity factor α from experiments arranged in a competition-like manner, fitting Equation 4 to the data (Table 2). To analyse the binding behaviour of the labelled orthosteric tracer ([^3^H]NMS) in the presence of multiple allosteric modulators, we applied Equation 5, which describes binding when two allosteric modulators compete for the same site, and Equation 6, which accounts for the presence of one competitive ligand and one allosteric modulator. These equations were fitted to data obtained with KH-5 at fixed concentrations, with K_D_, K_A_, and α held constant based on prior measurements (Figure 7). Our analysis revealed that Equation 6 provided a better fit than Equation 5 across all receptor subtypes, indicating that KH-5 competes with orthosteric tracer [^3^H]NMS and allosterically modulates binding of alcuronium.

**Table 2.** Interaction parameters between antagonist KH-5 and allosteric modulator alcuronium at individual subtypes of muscarinic receptors. Parameters of interaction between KH-5 and alcuronium were obtained by fitting Equation 6 to the data. The equilibrium dissociation constant of alcuronium (K_A_) is expressed as a negative decadic logarithm. α, factor of binding cooperativity between alcuronium and [^3^H]NMS. β, factor of binding cooperativity between alcuronium and KH-5. Data are means ± SD from 5 independent experiments performed in quadruplicate.

| | pK<sub>A</sub> | | | $\alpha$ | | | $\beta$ | | |
| --- | --- | --- | --- | --- | --- | --- | --- | --- | --- |
| M<sub>1</sub> | 5.15 | ± | 0.30 | 0.618 | ± | 0.073 | 0.165 | ± | 0.006 |
| M<sub>2</sub> | 5.96 | ± | 0.12 | 1.89 | ± | 0.04 | 10.9 | ± | 0.2 |
| M<sub>3</sub> | 4.68 | ± | 0.21 | 0.615 | ± | 0.035 | 0.508 | ± | 0.113 |
| M<sub>4</sub> | 5.18 | ± | 0.47 | 1.36 | ± | 0.14 | 0.830 | ± | 0.062 |
| M<sub>5</sub> | 4.64 | ± | 0.10 | 0.340 | ± | 0.035 | 0.308 | ± | 0.011 |

**Figure 7.**
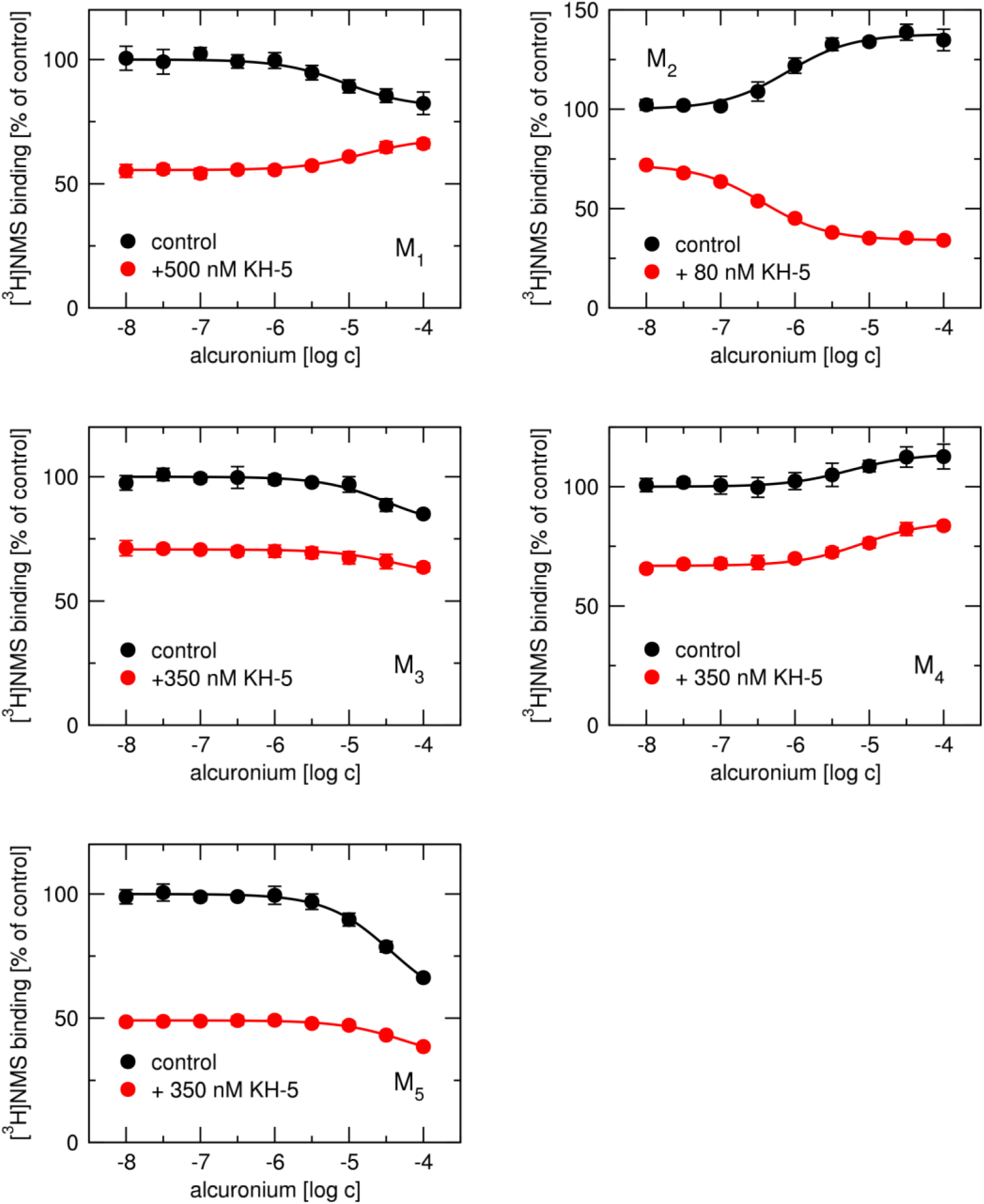
Interaction between antagonist KH-5 and allosteric modulator alcuronium at individual subtypes of muscarinic receptors. The mode of interaction between KH-5 and alcuronium was inferred from the effects of alcuronium on [^3^H]NMS binding in the presence versus absence of KH-5. Curves were obtained by fitting Equation 4 (absence of KH-5, black) or Equation 6 (presence of KH-5, red) to the data. The concentration used is indicated in the data labels. Abscissa, concentration of alcuronium expressed as decadic logarithm of molar concentration. Ordinate, [^3^H]NMS binding expressed as a percentage of [^3^H]NMS binding in the absence of alcuronium. Data are means ± SD from 5 independent experiments performed in quadruplicate.

We observed that KH-5 exhibited strong positive cooperativity—more than tenfold—with alcuronium at the M_2_ receptor subtype. Conversely, at other subtypes, KH-5 demonstrated negative cooperativity with alcuronium, with the degree of interaction varying. Specifically, at the M_4_ receptor, the affinity decreased by approximately 20%, whereas at the M_1_ receptor, the decrease was as much as sixfold. These results exclude KH-5 binding to the same allosteric site as alcuronium.

## Molecular modelling

Based on our analysis of experimental data, we identified an allosteric component in the binding of KH-5 to muscarinic receptors. To explore this further, we modelled the potential binding of KH-5 to the allosteric site located in the extracellular vestibule, adjacent to the orthosteric binding site. Our initial step involved docking KH-5 to the ectopic binding site, which was derived from the known ectopic/allosteric binding of xanomeline observed in the cryo-EM structure of the M_4_ receptor (PDB ID: 8FX5). We performed docking simulations across five receptor subtypes in their inactive conformations, specifically M_1_ (PDB ID: 5CXV), M_2_ (3UON), M_3_ (4DAJ), M_4_ (5DSG), and M_5_ (6OL9). Additionally, we included three subtypes with available active conformations: M_1_ (6OIJ), M_2_ (4MQT), and M_4_ (8FX5). Our estimates of KH-5 binding energy to the ectopic sites in inactive receptor conformations ranged from 5.3 kcal/mol at M_1_ to 7 kcal/mol at M_3_. These energies corresponded to dissociation constants between 130 μM and 4 μM (Supplementary Information Table S8), indicating possible binding. When compared to the binding energy at the orthosteric site, which ranged from 6 kcal/mol at M_2_ to 7.8 kcal/mol at M_3_, the difference was less than 1 kcal/mol, indicating very similar affinity for the ectopic and orthosteric sites and concurrent population of both sites in experiments. Notably, the affinity of KH-5 for the ectopic site in active conformations was higher for M_1_, similar for M_2_, and lower for M_4_ compared to inactive conformations. Altogether, docking results support the possibility that KH-5 specifically binds to the ecopic/allosteric site.

Generally, we observed that KH-5 bound more deeply within the binding pocket of inactive receptor conformations than in active ones. For instance, in the case of the M_1_ receptor (Figure 8) in an inactive state, the top binding pose of KH-5 was positioned adjacent to the orthosteric site, forming a hydrogen bond with Y381 (Y^6.51^ according to Ballesteros-Weinstein numbering (Ballesteros and Weinstein, 1995)). This pose did not interact with Y179 (Y^45.51^) in the second extracellular loop (ECL2) nor with K^6.62^ at the extracellular edge of transmembrane helix 6 (TM6). Conversely, in the active conformation, KH-5 did not interact with Y^6.51^ but instead engaged in hydrophobic interactions with Y^45.51^ and ionic interactions with K^6.62^ at the ectopic binding site. No substantial differences between stereoisomers were detected. These findings suggest that the binding mode of KH-5 varies depending on the receptor’s conformational state, which could influence its pharmacological profile and functional effects, as well as the observed binding affinity and/or binding mode may be dependent on receptor activation state.

**Figure 8.**
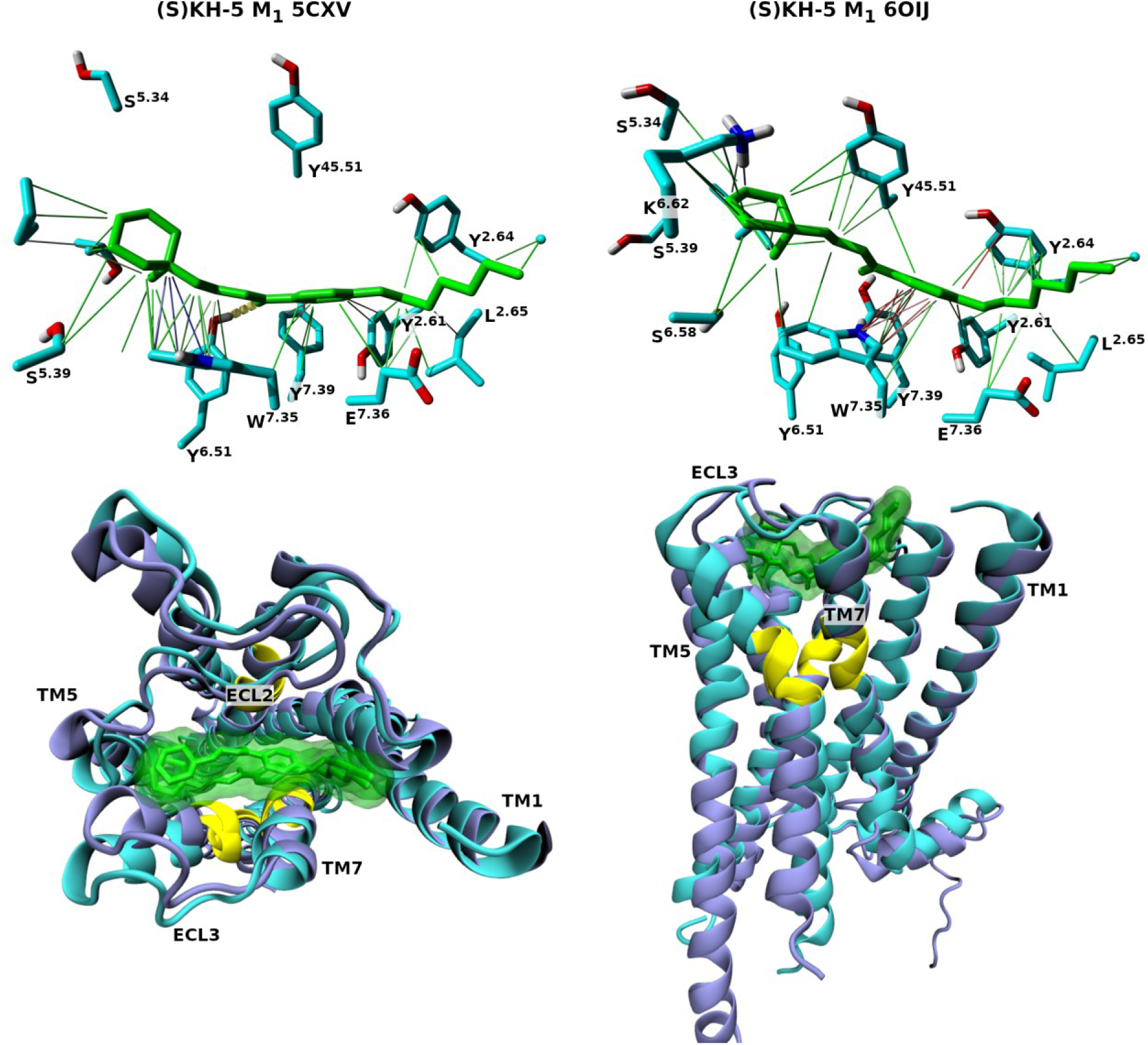
Interactions of KH-5 in top binding poses at the M_1_ receptor. **Top**, binding of (S)KH-5 (green) stereoisomer to the ectopic binding site of the M_1_ receptor in an inactive (left) and active(right) conformation. Atoms: cyan – carbon; blue – nitrogen; red – oxygen; white – hydrogen. Interactions: dashed yellow – hydrogen bonds; purple – ionic; green – hydrophobic; red – π-π; blue or white – cation-π. **Bottom**, location of (S)KH-5 binding to the ectopic site of the M_1_ receptor in an inactive (cyan) and an active (ice blue) conformation. Yellow – the orthosteric binding site.

To investigate the stability of modelled KH-5 binding to muscarinic receptors and identify key amino acid residues involved in binding at the ectopic site, we conducted three independent 120-ns molecular dynamics (MD) simulations. We analysed the trajectories obtained from these simulations to assess the stability and interactions of KH-5. With two exceptions (1 replica of (S)KH-5 at M3 4DAJ and 1 replica of (R)KH-5 at M4 8FX5), KH-5 equilibrated in the ectopic site within the first 20 ns, and KH-5 remained bound to the ectopic site throughout the MD simulations (Supplementary Information Figure S4). In these two exceptions, KH-5 remained close to the initial pose (RMSD < 2 Å) for the initial 30 ns and then began to translocate. The RMSD values of the final 100 ns of MD ranged from 3.05 to 6.96 Å (Supplementary Information Table S9). While the RMSD of 7 Å may seem large, it is smaller than the ectopic/allosteric site that has a 22 Å diameter in the plane of the membrane and is 10 Å in height perpendicularly to the membrane. Thus, KH-5 exhibits significant mobility within the ectopic site, consistent with transient interactions rather than a fixed binding pose, as supported by the consistency of interacting residues in the site and frequencies of interaction (Figure 9). Although the hydrogen bond interaction with Y^6.51^, observed at the M_1_ top pose, was lost during the MD simulations, the analysis using the Simulation Interaction Diagram tool revealed that the principal residues involved in binding were consistently F/Y^2.61^, Y^2.64^, Y^45.51^, W^7.35^, and Y^7.39^. These residues participated in interactions at both inactive and active conformations of the receptor (Figure 9). Also, no substantial differences between stereoisomers were observed. For the sequence-conservation/position table for the ectopic site, see Supplementary Information Table S10.

**Figure 9.**
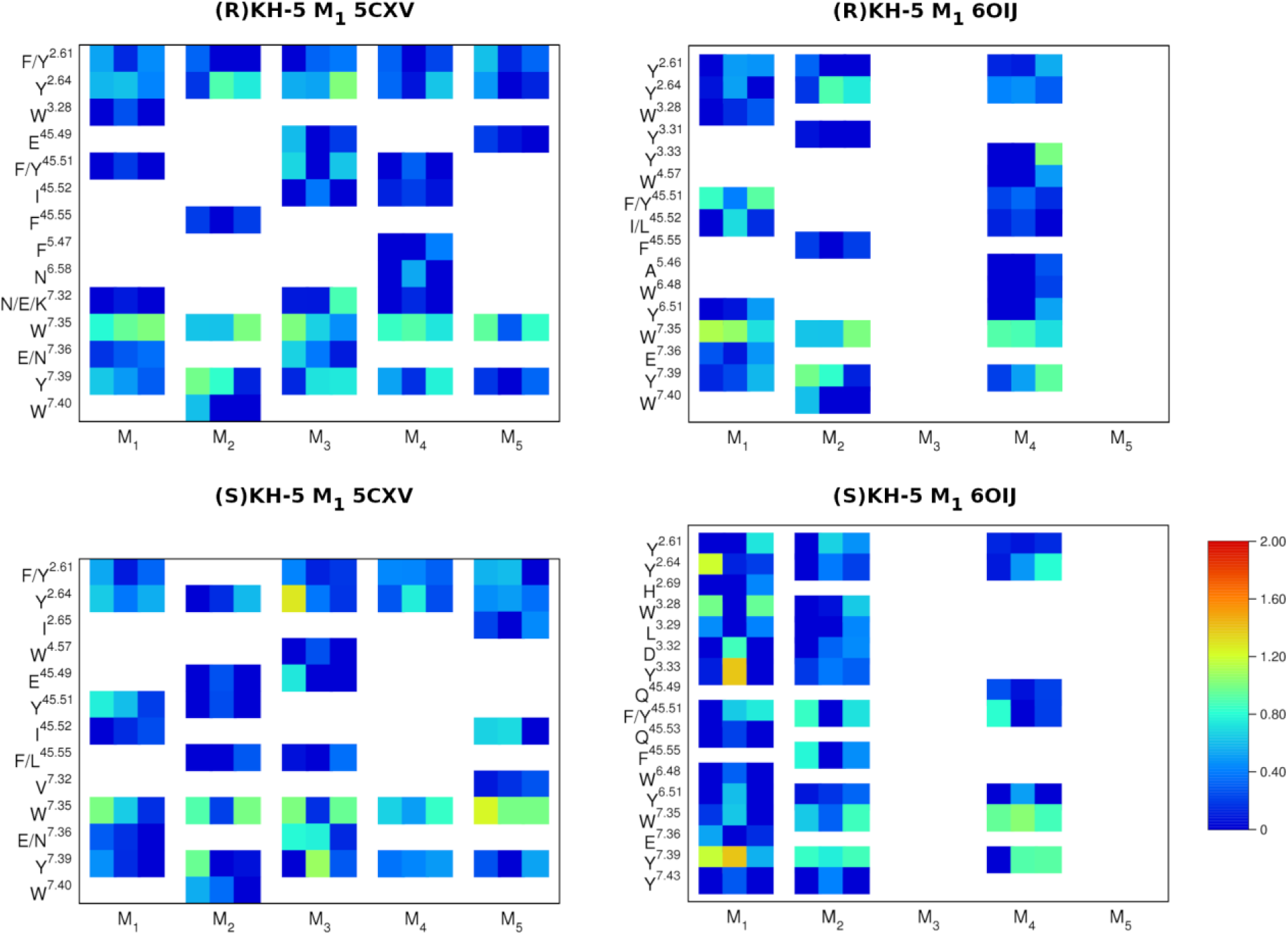
Interactions of KH-5 stereoisomers in the ectopic sites of muscarinic receptors. Histograms of interactions between KH-5 stereoisomers and receptors indicated on the x-axis, with amino acid residues in the ectopic site indicated on the y-axis, were calculated from MD trajectories using Ligand Interaction Diagram in Maestro. Only residues interacting with KH-5 at more than 30 % of time frames in at least one MD replica are displayed. Each square represents one of 3 independent MD replicas. Values over 1.0 are possible as some protein residues may make multiple contacts with KH-5 within one timeframe. However, no values greater than 1 were obtained.

## Mutant receptors

To test predictions of molecular modelling, amino acids Y82 (Y^2.61^), Y85 (Y^2.64^), Y179 (Y^45.51^), W400 (W^7.35^) and Y404 (Y^7.39^) of the M_1_ receptor were individually mutated to alanine and transiently expressed in CHO cells. Mutation Y404 exerted the greatest decrease, 24 times, in affinity for NMS (Supplementary Information Table S12). Mutations Y82A, Y85A, Y179A and W400A decreased binding affinity for NMS by only about two-fold. The effect of these mutations on affinity for acetylcholine was, in general, smaller. Mutations Y82A, Y85A and Y179A had no or negligible effect on binding affinity for acetylcholine. Mutation W400A decreased the affinity for acetylcholine 2.5 times, and mutation Y404 decreased the affinity for acetylcholine 3.9 times. Expression level highly varied among transfections (Supplementary Information Table S12). However, the expression level did not affect the binding affinities of traces or KH-5.

In contrast to NMS and acetylcholine, mutation Y404A did not affect binding affinity for KH-5, but mutation Y82A decreased it 4 times (Supplementary Information Table S13). At the wildtype M_1_ receptor, KH-5 at concentrations from 3 to 30 nM increased binding of acetylcholine up to 120 % of acetylcholine binding in the absence of KH-5 (Figure 10). Fitting the dualsteric binding model (Equation ^8^) to the data resulted in estimates of 3.5 nM binding affinity of KH-5 for the allosteric binding site (K_A_) and a factor of binding cooperativity between KH-5 and acetylcholine, α = 1.69 (Supplementary Information Table S13). This transient increase was followed by a decrease and complete inhibition at 30 μM concentration of KH-5, with an estimated 130 nM affinity of KH-5 for the orthosteric binding site (K_B_). Mutations Y82A. Y85A, Y179A and W400A abolished this transient increase in acetylcholine binding and decreased KH-5 affinity for the orthosteric site from two-fold (Y85A, W400A) to four-fold (Y82A). In contrast, mutation W404A exaggerated this transient increase, increasing acetylcholine binding at 3 to 100 nM concentration of KH-5 up to 130 % of control. Fitting Equation 8 gave estimates of K_A_ of 3 nM and α of 2.36. Mutation W404A did not affect KH-5 affinity for the orthosteric site (K_B_). These results suggest dualsteric binding (positive cooperativity from the allosteric site and competition for the orthosteric site) of KH-5 at wild-type and Y404A mutant M_1_ receptor. Contrast between the lack of effect of the Y404A mutation on the affinity of KH-5 and a strong decrease in affinity of orthosteric tracers NMS and acetylcholine indicates a distinct binding geometry to the orthosteric site. Mutations in the allosteric site affect KH-5 binding, further supporting an allosteric component of the dualsteric mode of binding of KH-5.

**Figure 10.**
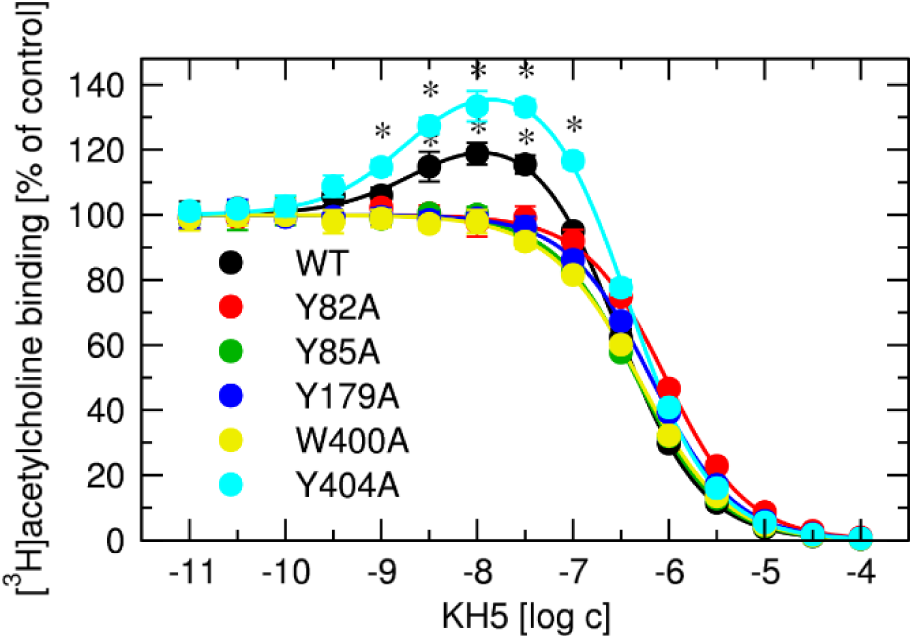
Interaction between KH-5 and tritiated acetylcholine at mutant M_1_ receptors. Effects of increasing concentrations of KH-5 on the binding of [^3^H]acetylcholine to membranes from CHO cells expressing M_1_ wildtype (WT) (black) and mutant receptors (indicated in the legend). Abscissa, concentration of KH-5 is expressed as a decadic logarithm of molar concentration. Ordinate, specific [^3^H]acetylcholine binding is expressed as a percentage of control in the absence of KH-5. Data are means ± SD from 5 independent experiments performed in quadruplicate. *, different from control (P < 0.05 according to one-sample t-test)

## Operational model

The presented data indicate that KH-5 interacts with both the orthosteric and allosteric binding sites of the receptor. Evidence from molecular modelling and mutagenesis studies supports the presence of an allosteric component in this interaction. To analyse these interactions quantitatively, we derived the operational model of dualsterically modulated agonism (OMDMA) (Supplementary Information), which we used to fit the functional-response curves in the presence of KH-5. First, we determined the maximal response of the system (E_MAX_) as described previously (Jakubík et al., 2019b). Then, we converted the magnitude of the functional responses of the M_1_ receptor to various agonists into the fraction of E_MAX_ (Supplementary Information Figure S5). The control curves, obtained in the absence of KH-5, were fitted using Equation 10 to determine key parameters, the equilibrium dissociation constant of the agonist (K_A_) and its operational efficacy (τ). Following this, a global three-dimensional fit was performed using the OMDMA (Equation 13). In this model, the concentrations of the agonist (x) and KH-5 (y) served as independent variables, while the functional response (z) was the dependent variable. During this process, the previously determined values of K_A_ and τ were held constant. Fitted parameters were constrained within specific ranges: equilibrium dissociation constants K_B_ and K_C_ from 1 nM to 1 mM, and cooperativity factors α, β and δ from 0.01 to 100. The fitting process successfully converged for most data sets, except for xanomeline, where the cooperativity factors reached the limits of the constraints.

Equation 13, which describes dualsteric inhibition, fitted the functional response data better than Equation 11, which describes competitive inhibition (Supplementary Information Figure S6). Specifically, Equation 11 failed to capture the saturation of functional antagonism observed at micromolar concentrations of KH-5. The AICc score for Equation 13 was better than that for Equation 11, except for oxotremorine-M and xanomeline, for which the scores were identical. For oxotremorine-M, both equations failed the residual test. For xanomeline, Equation 11 failed the residual test, whereas Equation 13 passed.

Equation 13 also fitted all data better than Equation 12, which describes allosteric inhibition. Specifically, Equation 12 failed to fit the data obtained with 10 μM KH-5. The parameter estimates derived from the model are summarised in Supplementary Information Table S14. Notably, estimates obtained using Equation 13 had large error margins, indicating substantial uncertainty and suggesting interdependence among the parameters. This uncertainty limits the extent to which the modulatory effects of KH-5 on receptor activity can be quantified.

The analysis predicted that KH-5 has a higher affinity for the orthosteric site, in the nanomolar range, than for the allosteric site, in the micromolar range. In addition, the cooperativity factors were generally positive, supporting positive modulation. However, these estimates should be interpreted with caution because of the substantial uncertainty and constraints encountered during model fitting.

Then we performed this global three-dimensional fit of OMDMA (Equation 13) to the data in Figure 4 (Figure 11). Unlike in previous analysis, we fixed the factor of binding cooperativity between KH-5 and agonist (α) to the values determined in binding experiments with radiolabelled agonists (Figure 6). The parameter estimates derived from the model are summarised in Table 3. Notably, the error margins of estimates for Equation 13 are smaller than in Supplementary Table S14, indicating that fixing the α value decreases model uncertainty.

**Figure 11.**
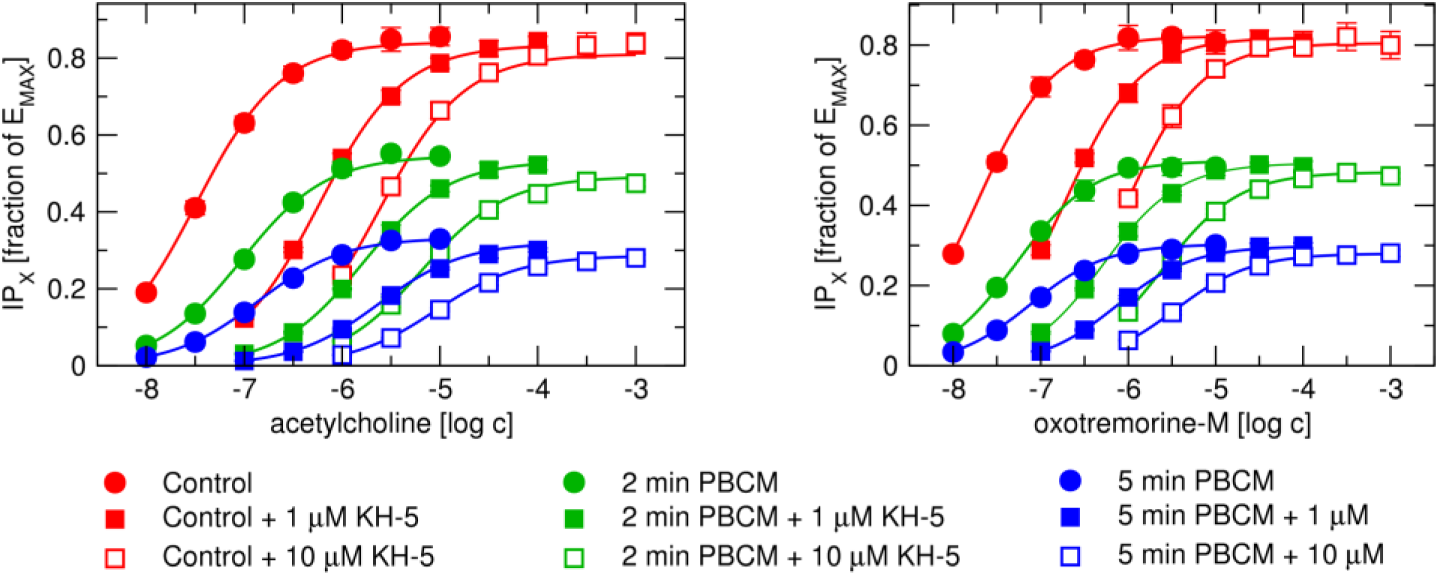
Fitting of the operational model of duasterically modulated agonism (OMDMA) to functional responses at M_1_ receptor. Data are pooled data of functional response of M_1_ receptors to agonists acetylcholine (left) and oxotremorine-M (right) in the absence (circles) or presence of 1 μM (closed squares) or 10 μM antagonist KH-5, measured as accumulation of inositol phosphates (IP_X_) in CHO cells shown in Figure 4, expressed as a fraction of system maximal response (E_MAX_). Curves are global 3D-fits of Equation 13, where the concentrations of an agonist and KH-5 are two independent variables. K_A_, τ and α were fixed to values predetermined in control and binding experiments. Data are means ± SD from five experiments performed in quadruplicate.

**Table 3.** Estimates of parameters ± SD of OMDMA fitted to data in Figure 11. Equilibrium dissociation constant of agonist (K_A_) and its operational efficacy (τ) were obtained by fitting Equation 10 to the functional response of control cells (τ_0_) and cells treated with PBCM for 2 (τ_1_) and 5 mins (τ_2_) in the absence of KH-5. The factor of binding cooperativity α was obtained by fitting Equation 8 to the binding data in Figure 6. Equilibrium dissociation constants of KH-5 for the allosteric (K_B_) and orthosteric (K_C_) binding sites, respectively, factor of binding cooperativity between two molecules of KH-5 bound to the allosteric and orthosteric binding sites (β) and cooperativity factor of operational efficacy (δ) were obtained by global fit of Equation 13 to all the data. Equilibrium dissociation constants are expressed as negative decadic logarithms. Cooperativity factors are expressed as decadic logarithms.

|  | acetylcholine | Oxotremorine-M |
| --- | --- | --- |
| pK<sub>A</sub> | 6.65 ± 0.03 | 6.97 ± 0.04 |
| $\tau_0$ | 4.91 ± 0.45 | 3.90 ± 0.32 |
| $\tau_1$ | 1.06 ± 0.04 | 0.863 ± 0.038 |
| $\tau_2$ | 0.422 ± 0.002 | 0.358 ± 0.006 |
| log($\alpha$) | 0.188 ± 0.019 | 0.114 ± 0.010 |
| pK<sub>B</sub> | 4.36 ± 2.53 | 4.06 ± 4.52 |
| pK<sub>C</sub> | 7.30 ± 0.16 | 7.00 ± 0.13 |
| log($\beta$) | 0.30 ± 2.60 | 2.00 ± 3.53 |
| log($\delta$) | 0.65 ± 2.53 | 1.83 ± 3.52 |

## Discussion

Our previous investigations revealed that the functional antagonism potency of KH-5 exceeds its binding affinity, suggesting a complex mechanism underlying its antagonistic action. We hypothesised that KH-5 operates as a dualsteric antagonist, exerting positive cooperativity from an allosteric site while also competing for the orthosteric site on the M_1_ muscarinic receptor. To test this hypothesis, we performed a comprehensive series of experiments, including assessments of KH-5 functional antagonism against various agonists, analysis of its binding interactions with both orthosteric and allosteric ligands across different experimental conditions, and evaluation of the effects of receptor mutations on KH-5 binding. We obtained mixed results. Some data supported characteristics typical of competitive antagonism, whereas other findings indicated allosteric mechanisms, implying that KH-5 may operate through multiple modes of action. Nonetheless, certain data points effectively exclude the possibility that KH-5 functions as a pure PAM antagonist. The most parsimonious model simulating such a complex behaviour is dualsteric dual-acting modulation (operational model of dualsterically modulated agonism – OMDMA), meaning KH-5 binding to both the orthosteric and allosteric sites and allosteric as well as competitive modulation of functional response.

Some of the data presented suggest that KH-5 functions as a dualsteric positive allosteric modulator (PAM) antagonist at the M_1_ receptor. Several key observations support this conclusion: First, the antagonistic potency of KH-5 varies depending on the agonist used, as demonstrated in Figure 2 andFigure 3 and summarised in Table 1. This probe dependency is characteristic of allosteric interactions, indicating that the effect of KH-5 depends on the specific ligand it interacts with. Second, its potency to antagonise functional responses to agonists is greater than its affinity (Table 1 vs Supplementary Information Table S11). This discrepancy suggests positive cooperativity in binding, where the presence of the agonist enhances the binding affinity of KH-5. Third, at nanomolar concentrations, KH-5 transiently increases the binding of radiolabelled acetylcholine and oxotremorine-M at the M_1_ receptor subtype, as depicted in Figure 6. This phenomenon aligns with Equation 8, which describes tracer binding in the context of a dualsteric ligand exhibiting both positive allosteric and competitive binding modes. However, pure allosteric antagonism is contradicted by the surmountable nature of KH-5 antagonism; KH-5 does not change the maximal response to agonist, E_MAX_ (Figure 2 andFigure 3).

For full agonists, a reduction in the observed maximal response (E’_MAX_) can, in principle, be masked by receptor reserve. In systems with high receptor reserve, substantial receptor occupancy or signalling capacity can be lost with little apparent effect on the maximal response. For example, in an operational model, a full agonist with high transducer efficacy (τ = 100) produces an E’_MAX_ of approximately 99% of the maximum system response (E_MAX_). Even if a full allosteric antagonist (β = 0) imposes 10-fold negative cooperativity on agonist binding (α = 0.1), the predicted maximal response decreases only modestly, to approximately 96.4% of E_MAX_. Such a small change would be difficult to resolve experimentally and, therefore, could be misinterpreted as an absence of antagonism.

However, this explanation is unlikely to account for the effects of KH-5. If the preservation of E’_MAX_ were simply due to receptor reserve, then reducing receptor availability should unmask a decrease in maximal response. This was not observed. Even in cells in which the receptor population was markedly reduced by chemical inactivation, KH-5 did not substantially decrease E’_MAX_. Importantly, this receptor depletion was functionally meaningful, as it produced pronounced reductions in both agonist potency and efficacy, confirming that receptor reserve had been substantially diminished under these conditions (Figure 4 and Supplementary Information Table S7). Thus, the persistence of maximal responses after receptor inactivation argues against receptor reserve as the primary explanation for the lack of an E’_MAX_ decrease.

Taken together, these findings indicate that KH-5 cannot be described as a simple allosteric antagonist whose effects are masked by spare receptors. Rather, its pharmacological profile is consistent with a more complex mode of action, in which antagonistic effects are counterbalanced, at least in part, by allosteric enhancement of receptor signalling efficacy.

To delineate the binding mode of KH-5, we performed complementary radioligand-binding experiments using orthosteric antagonist and agonist tracers. The results, again, presented evidence for both orthosteric and allosteric modes of interaction. In assays with the antagonist tracer [^3^H]NMS (Figure 5), the interaction between KH-5 and the tested agonists was best described by Equation 7, which assumes mutually exclusive binding of three orthosteric ligands. Under these conditions, KH-5 therefore behaved predominantly as an orthosteric competitor. A different profile was obtained with the agonist tracers [^3^H]acetylcholine and [^3^H]oxotremorine-M (Figure 6). In these assays, KH-5 produced a transient enhancement of agonist binding, and the data were best described by Equation 8. This model allows simultaneous occupancy of orthosteric and allosteric sites and is therefore consistent with a dualsteric interaction. The enhancement of agonist tracer binding suggests positive binding cooperativity between KH-5 bound at an allosteric or ectopic site and agonist occupancy of the orthosteric site, whereas the competitive profile observed with [^3^H]NMS reflects the orthosteric component of KH-5 binding. Such probe dependence is a recognised feature of GPCR allostery: the magnitude and even direction of an allosteric effect can change according to the orthosteric ligand used to probe the receptor (Valant et al., 2012; Wootten et al., 2013). It should be noted that, in these assays, radiolabelled agonists bind to high-affinity states that are sensitive to G-protein coupling and guanine nucleotides, whereas binding of muscarinic antagonists is unaffected by G-protein coupling or by the presence of guanine nucleotides (Flynn et al., 1989). Thus, these two assays involve different receptor activation states.

Together, these findings support a model in which KH-5 can engage the M_1_ receptor through more than one binding mode, with the apparent mechanism determined by both the tracer and the receptor conformational ensemble sampled in the assay (Borah et al., 2025; Elisi and Bottegoni, 2025; Ma et al., 2025). This interpretation is consistent with the broader view that GPCRs populate multiple active and inactive conformational states that can be differentially stabilised by orthosteric, allosteric, or bitopic ligands (Lane et al., 2017). It is also consistent with M_1_ muscarinic receptor studies showing that bitopic ligands can span the orthosteric site and extracellular allosteric/vestibular regions (Keov et al., 2014), thereby influencing ligand binding, receptor activation, and signalling bias.

Alcuronium functions as a classical allosteric modulator of muscarinic receptors, binding specifically between the second and third extracellular loops of the receptor (Krejcí and Tucek, 2001). In the binding assays, radiolabelled orthosteric tracer [^3^H]NMS was used in conjunction with alcuronium, an allosteric modulator, and investigated ligand KH-5 (Figure 7). The binding data obtained from these experiments were best described by Equation 6, which models the interactions among the orthosteric tracer, the competitive orthosteric ligand KH-5, and the allosteric ligand alcuronium. Notably, the data suggest that KH-5 binds rather to the orthosteric binding site than to the classical allosteric site located between the second and third extracellular loops of the receptor. Again, when [^3^H]NMS was used as a tracer, the detected binding mode of KH-5 is orthosteric.

Molecular modelling studies of KH-5 binding to the extracellular ectopic or allosteric site of the receptor (Figure 8) have provided insights into its possible interaction mechanisms in this site. Subsequent MD simulations demonstrated that KH-5 remains within this allosteric site (Supplementary Information Figure S4 and Table S9). Analysis of MD trajectories identified key amino acid residues involved in the interaction, notably phenylalanine or tyrosine at position 2.61, tyrosine at 2.64, tyrosine at 45.51, tryptophan at 7.35, and tyrosine at 7.39 (Figure 9). These residues appear to play a key role in KH-5 binding to the allosteric binding site.

Further investigations involving site-directed mutagenesis revealed that alterations at these principal residues significantly impacted KH-5 binding affinity and its interaction with acetylcholine. Mutations such as Y82A, Y85A, Y179A, and W400A, located in the extracellular allosteric region, abolished the positive cooperativity observed between acetylcholine and KH-5, confirming that the allosteric site is involved in ligand binding. Interestingly, mutation of Y404, situated at the edge of the orthosteric site within the so-called “tyrosine lid,” enhanced the positive cooperativity between KH-5 and acetylcholine. This may be due to the removal of a steric clash point that limits allosteric engagement in the wild type (Dror et al., 2013). Although this mutation reduced the affinity for orthosteric ligands like NMS and acetylcholine, it did not alter the affinity of KH-5 for the orthosteric site, suggesting a distinct binding geometry for KH-5 compared to these orthosteric ligands.

Mutations in the extracellular vestibule, such as Y82A, Y85A, and W400A, resulted in decreased affinity for both orthosteric ligands and KH-5, indicating that this region contributes significantly to the overall ligand affinity. These findings align with previous research (Jakubik et al., 2000; Bock et al., 2012; Kappel et al., 2015; Kistemaker et al., 2019), which emphasises the importance of the extracellular vestibule in ligand binding and receptor modulation. Overall, these results highlight the complex interplay between orthosteric and allosteric sites in receptor function and ligand interactions.

Taken together, pure allosteric antagonism is contradicted by the surmountable nature of KH-5 antagonism, but presented functional and binding data suggest the presence of an allosteric component that is further supported by molecular modelling and mutations in the ectopic/allosteric site. In principle, allosteric modulation is limited by a factor of binding cooperativity α that defines the maximal effect on binding of the second ligand. Once the agonist fully occupies its binding site, its effect reaches a ceiling because the allosteric antagonist does not compete directly for the same binding site, and the effect cannot be overcome by simple mass action. Therefore, increasing agonist concentration cannot displace the allosteric antagonist and restore full receptor activation. We derived Equation 13 of the OMDMA (Supplementary Information, Derivations) as the simplest model fitting the obtained data. According to Equation 13, unsurmountable antagonism also applies to dualsteric antagonists (Figure 12, upper graphs). Positive binding cooperativity between the dualsteric antagonist and agonist further exacerbates a decrease in E_MAX_ (Figure 12, top right vs. top left). If dualsteric antagonist exhibited positive cooperativity at the allosteric site (α>1) but did not antagonise the functional response from the allosteric site (δ=1) then for K_B_/α < K_C_, where K_B_ and K_C_ are equilibrium dissociation constants for allosteric and orthosteric binding sites, respectively, dualsteric antagonist would positively modulate functional response (Figure 12, middle left) and for K_B_/α ≥ K_C_ its antagonistic potency would equal to K_C_ (Figure 12, middle right). Therefore, observed potency greater than binding affinity cannot be the result of positive cooperativity between KH-5 and agonist. Rather, it is the result of positive cooperativity between two molecules of KH-5 bound to the orthosteric and allosteric sites (Figure 12, bottom). At the same time, the decrease in E_MAX_ must be compensated by the positive operational cooperativity of KH-5 from the allosteric site (δ > 1). Thus, KH-5 is not a pure dualsteric antagonist but probably a dualsteric dual-acting modulator. KH-5 is a dualsteric ligand, binding to both orthosteric and allosteric sites. At the same time, KH-5 is a dual-acting modulator, exerting positive operational cooperativity with agonist from the allosteric binding site and competitive antagonism from the orthosteric site. On top of it, two molecules of KH-5 bound to the allosteric and orthosteric sites, respectively, exert positive binding cooperativity.

**Figure 12.**
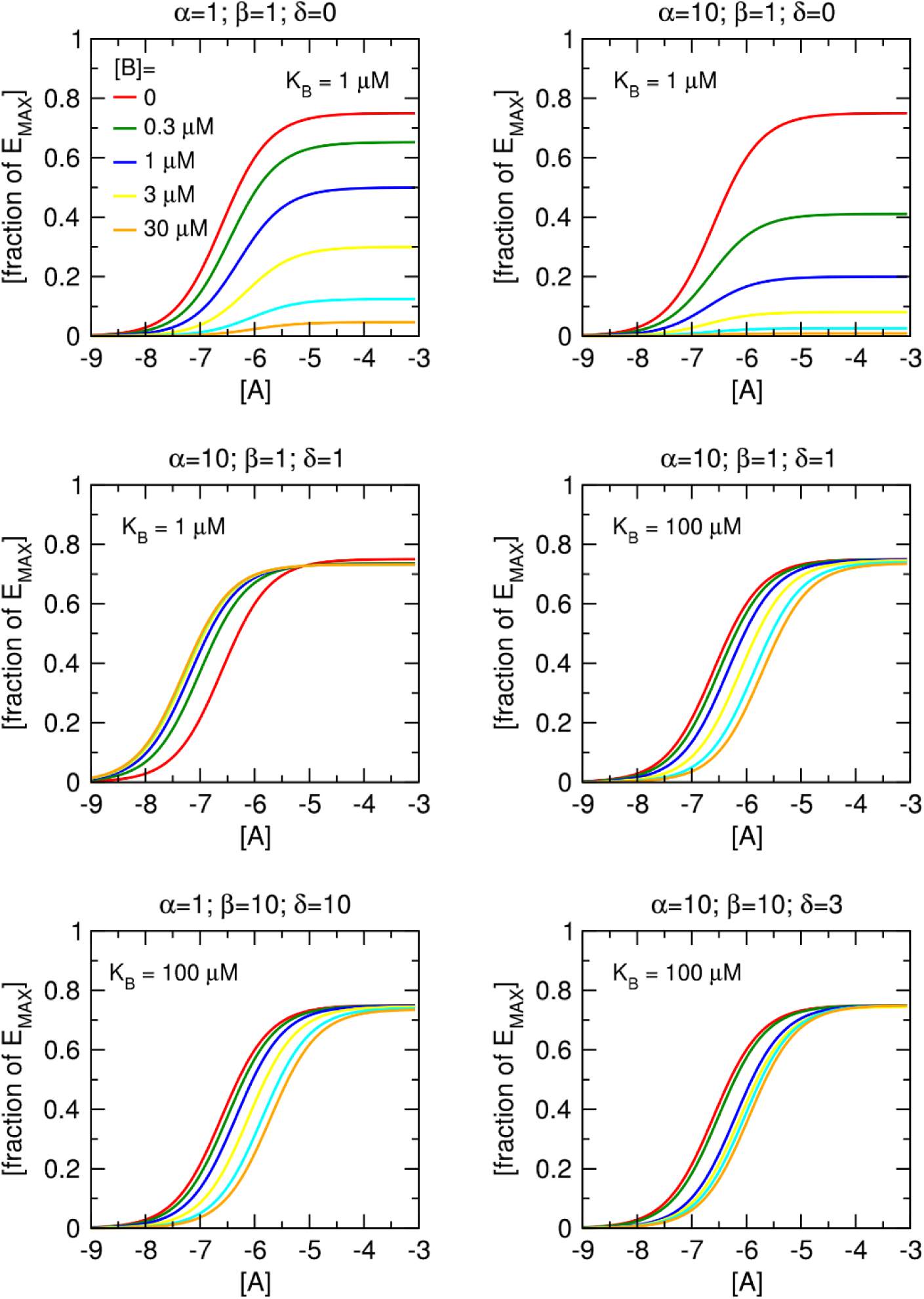
Theoretical functional responses to an agonist, A, in the presence of the dualsteric ligand B. Curves of functional responses were modelled according to Equation 13. The concentration of the dualsteric ligand is indicated in the data labels. Factors of cooperativity are shown in the graph headers. Equilibrium dissociation constants K_A_ and K_C_ were equal to 1 μM. K_B_ was equal to 1 or 100 μM as indicated in the legends. Abscissa, concentration of an agonist expressed as decadic logarithm of molar concentration. Ordinate, functional response expressed as a fraction of maximal response E_MAX_.

To pharmacologically model the mechanism of action of KH-5 at the M_1_ receptor, OMDMA Equation ^1^^3^ was fitted to functional response data (Supplementary Information Figure S5 and S6 and Table S14). Results indicate that, aside from xanomeline, the OMDMA Equation 13 effectively describes the functional-response curves observed in experimental settings. This suggests that the model is a plausible representation of the underlying biological processes involved in receptor modulation by KH-5. Furthermore, the theoretical analysis of OMDMA supports the empirical findings, especially regarding the estimated values of the three cooperativity factors: α, β, and δ. The fits are consistent with positive cooperativity/positive operational modulation, but the magnitudes of β and δ are not well identified and should not be interpreted quantitatively. Also, affinity for the allosteric site (K_B_) was predicted to be lower than affinity for the orthosteric site (K_C_). However, it is important to interpret these estimates with caution because of large uncertainties in parameter estimates. These uncertainties could stem from the inherent complexity of receptor modulation mechanisms and interdependent model parameters (Jakubík et al., 2020). Therefore, while the findings support the OMDMA, further studies with more refined experimental designs are necessary to confirm and quantify these results. Namely, because of parameter interdependence, it will be necessary to experimentally determine KH-5 parameters K_B_, K_C_ and β. As these parameters cannot be deduced directly from competition with [^3^H]NMS under equilibrium, sophisticated measurements of labelled KH-5 binding will need to be designed and executed. Labelling of KH-5 is potentially problematic. Radiolabelled KH-5 will likely exert high non-specific binding due to its lipophilicity. Fluorescent labelling will likely completely change the pharmacology of KH-5. Nevertheless, the analysis underscores the potential of OMDMA as a valid model of the mechanism of action of KH-5 at the M_1_ receptor.

Xanomeline is an atypical muscarinic receptor agonist characterised by sustained efficacy that remains resistant to washing (Christopoulos et al., 1998). Structurally, xanomeline shares a hexyloxy group with KH-5, a feature that contributes to their prolonged pharmacological effects (Jakubík et al., 2004; Randáková et al., 2018). The interaction between KH-5 and xanomeline in modulating receptor responses significantly differs from the mechanisms observed with classical antagonists (see Figure 2 andFigure 3). For instance, at the M_1_ receptor, KH-5 exhibits more than threefold greater potency in antagonising xanomeline than in antagonising acetylcholine. Additionally, Schild plot analyses reveal that while orthosteric agonists typically produce saturation in antagonism curves, xanomeline induces upward-curving responses, indicating positive operational cooperativity. In the end, OMDMA (Equation 13) does not fit the functional response data of xanomeline (Figure 11 and Table 3). Xanomeline is known to engage both orthosteric and allosteric binding sites simultaneously, suggesting a dual binding mode (Burger et al., 2023). The data imply that KH-5 likely shares this dual binding characteristic. Consequently, accurately modelling KH-5 antagonism of xanomeline-induced responses may require an even more complex framework than the OMDMA.

The M_1_ functional data were obtained using IP_X_ accumulation. Therefore, pathway/assay-dependent components cannot be completely excluded. However, KH-5 exerted greater potency to antagonise [^35^S]GTPγS binding (pKB) than binding affinity (pK_I_) in our previous study (Randáková et al., 2018). Moreover, the binding, receptor-depletion, modelling and mutagenesis data provide convergent evidence for a binding-level allosteric component.

An additional limitation is that the present equilibrium binding and functional data cannot formally exclude contributions from receptor oligomerisation (Marsango et al., 2018). Apparent cooperative interactions could, in principle, arise from inter-protomer effects in M_1_ receptor oligomers (Jakubík and Randáková, 2022). The mutagenesis and structural modelling data are consistent with an intra-protomer extracellular vestibule interaction, but they do not distinguish this mechanism from oligomer-dependent cooperativity.

In summary, the presented data are a mix of evidence for allosteric interaction as well as for competition. At M_1_ receptors, the most parsimonious model among those tested combines orthosteric competition with an ectopic/allosteric component. From the allosteric site, it positively modulates functional responses to agonists. From the orthosteric site, it exerts competitive antagonism of functional responses. Additionally, molecules of KH-5 bound to allosteric and orthosteric sites exert positive cooperativity. At M_2_ receptors, KH-5 behaves predominantly as an orthosteric antagonist under the present conditions, although a weak or probe-specific allosteric component cannot be excluded.

## Nomenclature of Targets and Ligands

Key protein targets and ligands in this article are hyperlinked to corresponding entries in http://guidetopharmacology.org, and are permanently archived in Concise Guide to Pharmacology 2021/22(151-157) (Alexander et al., 2021).

## Declaration of Transparency and Scientific Rigour

The authors declare that this study was designed, analysed and reported in accordance with the principles of transparent reporting and scientific rigour recommended by the British Journal of Pharmacology, including the relevant BJP guidance on experimental design and analysis in pharmacology. Experiments were performed as independent biological replicates with quadruplicate determinations, all data were included in the analyses, and statistical tests and model-comparison procedures are described in the Methods. Experimenters were blinded to the chemical structures of the tested compounds where applicable. Cell lines are identified using RRIDs, and targets and ligands are named according to BJP/IUPHAR nomenclature. This study did not involve animal experiments, human participants, immunoblotting, immunohistochemistry, or natural-product research; therefore, the corresponding BJP-specific reporting checklists were not applicable.

## Acknowledgements

The work was supported by the Czech Academy of Sciences institutional support [RVO:67985823] and the Grant Agency of the Czech Republic grant [23-04670S].

## Author contributions

All authors contributed to, read and approved the manuscript. AJR designed, supervised and conducted cell culturing, ligand binding and functional experiments, constructed and expressed mutants and performed raw data analysis. ED and NC performed cell culturing, ligand binding, functional experiments and raw data analysis. JJ performed molecular modelling, derivation of model equations and final data analysis. JJ conceived the project and its experimental design, wrote and finalised the manuscript and secured project financing.

## Data availability

The data that support the findings of this study are available from the corresponding author upon reasonable request.

## Abbreviations

CHO: cells Chinese hamster ovary cells
DMEM: Dulbecco modified Eagle’s medium
KHB: Krebs HEPES buffer
NMS: N-methyl scopolamine
PBCM: propylbezilyl choline mustard
TCA: trichloroacetic acid

