## Supplementary Information for "Dualsteric and dual-acting modulation of muscarinic receptors by antagonist KH-5"

### Supplementary Figures and Tables

***Table S1 Binding parameters of radioligands to membranes of CHO cells expressing individual subtypes of muscarinic receptors.***

*Equilibrium dissociation constants ($K_D$) of tritiated N-methylscopolamine (NMS), acetylcholine and oxotremorine-M were obtained by fitting Equation 1 to the data from saturation binding experiments and are expressed as negative decadic logarithms. Maximum binding capacity is expressed as pmol of binding sites per mg of membrane proteins. Data are means ± SD from 5 independent experiments performed in quadruplicates.*

| | [$^3$H]NMS | | | | | | [$^3$H]acetylcholine | | | | | | [$^3$H]oxotremorine-M | | | | | |
|---|---|---|---|---|---|---|---|---|---|---|---|---|---|---|---|---|---|---|
| | $pK_D$ | | | $B_{MAX}$ [pmol/mg prot.] | | | $pK_D$ | | | $B_{MAX}$ [pmol/mg prot.] | | | $pK_D$ | | | $B_{MAX}$ [pmol/mg prot.] | | |
| $M_1$ | 9.63 | ± | 0.03 | 1.76 | ± | 0.05 | 7.18 | ± | 0.03 | 0.342 | ± | 0.013 | 7.56 | ± | 0.03 | 0.364 | ± | 0.009 |
| $M_2$ | 9.43 | ± | 0.02 | 1.31 | ± | 0.03 | 7.22 | ± | 0.02 | 0.574 | ± | 0.018 | 7.60 | ± | 0.03 | 0.452 | ± | 0.013 |
| $M_3$ | 9.64 | ± | 0.04 | 1.69 | ± | 0.05 | 7.26 | ± | 0.02 | 0.370 | ± | 0.007 | 7.67 | ± | 0.03 | 0.292 | ± | 0.008 |
| $M_4$ | 9.67 | ± | 0.01 | 1.61 | ± | 0.03 | 7.28 | ± | 0.03 | 0.664 | ± | 0.021 | 7.69 | ± | 0.04 | 0.488 | ± | 0.018 |
| $M_5$ | 9.50 | ± | 0.03 | 0.914 | ± | 0.021 | 7.27 | ± | 0.03 | 0.194 | ± | 0.005 | 7.68 | ± | 0.03 | 0.158 | ± | 0.004 |

***Table S2 Binding parameters of muscarinic agonists to membranes of CHO cells expressing individual subtypes of muscarinic receptors.***

*Inhibition constants ($K_I$) of high and low affinity binding and fraction of low affinity sites were obtained by fitting Equation 3 to the data from competition experiments with [$^3$H]NMS. Constants are expressed as negative decadic logarithms. The fraction of low-affinity sites is expressed in percentages. Data are means ± SD from 5 independent experiments performed in quadruplicates.*

| | $pK_{i1}$ | | | $pK_{i2}$ | | | $F_2$ [%] | | |
|---|---|---|---|---|---|---|---|---|---|
| | | | | acetylcholine | | | | | |
| $M_1$ | 6.56 | ± | 0.12 | 4.09 | ± | 0.02 | 81 | ± | 3 |
| $M_2$ | 6.83 | ± | 0.11 | 4.13 | ± | 0.05 | 56 | ± | 3 |
| $M_3$ | 6.96 | ± | 0.11 | 4.31 | ± | 0.04 | 78 | ± | 3 |
| $M_4$ | 6.79 | ± | 0.10 | 4.23 | ± | 0.04 | 59 | ± | 3 |
| $M_5$ | 6.67 | ± | 0.15 | 4.12 | ± | 0.04 | 79 | ± | 3 |
| | | | | iperoxo | | | | | |
| $M_1$ | 9.60 | ± | 0.17 | 6.66 | ± | 0.03 | 81 | ± | 3 |
| $M_2$ | 9.53 | ± | 0.09 | 6.35 | ± | 0.03 | 51 | ± | 3 |
| $M_3$ | 9.72 | ± | 0.16 | 6.63 | ± | 0.03 | 81 | ± | 3 |
| $M_4$ | 9.77 | ± | 0.06 | 6.92 | ± | 0.04 | 57 | ± | 2 |
| $M_5$ | 9.51 | ± | 0.15 | 6.33 | ± | 0.02 | 79 | ± | 3 |
| | | | | oxotremorine-M | | | | | |
| $M_1$ | 7.36 | ± | 0.27 | 6.01 | ± | 0.05 | 80 | ± | 6 |
| $M_2$ | 7.92 | ± | 0.09 | 6.23 | ± | 0.03 | 66 | ± | 3 |
| $M_3$ | 8.13 | ± | 0.26 | 6.05 | ± | 0.04 | 83 | ± | 4 |
| $M_4$ | 8.34 | ± | 0.10 | 6.70 | ± | 0.02 | 69 | ± | 3 |
| $M_5$ | 7.93 | ± | 0.09 | 6.23 | ± | 0.04 | 83 | ± | 3 |
| | | | | pilocarpine | | | | | |
| $M_1$ | 7.27 | ± | 0.18 | 5.82 | ± | 0.04 | 83 | ± | 5 |
| $M_2$ | 7.20 | ± | 0.19 | 5.66 | ± | 0.11 | 59 | ± | 8 |
| $M_3$ | 7.29 | ± | 0.33 | 5.83 | ± | 0.03 | 83 | ± | 4 |
| $M_4$ | 7.54 | ± | 0.22 | 5.94 | ± | 0.07 | 64 | ± | 7 |
| $M_5$ | 6.85 | ± | 0.53 | 5.59 | ± | 0.08 | 73 | ± | 12 |
| | | | | xanomeline | | | | | |
| $M_1$ | 10.33 | ± | 0.19 | 7.20 | ± | 0.03 | 83 | ± | 3 |
| $M_2$ | 9.54 | ± | 0.08 | 7.47 | ± | 0.04 | 59 | ± | 3 |

| | $pK_{i1}$ | | | $pK_{i2}$ | | | $F_2$ [%] | | |
|---|---|---|---|---|---|---|---|---|---|
| $M_3$ | 9.67 | ± | 0.23 | 7.48 | ± | 0.03 | 81 | ± | 3 |
| $M_4$ | 10.34 | ± | 0.16 | 7.40 | ± | 0.02 | 64 | ± | 3 |
| $M_5$ | 9.28 | ± | 0.17 | 7.07 | ± | 0.03 | 81 | ± | 3 |

***Table S3 Parameters of functional responses of $M_1$ muscarinic receptors to agonists in the presence of KH-5.***

*Half-efficient concentrations ($EC_{50}$), observed maximal responses to agonists ($E'_{MAX}$) and Hill coefficients of functional responses of $M_1$ receptors to agonists were obtained by fitting Equation 9 to the experimental data. The $EC_{50}$ values are expressed as the decadic logarithm of molar concentration. Data are means ± SD from five independent experiments.*

| log[KH-5] | $logEC_{50}$ | | | $E'_{MAX}$ | | | $n_H$ | | |
|---|---|---|---|---|---|---|---|---|---|
| acetylcholine | | | | | | | | | |
| -- | -7.29 | ± | 0.04 | 3.70 | ± | 0.10 | 1.41 | ± | 0.14 |
| -7 | -6.71 | ± | 0.05 | 3.61 | ± | 0.13 | 1.54 | ± | 0.20 |
| -6.5 | -6.35 | ± | 0.05 | 3.64 | ± | 0.17 | 1.48 | ± | 0.24 |
| -6 | -6.01 | ± | 0.03 | 3.61 | ± | 0.09 | 1.48 | ± | 0.08 |
| -5.5 | -5.71 | ± | 0.03 | 3.61 | ± | 0.04 | 1.53 | ± | 0.06 |
| -5 | -5.39 | ± | 0.05 | 3.68 | ± | 0.06 | 1.52 | ± | 0.17 |
| iperoxo | | | | | | | | | |
| -- | -9.38 | ± | 0.06 | 4.55 | ± | 0.12 | 1.36 | ± | 0.17 |
| -7 | -9.00 | ± | 0.03 | 4.40 | ± | 0.05 | 1.50 | ± | 0.17 |
| -6.5 | -8.64 | ± | 0.02 | 4.44 | ± | 0.08 | 1.45 | ± | 0.09 |
| -6 | -8.20 | ± | 0.06 | 4.46 | ± | 0.14 | 1.42 | ± | 0.11 |
| -5.5 | -7.90 | ± | 0.05 | 4.44 | ± | 0.10 | 1.34 | ± | 0.15 |
| -5 | -7.57 | ± | 0.03 | 4.49 | ± | 0.09 | 1.39 | ± | 0.04 |
| oxotremorine-M | | | | | | | | | |
| -- | -7.52 | ± | 0.04 | 3.58 | ± | 0.07 | 1.60 | ± | 0.14 |
| -7 | -7.10 | ± | 0.03 | 3.61 | ± | 0.04 | 1.69 | ± | 0.09 |
| -6.5 | -6.79 | ± | 0.03 | 3.56 | ± | 0.08 | 1.70 | ± | 0.23 |
| -6 | -6.44 | ± | 0.04 | 3.56 | ± | 0.06 | 1.74 | ± | 0.17 |
| -5.5 | -6.05 | ± | 0.01 | 3.53 | ± | 0.05 | 1.79 | ± | 0.09 |
| -5 | -5.72 | ± | 0.02 | 3.56 | ± | 0.10 | 1.77 | ± | 0.18 |
| pilocarpine | | | | | | | | | |
| -- | -6.36 | ± | 0.02 | 3.13 | ± | 0.05 | 1.15 | ± | 0.06 |
| -7 | -5.85 | ± | 0.07 | 3.22 | ± | 0.13 | 1.14 | ± | 0.23 |
| -6.5 | -5.55 | ± | 0.05 | 3.17 | ± | 0.08 | 1.15 | ± | 0.09 |
| -6 | -5.19 | ± | 0.04 | 3.20 | ± | 0.08 | 1.15 | ± | 0.10 |
| -5.5 | -4.87 | ± | 0.06 | 3.16 | ± | 0.07 | 1.15 | ± | 0.08 |
| -5 | -4.58 | ± | 0.03 | 3.17 | ± | 0.03 | 1.19 | ± | 0.09 |
| xanomeline | | | | | | | | | |
| -- | -10.59 | ± | 0.06 | 3.76 | ± | 0.05 | 0.96 | ± | 0.07 |
| -7.5 | -10.16 | ± | 0.02 | 3.71 | ± | 0.02 | 1.02 | ± | 0.08 |
| -7 | -9.82 | ± | 0.04 | 3.69 | ± | 0.08 | 1.04 | ± | 0.09 |
| -6.5 | -9.19 | ± | 0.05 | 3.72 | ± | 0.07 | 1.03 | ± | 0.10 |
| -6 | -8.62 | ± | 0.05 | 3.69 | ± | 0.10 | 1.05 | ± | 0.07 |
| -5.5 | -7.95 | ± | 0.05 | 3.68 | ± | 0.06 | 1.02 | ± | 0.10 |
| -5 | -7.24 | ± | 0.02 | 3.71 | ± | 0.07 | 0.99 | ± | 0.08 |

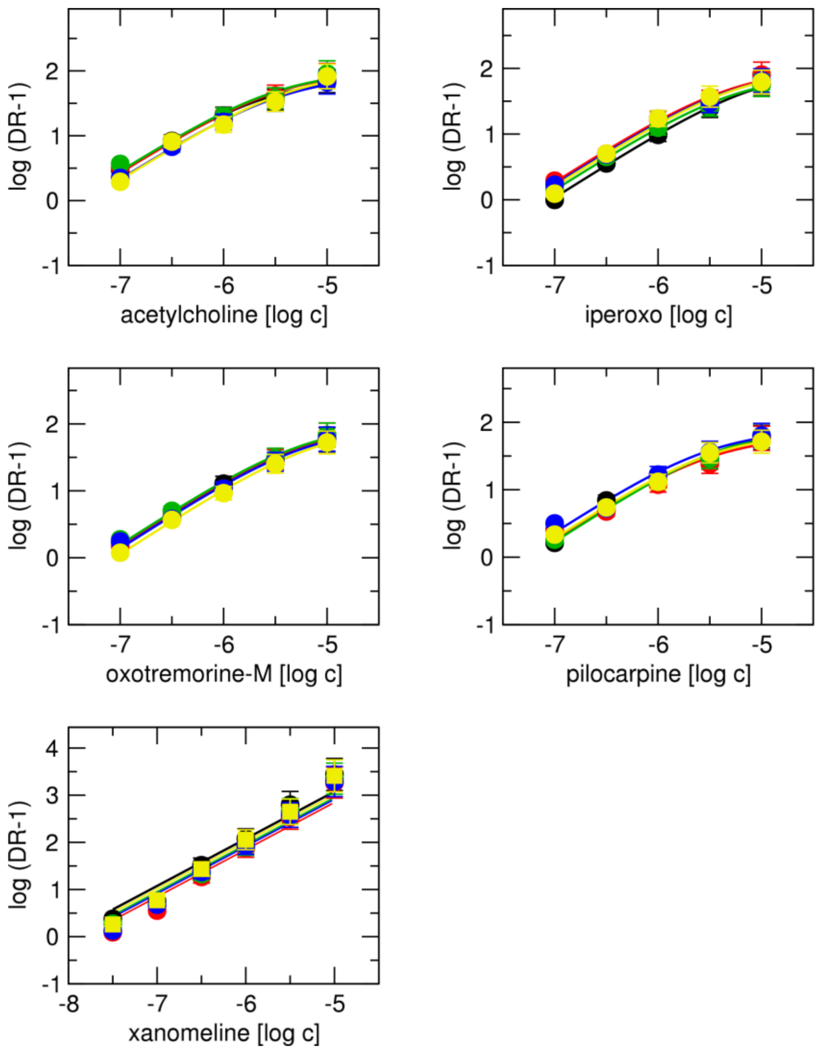

***Figure S1 Schild analysis of functional responses of $M_1$ receptor to agonists.***

*Schild plots of KH-5 effects on functional response to individual agonists are indicated in the legend. Abscissa, concentration of KH-5 expressed as a decadic logarithm of molar concentration. Ordinate, a decadic logarithm of DR-1, where DR is the ratio of agonist $EC_{50}$ in the presence of KH-5 to $EC_{50}$ in the absence of KH-5. Data are means ± SD from individual experiments performed in quadruplicate.*

***Table S4 Comparison of potencies ($pK_B$ derived from Schild analysis) of KH-5 to antagonise functional responses of individual agonists at $M_1$ receptor (Figure S1).***

| group1 | group2 | meandiff | p-adj | lower | upper | |
|---|---|---|---|---|---|---|
| acetylcholine | iperoxo | -0.224 | 0.0009 | -0.3636 | -0.0844 | TRUE |
| acetylcholine | oxotremorine-M | -0.256 | 0.0002 | -0.3956 | -0.1164 | TRUE |
| acetylcholine | pilocarp | -0.116 | 0.1336 | -0.2556 | 0.0236 | FALSE |
| acetylcholine | xanomeline | 0.548 | 0 | 0.4084 | 0.6876 | TRUE |
| iperoxo | oxotremorine-M | -0.032 | 0.9572 | -0.1716 | 0.1076 | FALSE |
| iperoxo | pilocarpine | 0.108 | 0.1811 | -0.0316 | 0.2476 | FALSE |
| iperoxo | xanomeline | 0.772 | 0 | 0.6324 | 0.9116 | TRUE |
| oxotremorine-M | pilocarpine | 0.14 | 0.0491 | 0.0004 | 0.2796 | TRUE |
| oxotremorine-M | xanomeline | 0.804 | 0 | 0.6644 | 0.9436 | TRUE |
| pilocarpine | xanomeline | 0.664 | 0 | 0.5244 | 0.8036 | TRUE |

Multiple Comparison of Means - Tukey HSD, FWER=0.05

***Table S5 Parameters of functional responses of $M_2$ muscarinic receptors to agonists in the presence of KH-5.***

*Half-efficient concentrations ($EC_{50}$), observed maximal responses to agonists ($E'_{MAX}$) and Hill coefficients of functional responses of $M_2$ receptors to agonists were obtained by fitting Equation 9 to the experimental data. The $EC_{50}$ values are expressed as the decadic logarithm of molar concentration. Data are means ± SD from five independent experiments.*

| log[KH-5] | $logEC_{50}$ | | | $E'_{MAX}$ | | | $n_H$ | | |
|---|---|---|---|---|---|---|---|---|---|
| acetylcholine | | | | | | | | | |
| -- | -6.13 | ± | 0.07 | 3.83 | ± | 0.09 | 0.73 | ± | 0.06 |
| -5.5 | -5.82 | ± | 0.05 | 3.69 | ± | 0.05 | 0.74 | ± | 0.05 |
| -5 | -5.42 | ± | 0.05 | 3.71 | ± | 0.06 | 0.78 | ± | 0.06 |
| -4.5 | -4.94 | ± | 0.08 | 3.78 | ± | 0.09 | 0.69 | ± | 0.07 |
| -4 | -4.58 | ± | 0.06 | 3.74 | ± | 0.07 | 0.75 | ± | 0.06 |
| iperoxo | | | | | | | | | |
| -- | -8.25 | ± | 0.05 | 3.95 | ± | 0.05 | 0.79 | ± | 0.06 |
| -5.5 | -8.03 | ± | 0.06 | 3.97 | ± | 0.07 | 0.74 | ± | 0.07 |
| -5 | -7.55 | ± | 0.04 | 3.98 | ± | 0.05 | 0.76 | ± | 0.05 |
| -4.5 | -7.09 | ± | 0.05 | 3.86 | ± | 0.05 | 0.75 | ± | 0.05 |
| -4 | -6.64 | ± | 0.06 | 3.96 | ± | 0.07 | 0.74 | ± | 0.07 |
| oxotremorine-M | | | | | | | | | |
| -- | -6.49 | ± | 0.05 | 3.51 | ± | 0.05 | 0.70 | ± | 0.04 |
| -5.5 | -6.35 | ± | 0.06 | 3.49 | ± | 0.05 | 0.78 | ± | 0.07 |
| -5 | -5.80 | ± | 0.06 | 3.55 | ± | 0.06 | 0.74 | ± | 0.05 |
| -4.5 | -5.29 | ± | 0.05 | 3.56 | ± | 0.05 | 0.73 | ± | 0.04 |
| -4 | -4.95 | ± | 0.04 | 3.45 | ± | 0.05 | 0.75 | ± | 0.04 |
| pilocarpine | | | | | | | | | |
| -- | -5.86 | ± | 0.06 | 3.24 | ± | 0.06 | 0.73 | ± | 0.06 |
| -5.5 | -5.78 | ± | 0.05 | 3.13 | ± | 0.04 | 0.85 | ± | 0.07 |
| -5 | -5.26 | ± | 0.05 | 3.20 | ± | 0.04 | 0.80 | ± | 0.06 |
| -4.5 | -4.68 | ± | 0.05 | 3.23 | ± | 0.04 | 0.73 | ± | 0.05 |
| -4 | -4.29 | ± | 0.06 | 3.21 | ± | 0.06 | 0.72 | ± | 0.06 |
| xanomeline | | | | | | | | | |
| -- | -8.80 | ± | 0.06 | 3.76 | ± | 0.06 | 0.76 | ± | 0.06 |
| -5.5 | -8.41 | ± | 0.06 | 3.75 | ± | 0.05 | 0.80 | ± | 0.08 |
| -5 | -7.91 | ± | 0.05 | 3.77 | ± | 0.04 | 0.75 | ± | 0.05 |
| -4.5 | -7.44 | ± | 0.06 | 3.67 | ± | 0.06 | 0.79 | ± | 0.08 |
| -4 | -6.90 | ± | 0.06 | 3.75 | ± | 0.06 | 0.75 | ± | 0.06 |

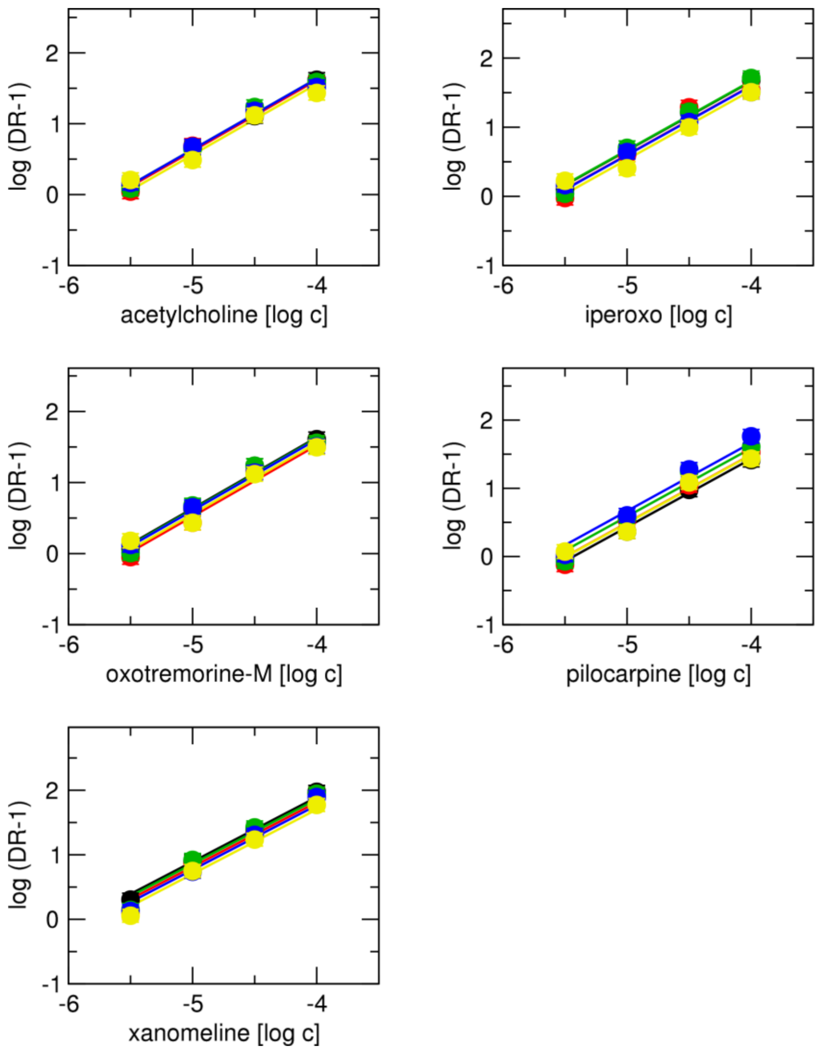

***Figure S2 Schild analysis of functional responses of $M_2$ receptor to agonists.***
*Schild plots of KH-5 effects on functional response to individual agonists are indicated in the legend. Abscissa, concentration of KH-5 expressed as a decadic logarithm of molar concentration. Ordinate, a decadic logarithm of DR-1, where DR is the ratio of agonist $EC_{50}$ in the presence of KH-5 to $EC_{50}$ in the absence of KH-5. Data are means ± SD from individual experiments performed in quadruplicate.*

***Table S6 Comparison of potencies ($pK_B$ derived from Schild analysis) of KH-5 to antagonise functional responses of individual agonists at $M_2$ receptor (Figure S2)***

| group1 | group2 | meandiff | p-adj | lower | upper | |
|---|---|---|---|---|---|---|
| acetylcholine | iperoxo | 0.002 | 1 | -0.1607 | 0.1647 | FALSE |
| acetylcholine | oxotremoine-M | -0.014 | 0.9989 | -0.1767 | 0.1487 | FALSE |
| acetylcholine | pilocarp | -0.06 | 0.8028 | -0.2227 | 0.1027 | FALSE |
| acetylcholine | xanomeline | 0.194 | 0.0147 | 0.0313 | 0.3567 | TRUE |
| iperoxo | oxotremoine-M | -0.016 | 0.9982 | -0.1787 | 0.1467 | FALSE |
| iperoxo | pilocarpine | -0.062 | 0.7838 | -0.2247 | 0.1007 | FALSE |
| iperoxo | xanomeline | 0.192 | 0.016 | 0.0293 | 0.3547 | TRUE |
| oxotremoine-M | pilocarpine | -0.046 | 0.9129 | -0.2087 | 0.1167 | FALSE |
| oxotremoine-M | xanomeline | 0.208 | 0.0083 | 0.0453 | 0.3707 | TRUE |
| pilocarpine | xanomeline | 0.254 | 0.0012 | 0.0913 | 0.4167 | TRUE |

Multiple Comparison of Means - Tukey HSD, FWER=0.05

***Table S7 Parameters of functional responses of $M_1$ muscarinic receptors in the cells treated by PBCM to agonists in the presence of KH-5.***

*Half-efficient concentrations ($EC_{50}$), observed maximal responses to agonists ($E'_{MAX}$) of functional responses of $M_1$ receptors to agonists were obtained by fitting Equation 9 to the experimental data. Hill coefficients were not different from 1. The $EC_{50}$ values are expressed as the negative decadic logarithm of molar concentration. Data are means ± SD from five independent experiments.*

| | control | | 1 µM KH-5 | | 10 µM KH-5 | |
|---|---|---|---|---|---|---|
| | $pEC_{50}$ | $E'_{MAX}$ | $pEC_{50}$ | $E'_{MAX}$ | $pEC_{50}$ | $E'_{MAX}$ |
| | | | acetylcholine | | | |
| Non-treated | 7.46 ± 0.02 | 0.86 ± 0.01 | 6.24 ± 0.02 | 0.84 ± 0.01 | 5.59 ± 0.05 | 0.84 ± 0.03 |
| PBCM 2 min | 7.00 ± 0.03 | 0.56 ± 0.01 | 5.78 ± 0.02 | 0.53 ± 0.01 | 5.19 ± 0.02 | 0.48 ± 0.01 |
| PBCM 5 min | 6.84 ± 0.01 | 0.34 ± 0.01 | 5.66 ± 0.02 | 0.31 ± 0.00* | 5.03 ± 0.01 | 0.28 ± 0.01* |
| | | | oxotremorine-M | | | |
| Non-treated | 7.71 ± 0.02 | 0.82 ± 0.01 | 6.73 ± 0.02 | 0.82 ± 0.01 | 6.02 ± 0.04 | 0.81 ± 0.02 |
| PBCM 2 min | 7.30 ± 0.05 | 0.50 ± 0.01 | 6.29 ± 0.03 | 0.51 ± 0.01 | 5.60 ± 0.02 | 0.48 ± 0.01 |
| PBCM 5 min | 7.10 ± 0.03 | 0.30 ± 0.00 | 6.10 ± 0.03 | 0.30 ± 0.01 | 5.45 ± 0.03 | 0.28 ± 0.01* |

*, different from control ($P < 0.05$ according to ANOVA and Tukey-HSD post-test)

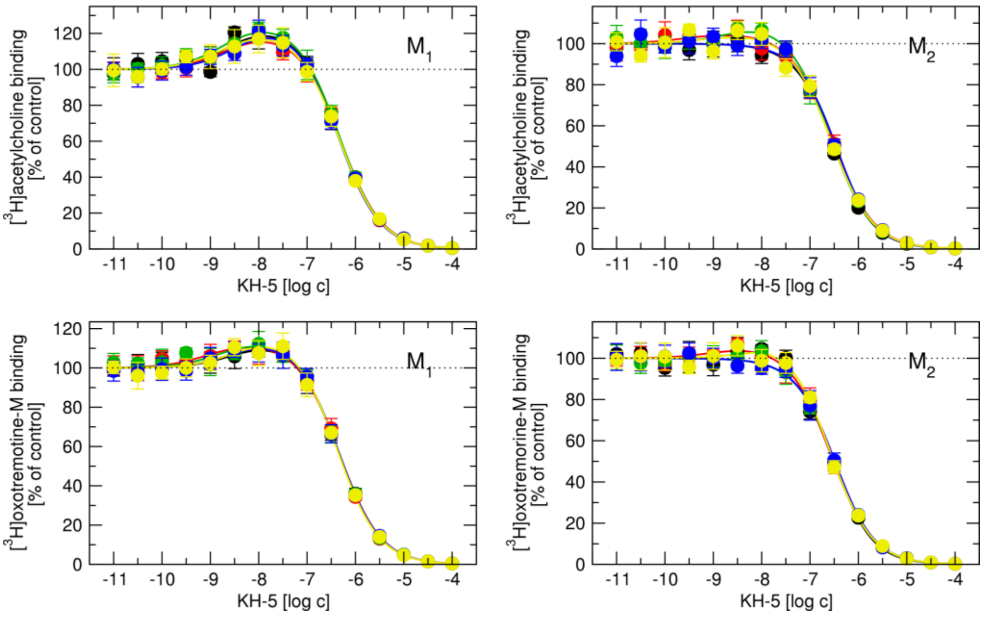

***Figure S3 Interaction between KH-5 and tritiated agonists at $M_1$ and $M_2$ receptors.***

*Effects of increasing concentrations of KH-5 on the binding of [$^3$H]acetylcholine (top row) and [$^3$H]oxotremorine-M (bottom row) to membranes from CHO cells expressing $M_1$ (left column) and $M_2$ (right column) receptors. Abscissa, concentration of KH-5 is expressed as a decadic logarithm of molar concentration. Ordinate, specific radioligand binding is expressed as a percentage of control in the absence of KH-5. Data are means ± SD from 5 individual experiments performed in quadruplicate.*

***Table S8 Estimates of binding energies of KH-5 docked to muscarinic receptors.***

Estimates of Vina/YASARA binding energies in kcal/mol of top poses of rescored docking poses of KH-5 to the orthosteric and ectopic binding sites of individual receptors in inactive and active conformations. The reported docking energies are Vina/YASARA scoring estimates used only for relative pose prioritisation; they are not treated as experimentally determined free energies.

| | $M_1$ | $M_2$ | $M_3$ | $M_4$ | $M_5$ | |
|---|---|---|---|---|---|---|
| (R)KH-5 | 6.954 | 5.852 | 7.774 | 6.335 | 7.104 | orthosteric |
| (S)KH-5 | 7.006 | 6.109 | 7.762 | 6.859 | 7.120 | |
| (R)KH-5 | 5.594 | 6.044 | 7.004 | 6.558 | 6.107 | ectopic inactive |
| (S)KH-5 | 5.314 | 6.017 | 6.910 | 6.394 | 5.567 | |
| (R)KH-5 | 6.534 | 6.316 | | 4.980 | | ectopic active |
| (S)KH-5 | 5.960 | 4.875 | | 4.989 | | |

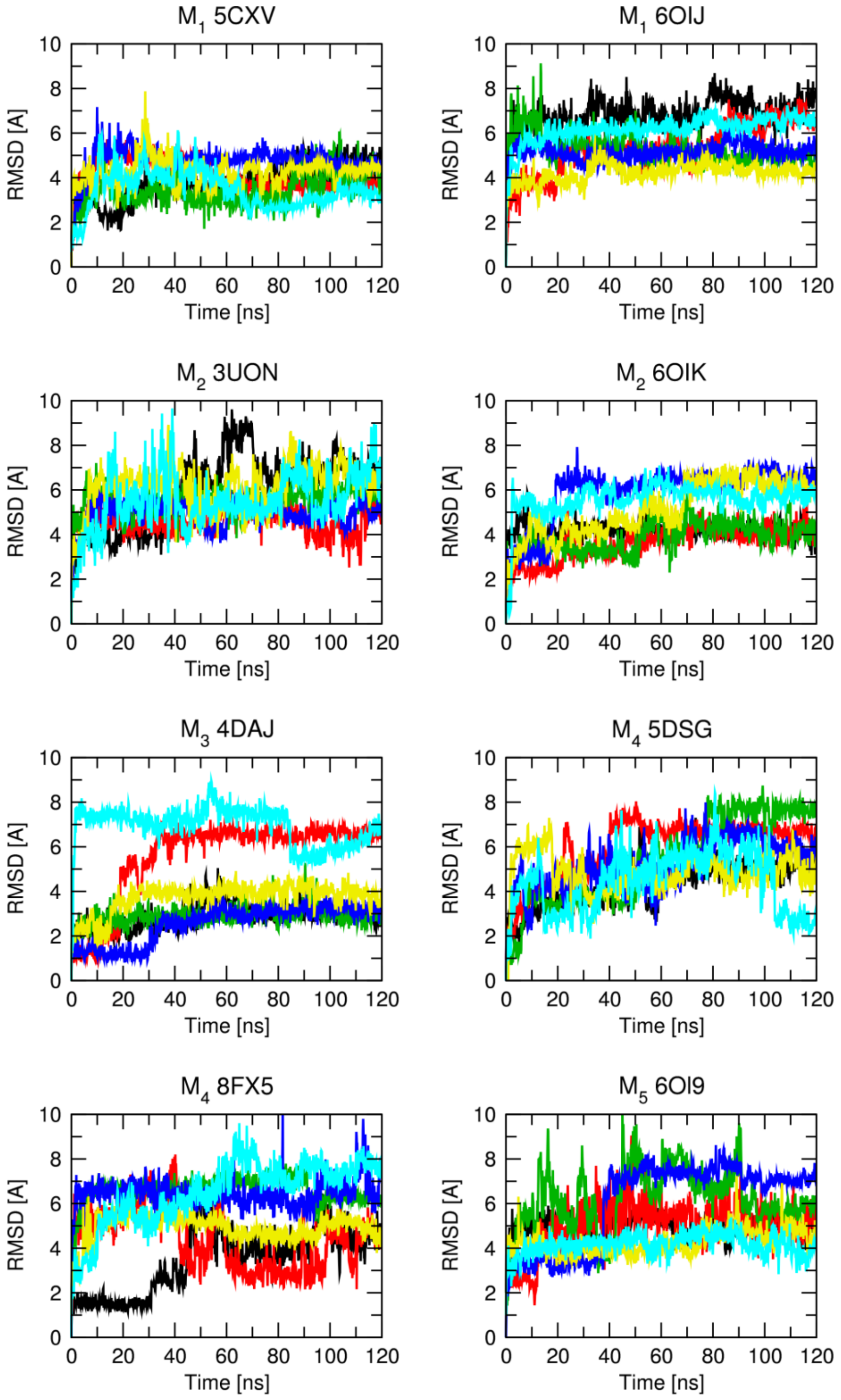

***Figure S4 RMSDs of KH-5 in the ectopic site***

*The protein-ligand complex was first aligned on the protein backbone of the reference (initial time frame), and then the RMSD of the KH-5 heavy atoms was calculated. Traces from individual MD runs of (R)KH-5 (black, red, green) and (S)KH-5 (blue, yellow, cyan).*

***Table S9 Last 100 ns of MD RMSD of KH-5 in the ectopic site.***

*The protein-ligand complex was first aligned on the protein backbone of the reference (initial time frame), and then the RMSD of the ligand heavy atoms was calculated. Data are means ± SD of RMSDs of the last 100 ns of individual MD simulations.*

| Structure | Ligand | run | RMSD | | |
|---|---|---|---|---|---|
| $M_1$ 5CXV | (R)KH-5 | 1 | 4.51 | ± | 1.50 |
| $M_1$ 5CXV | (R)KH-5 | 2 | 4.06 | ± | 0.57 |
| $M_1$ 5CXV | (R)KH-5 | 3 | 6.49 | ± | 0.69 |
| $M_1$ 5CXV | (S)KH-5 | 1 | 4.83 | ± | 0.25 |
| $M_1$ 5CXV | (S)KH-5 | 2 | 5.33 | ± | 1.77 |
| $M_1$ 5CXV | (S)KH-5 | 3 | 4.46 | ± | 2.13 |
| $M_1$ 6OIJ | (R)KH-5 | 1 | 6.96 | ± | 0.61 |
| $M_1$ 6OIJ | (R)KH-5 | 2 | 5.77 | ± | 0.76 |
| $M_1$ 6OIJ | (R)KH-5 | 3 | 5.18 | ± | 0.48 |
| $M_1$ 6OIJ | (S)KH-5 | 1 | 5.14 | ± | 0.34 |
| $M_1$ 6OIJ | (S)KH-5 | 2 | 4.34 | ± | 0.32 |
| $M_1$ 6OIJ | (S)KH-5 | 3 | 6.36 | ± | 0.29 |
| $M_2$ 3UON | (R)KH-5 | 1 | 6.52 | ± | 1.34 |
| $M_2$ 3UON | (R)KH-5 | 2 | 4.60 | ± | 0.58 |
| $M_2$ 3UON | (R)KH-5 | 3 | 5.63 | ± | 0.44 |
| $M_2$ 3UON | (S)KH-5 | 1 | 6.05 | ± | 0.88 |
| $M_2$ 3UON | (S)KH-5 | 2 | 6.69 | ± | 2.55 |
| $M_2$ 3UON | (S)KH-5 | 3 | 5.75 | ± | 1.03 |
| $M_2$ 6OIK | (R)KH-5 | 1 | 4.23 | ± | 0.30 |
| $M_2$ 6OIK | (R)KH-5 | 2 | 4.88 | ± | 0.49 |
| $M_2$ 6OIK | (R)KH-5 | 3 | 4.99 | ± | 0.67 |
| $M_2$ 6OIK | (S)KH-5 | 1 | 6.42 | ± | 0.36 |
| $M_2$ 6OIK | (S)KH-5 | 2 | 5.51 | ± | 0.98 |
| $M_2$ 6OIK | (S)KH-5 | 3 | 5.80 | ± | 0.37 |
| $M_3$ 4DAJ | (R)KH-5 | 1 | 3.05 | ± | 0.47 |
| $M_3$ 4DAJ | (R)KH-5 | 2 | 6.40 | ± | 0.52 |
| $M_3$ 4DAJ | (R)KH-5 | 3 | 4.96 | ± | 0.36 |
| $M_3$ 4DAJ | (S)KH-5 | 1 | 4.87 | ± | 0.60 |
| $M_3$ 4DAJ | (S)KH-5 | 2 | 3.97 | ± | 0.31 |
| $M_3$ 4DAJ | (S)KH-5 | 3 | 6.86 | ± | 0.83 |
| $M_4$ 5DSG | (R)KH-5 | 1 | 4.67 | ± | 0.72 |
| $M_4$ 5DSG | (R)KH-5 | 2 | 6.53 | ± | 1.27 |
| $M_4$ 5DSG | (R)KH-5 | 3 | 6.02 | ± | 1.65 |
| $M_4$ 5DSG | (S)KH-5 | 1 | 5.49 | ± | 0.96 |
| $M_4$ 5DSG | (S)KH-5 | 2 | 6.42 | ± | 0.63 |
| $M_4$ 5DSG | (S)KH-5 | 3 | 4.34 | ± | 3.62 |
| $M_4$ 8FX5 | (R)KH-5 | 1 | 5.96 | ± | 1.15 |
| $M_4$ 8FX5 | (R)KH-5 | 2 | 4.19 | ± | 1.38 |
| $M_4$ 8FX5 | (R)KH-5 | 3 | 6.68 | ± | 0.44 |
| $M_4$ 8FX5 | (S)KH-5 | 1 | 5.78 | ± | 0.64 |
| $M_4$ 8FX5 | (S)KH-5 | 2 | 4.95 | ± | 0.40 |
| $M_4$ 8FX5 | (S)KH-5 | 3 | 6.98 | ± | 0.99 |
| $M_5$ 6OL9 | (R)KH-5 | 1 | 4.92 | ± | 0.47 |
| $M_5$ 6OL9 | (R)KH-5 | 2 | 6.86 | ± | 5.05 |
| $M_5$ 6OL9 | (R)KH-5 | 3 | 6.44 | ± | 1.25 |
| $M_5$ 6OL9 | (S)KH-5 | 1 | 6.73 | ± | 1.39 |
| $M_5$ 6OL9 | (S)KH-5 | 2 | 4.33 | ± | 0.55 |
| $M_5$ 6OL9 | (S)KH-5 | 3 | 4.25 | ± | 0.42 |

***Table S10 Conserved residues in ectopic site***

*Subtype list of amino acid residues in the ectopic site. Residue position according to Ballestero-Weinstein numbering.*

| position | $M_1$ | $M_2$ | $M_3$ | $M_4$ | $M_5$ |
|---|---|---|---|---|---|
| 2.61 | Y | Y | Y | Y | Y |
| 2.64 | Y | Y | Y | Y | Y |
| 3.28 | W | W | W | W | W |
| 3.31 | Y | Y | Y | Y | Y |
| 3.33 | Y | Y | Y | Y | Y |
| 4.57 | W | W | W | W | W |
| 45.51 | Y | Y | F | F | Q |
| 45.52 | I | I | I | I | I |
| 45.55 | F | F | F | F | F |
| 5.46 | A | A | A | A | A |
| 5.47 | F | F | F | F | F |
| 6.48 | W | W | W | W | W |
| 6.51 | Y | Y | Y | Y | Y |
| 7.32 | N | N | K | D | K |
| 7.35 | W | W | W | W | W |
| 7.36 | E | T | N | S | T |
| 7.39 | Y | Y | Y | Y | Y |
| 7.40 | W | W | W | W | W |

***Table S11 Binding parameters of KH-5 to membranes of CHO cells expressing individual subtypes of muscarinic receptors.***

*Inhibition constants ($K_I$) were obtained by fitting Equation 2 to the data from competition experiments with [$^3$H]NMS. Constants are expressed as negative decadic logarithms. Data are means ± SD from 5 independent experiments performed in quadruplicates.*

| | $pK_I$ | | |
|---|---|---|---|
| $M_1$ | 6.60 | ± | 0.05 |
| $M_2$ | 6.92 | ± | 0.05 |
| $M_3$ | 6.48 | ± | 0.05 |
| $M_4$ | 6.46 | ± | 0.05 |
| $M_5$ | 6.82 | ± | 0.05 |

***Table S12 Binding parameters of radioligands to membranes of CHO cells expressing wild-type (WT) and mutant $M_1$ receptors.***

*Equilibrium dissociation constants ($K_D$) of tritiated N-methylscopolamine (NMS) and acetylcholine were obtained by fitting Equation 1 to the data from saturation binding experiments and are expressed as negative decadic logarithms. Maximum binding capacity is expressed as pmol of binding sites per mg of membrane proteins. Data are means ± SD or maximum to minimum values from 5 independent experiments performed in quadruplicates.*

| | [$^3$H]NMS | | | | | | [$^3$H]acetylcholine | | | | | |
|---|---|---|---|---|---|---|---|---|---|---|---|---|
| receptor | $pK_D$ | | | $B_{MAX}$ [pmol/mg prot.] | | | $pK_D$ | | | $B_{MAX}$ [pmol/mg prot.] | | |
| $M_1$ WT | 9.75 | ± | 0.08 | 1.05 | to | 3.95 | 7.19 | ± | 0.03 | 0.21 | to | 0.8 |
| $M_1$ Y82A | 9.41 | ± | 0.08* | 0.17 | to | 2.67 | 7.10 | ± | 0.04* | 0.12 | to | 0.54 |
| $M_1$ Y85A | 9.36 | ± | 0.05* | 0.59 | to | 1.65 | 7.10 | ± | 0.03* | 0.12 | to | 0.31 |
| $M_1$ Y179A | 9.46 | ± | 0.08* | 0.35 | to | 2.7 | 7.18 | ± | 0.02 | 0.07 | to | 0.65 |
| $M_1$ W400A | 9.44 | ± | 0.07* | 0.40 | to | 0.96 | 6.79 | ± | 0.03* | 0.08 | to | 0.19 |
| $M_1$ Y404A | 8.36 | ± | 0.02* | 0.18 | to | 2.34 | 6.60 | ± | 0.04* | 0.03 | to | 0.47 |

*, different from WT ($P < 0.05$ according to ANOVA and Tukey-HSD post-test)

***Table S13 Binding parameters of KH-5 to membranes of CHO cells expressing wild-type (WT) and mutant $M_1$ receptors.***

*Inhibition constants ($K_I$) were obtained by fitting Equation 2 to the data from competition experiments with [$^3$H]NMS. Equilibrium dissociation constants of KH-5 for the allosteric ($K_A$) and orthosteric ($K_B$) sites, respectively, and the factor of binding cooperativity (α) between [$^3$H]acetylcholine and KH-5 were obtained by fitting Equation 8 to the data from competition-like experiments (main document Figure 8). Constants are expressed as negative decadic logarithms. Data are means ± SD from 5 independent experiments performed in quadruplicates.*

| | [$^3$H]NMS | [$^3$H]acetylcholine | | |
|---|---|---|---|---|
| receptor | $pK_i$ | $pK_A$ | α | $pK_B$ |
| $M_1$ wt | 6.85 ± 0.13 | 8.46 ± 0.23 | 1.69 ± 0.09 | 6.89 ± 0.04 |
| $M_1$ Y82A | 6.24 ± 0.07* | n.d. | n.d. | 6.25 ± 0.02* |
| $M_1$ Y85A | 6.55 ± 0.11* | n.d. | n.d. | 6.55 ± 0.01* |
| $M_1$ Y179A | 6.44 ± 0.06* | n.d. | n.d. | 6.43 ± 0.02* |
| $M_1$ W400A | 6.53 ± 0.12* | n.d. | n.d. | 6.54 ± 0.01* |
| $M_1$ Y404A | 6.86 ± 0.09 | 8.53 ± 0.22 | 2.36 ± 0.29 | 6.88 ± 0.07 |

n.d., not determined; *, different from WT ($P < 0.05$ according to ANOVA and Tuckey-HSD post-test).

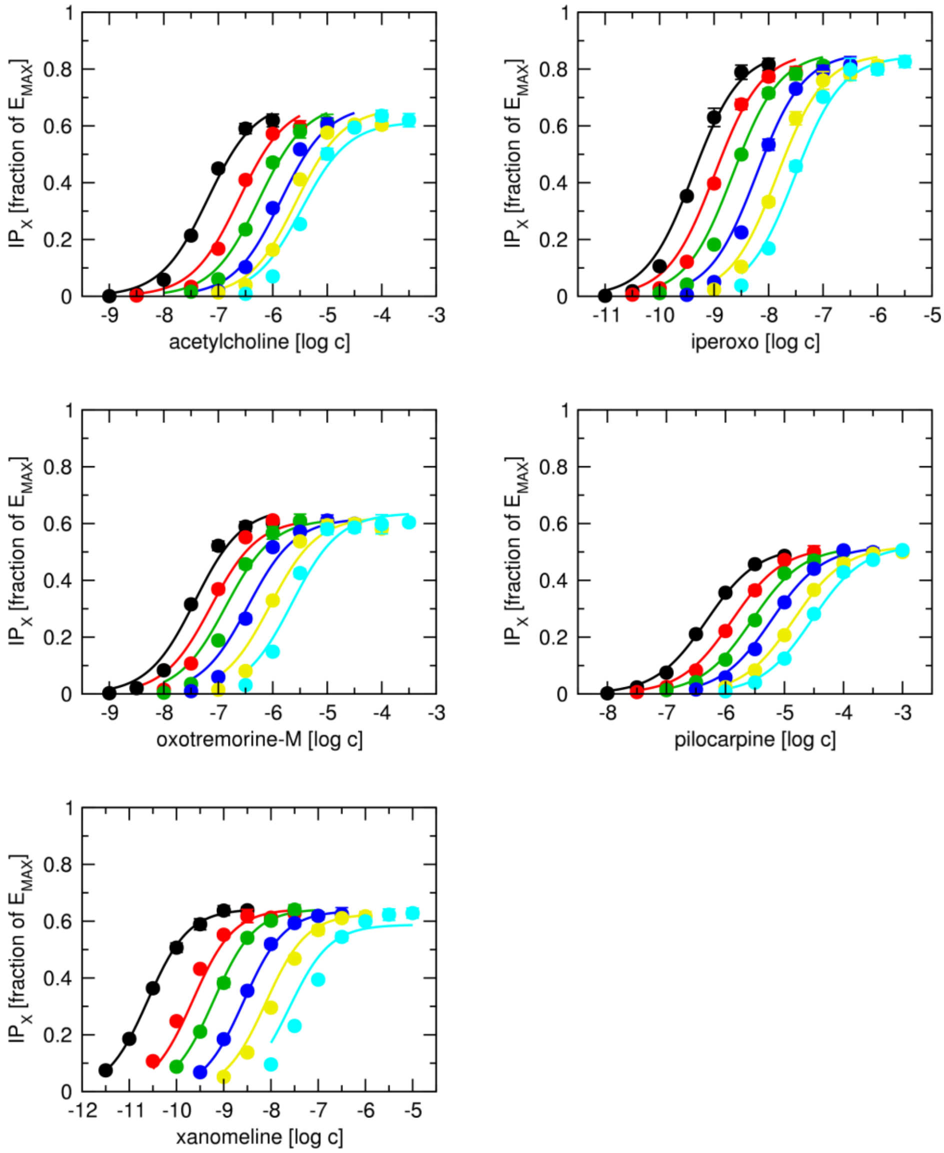

***Figure S5 Fitting of the operational model of duasterically modulated agonism (OMDMA) to functional responses at $M_1$ receptor.***

Data are pooled data of functional response of $M_1$ receptors to agonists (acetylcholine, iperoxo, oxotremorine-M, pilocarpine and xanomeline) in the absence or presence of antagonist KH-5 at concentrations indicated in the legends, measured as accumulation of inositol phosphates ($IP_X$) in CHO cells shown in Figure 2 of the main manuscript, expressed as a fraction of system maximal response ($E_{MAX}$). Curves are global 3D-fits of Equation 13 of the main manuscript, where the concentrations of an agonist and KH-5 are two independent variables. $K_A$ and τ were fixed to values predetermined in control experiments. Data are means ± SD from five experiments performed in quadruplicate.

***Table S14 Estimates of parameters ± SD of OMDMA fitted to data in Figure S5***

Equilibrium dissociation constant of agonist ($K_A$) and its operational efficacy (τ) were obtained by fitting **Error! Reference source not found.** to the functional response data in the absence of KH-5. Equilibrium dissociation constants of KH-5 for the allosteric ($K_B$) and orthosteric ($K_C$) binding sites, respectively, and are factors of binding coopertivity between molecule of KH-5 bound to the allosteric binding site and molecule of agonist (α) or the second molecule of KH-5 (β) bound to the orthosteric binding site and cooperativity factor of operational efficacy (δ) were obtained by global fit of Equation 13 of the main manuscript to all the data.

| | acetylcholine | | | iperoxo | | | oxotremorine-M | | | pilocarpine | | | xanomeline | | |
|---|---|---|---|---|---|---|---|---|---|---|---|---|---|---|---|
| $pK_A$ | 6.69 | ± | 0.11 | 8.50 | ± | 0.16 | 6.97 | ± | 0.12 | 6.00 | ± | 0.04 | 10.16 | ± | 0.02 |
| τ | 2.18 | ± | 0.30 | 6.23 | ± | 1.78 | 1.90 | ± | 0.28 | 1.08 | ± | 0.04 | 1.80 | ± | 0.03 |
| $pK_B$ | 3.34 | ± | 4.26 | 3.60 | ± | 3.88 | 3.11 | ± | 4.60 | 3.00 | ± | 4.16 | 4.55 | ± | 3.17 |
| $pK_C$ | 7.46 | ± | 0.24 | 7.31 | ± | 0.24 | 7.01 | ± | 0.74 | 7.24 | ± | 0.02 | 8.03 | ± | 0.08 |
| log(α) | 1.08 | ± | 6.39 | 1.74 | ± | 5.25 | 1.18 | ± | 6.42 | 1.14 | ± | 6.10 | -1.98 | ± | 8.17 |
| log(β) | 1.48 | ± | 5.23 | 1.88 | ± | 6.33 | 1.71 | ± | 5.85 | 1.05 | ± | 4.93 | 2.00 | ± | 6.32 |
| log(δ) | 1.12 | ± | 6.39 | -0.05 | ± | 5.25 | 0.75 | ± | 6.42 | 1.20 | ± | 6.10 | -2.00 | ± | 8.17 |

**A**

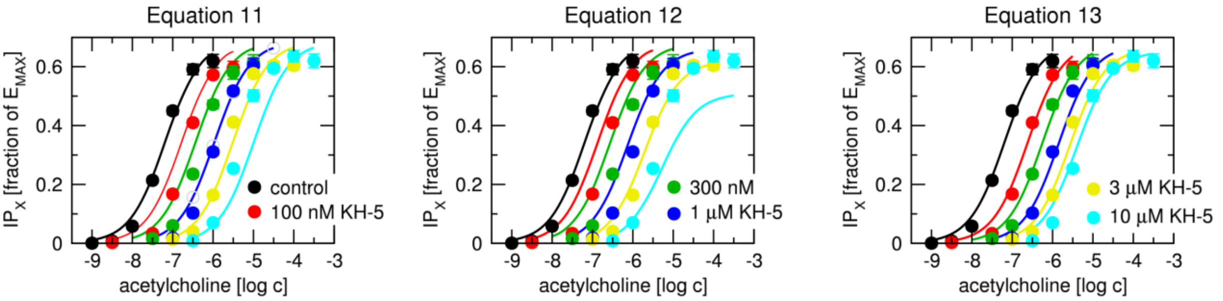

```
--- Model AICc ---
  n = 42
  k = 1.0
  SS_res = 1.093615e-01
  MSE = 2.603845e-03
  log-likelihood = 65.3707
  AIC = -128.74
  AICc = -128.64

--- Residual analysis ---
  Mean: -1.53e-02
  Std Dev: 4.87e-02
  Min / Max: -9.15e-02 / 1.61e-01
  Shapiro-Wilk p-value: 0.0012
```

```
--- Model AICc ---
  n = 42
  k = 2.0
  SS_res = 2.088115e-01
  MSE = 4.971702e-03
  log-likelihood = 51.7884
  AIC = -99.58
  AICc = -99.27

--- Residual analysis ---
  Mean: -1.69e-02
  Std Dev: 6.85e-02
  Min / Max: -1.30e-01 / 1.67e-01
  Shapiro-Wilk p-value: 0.0012
```

```
--- Model AICc ---
  n = 42
  k = 5.0
  SS_res = 3.987945e-02
  MSE = 9.495108e-04
  log-likelihood = 86.5554
  AIC = -163.11
  AICc = -161.44

--- Residual analysis ---
  Mean: -8.82e-03
  Std Dev: 2.95e-02
  Min / Max: -6.14e-02 / 5.27e-02
  Shapiro-Wilk p-value: 0.0383
```

**B**

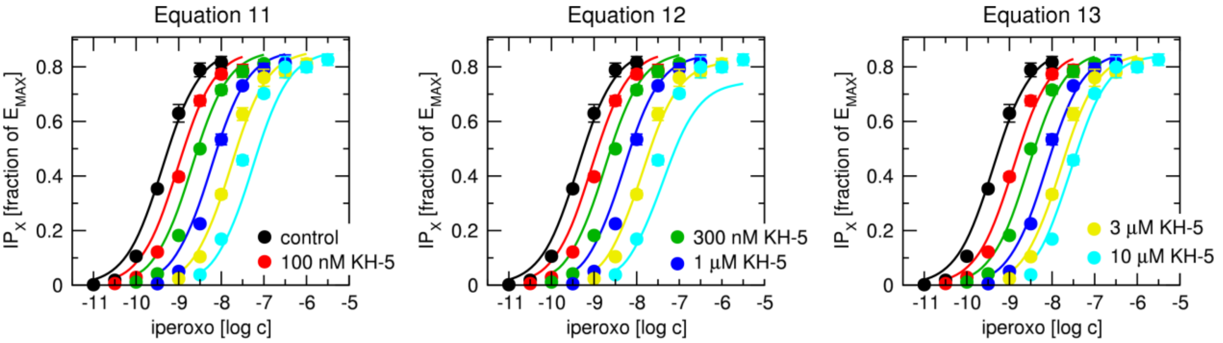

```
--- Model AICc ---
  n = 42
  k = 1.0
  SS_res = 1.140922e-01
  MSE = 2.716481e-03
  log-likelihood = 64.4814
  AIC = -126.96
  AICc = -126.86

--- Residual analysis ---
  Mean: -1.51e-02
  Std Dev: 4.99e-02
  Min / Max: -1.03e-01 / 1.43e-01
  Shapiro-Wilk p-value: 0.0008
```

```
--- Model AICc ---
  n = 42
  k = 2.0
  SS_res = 1.693147e-01
  MSE = 4.031302e-03
  log-likelihood = 56.1916
  AIC = -108.38
  AICc = -108.08

--- Residual analysis ---
  Mean: -1.31e-02
  Std Dev: 6.21e-02
  Min / Max: -1.26e-01 / 1.68e-01
  Shapiro-Wilk p-value: 0.0056
```

```
--- Model AICc ---
  n = 42
  k = 5.0
  SS_res = 5.022832e-02
  MSE = 1.195912e-03
  log-likelihood = 81.7103
  AIC = -153.42
  AICc = -151.75

--- Residual analysis ---
  Mean: -9.99e-03
  Std Dev: 3.31e-02
  Min / Max: -6.15e-02 / 5.97e-02
  Shapiro-Wilk p-value: 0.0371
```

**C**

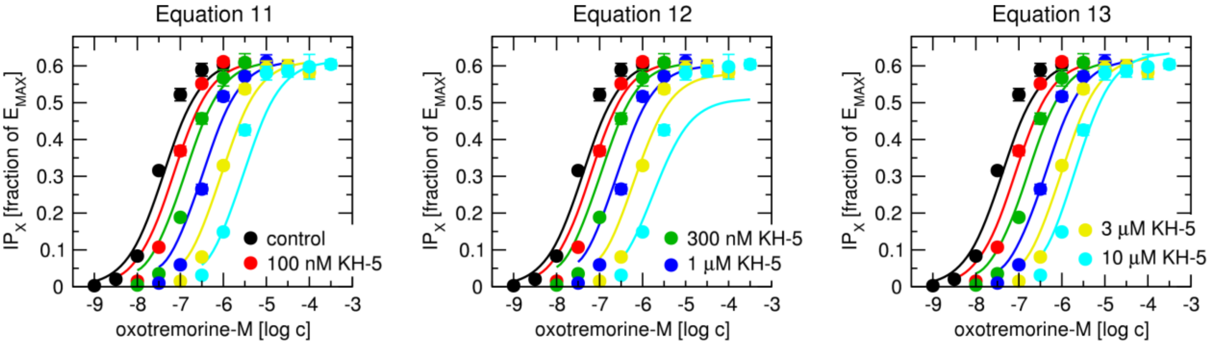

```
--- Model AICc ---
  n = 42
  k = 1.0
  SS_res = 9.726727e-02
  MSE = 2.315887e-03
  log-likelihood = 67.8318
  AIC = -133.66
  AICc = -133.56

--- Residual analysis ---
  Mean: -1.27e-03
  Std Dev: 4.81e-02
  Min / Max: -8.59e-02 / 1.08e-01
  Shapiro-Wilk p-value: 0.4845
```

```
--- Model AICc ---
  n = 42
  k = 2.0
  SS_res = 1.674430e-01
  MSE = 3.986738e-03
  log-likelihood = 56.4250
  AIC = -108.85
  AICc = -108.54

--- Residual analysis ---
  Mean: -2.99e-04
  Std Dev: 6.31e-02
  Min / Max: -1.12e-01 / 1.48e-01
  Shapiro-Wilk p-value: 0.5821
```

```
--- Model AICc ---
  n = 42
  k = 5.0
  SS_res = 7.733251e-02
  MSE = 1.841250e-03
  log-likelihood = 72.6481
  AIC = -135.30
  AICc = -133.63

--- Residual analysis ---
  Mean: -1.14e-03
  Std Dev: 4.29e-02
  Min / Max: -6.97e-02 / 8.58e-02
  Shapiro-Wilk p-value: 0.1340
```

**D**

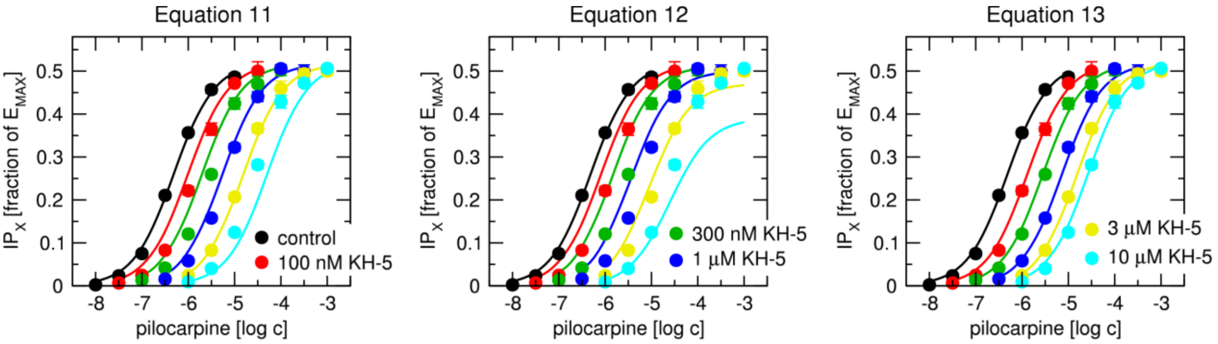

```
--- Model AICc ---
  n = 42
  k = 1.0
  SS_res = 3.242893e-02
  MSE = 7.721174e-04
  log-likelihood = 90.8984
  AIC = -179.80
  AICc = -179.70

--- Residual analysis ---
  Mean: -4.48e-03
  Std Dev: 2.74e-02
  Min / Max: -5.98e-02 / 8.33e-02
  Shapiro-Wilk p-value: 0.0008
```

```
--- Model AICc ---
  n = 42
  k = 2.0
  SS_res = 9.593553e-02
  MSE = 2.284179e-03
  log-likelihood = 68.1213
  AIC = -132.24
  AICc = -131.93

--- Residual analysis ---
  Mean: -7.35e-03
  Std Dev: 4.72e-02
  Min / Max: -9.23e-02 / 1.26e-01
  Shapiro-Wilk p-value: 0.0036
```

```
--- Model AICc ---
  n = 42
  k = 5.0
  SS_res = 5.375369e-03
  MSE = 1.279850e-04
  log-likelihood = 128.6401
  AIC = -247.28
  AICc = -245.61

--- Residual analysis ---
  Mean:  -2.19e-03
  Std Dev: 1.11e-02
  Min / Max: -1.93e-02 / 2.19e-02
  Shapiro-Wilk p-value: 0.0082
```

**E**

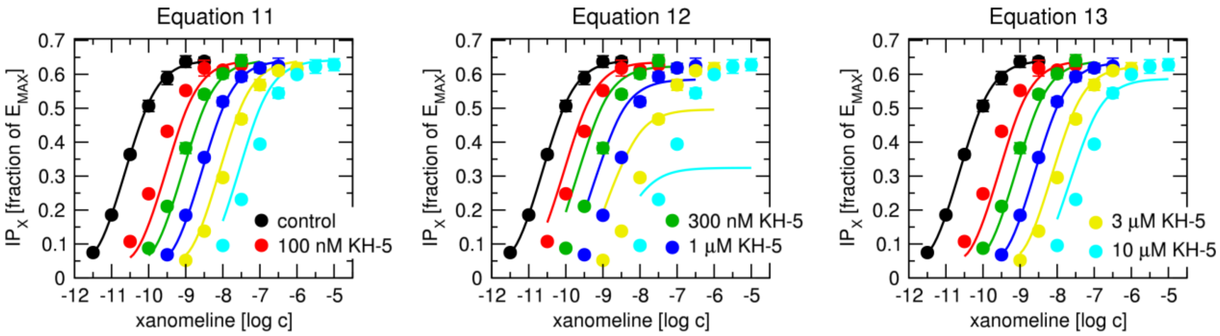

```
--- Model AICc ---
  n = 42
  k = 1.0
  SS_res = 7.208625e-02
  MSE = 1.716339e-03
  log-likelihood = 74.1234
  AIC = -146.25
  AICc = -146.15

--- Residual analysis ---
  Mean: -2.00e-03
  Std Dev: 4.14e-02
  Min / Max: -1.08e-01 / 1.06e-01
  Shapiro-Wilk p-value: 0.0975
```

```
--- Model AICc ---
  n = 42
  k = 2.0
  SS_res = 5.854000e-01
  MSE = 1.393810e-02
  log-likelihood = 30.1403
  AIC = -56.28
  AICc = -55.97

--- Residual analysis ---
  Mean: -5.42e-03
  Std Dev: 1.18e-01
  Min / Max: -1.92e-01 / 3.04e-01
  Shapiro-Wilk p-value: 0.0016
```

```
--- Model AICc ---
  n = 42
  k = 5.0
  SS_res = 6.000948e-02
  MSE = 1.428797e-03
  log-likelihood = 77.9739
  AIC = -145.95
  AICc = -144.28

--- Residual analysis ---
  Mean: 2.37e-03
  Std Dev: 3.77e-02
  Min / Max: -1.03e-01 / 9.56e-02
  Shapiro-Wilk p-value: 0.0387
```

***Figure S6 Comparison of fitting of models to functional response data***

*Data are pooled data of functional response of $M_1$ receptors to acetylcholine (**A**), iperoxo (**B**), oxotremorine-M (**C**), pilocarpine (**D**) and xanomeline (**E**) in the absence or presence of antagonist KH-5 at concentrations indicated in the legends, measured as accumulation of inositol phosphates ($IP_X$) in CHO cells shown in Figure 2 of the main manuscript, expressed as a fraction of system maximal response ($E_{MAX}$). Curves are global 3D-fits of Equation 11, describing competitive inhibition, Equation 12, describing allosteric inhibition, and Equation 13, describing dualsteric inhibition, where the concentrations of an agonist and KH-5 are two independent variables. $K_A$ and τ were fixed to values predetermined in control experiments. Data are means ± SD from five experiments performed in quadruplicate. Model AICc and residual analysis are in the insets.*

### Derivations

#### Dualsteric interaction

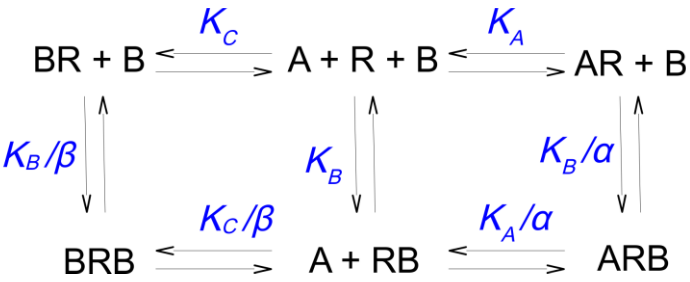

Tracer A binds to the orthosteric site with equilibrium dissociation constant $K_A$. Dualsteric ligand B binds to the allosteric binding site with equilibrium dissociation constant $K_B$ and to the orthosteric binding site with equilibrium dissociation constant $K_C$. Alpha is a factor of binding cooperativity between the orhosteric tracer A and dualsteric ligand B. Beta is a factor of binding cooperativity between two molecules of dualsteric ligand B.

$$K_A = \frac{[A][R]}{[AR]}$$

$$K_B = \frac{[B][R]}{[RB]}$$

$$K_C = \frac{[B][R]}{[BR]}$$

$$\frac{K_A}{\alpha} = \frac{[A][RB]}{[ARB]}$$

$$\frac{K_B}{\alpha} = \frac{[AR][B]}{[ARB]}$$

$$\frac{K_C}{\beta} = \frac{[B][RB]}{[BRB]}$$

$$\frac{K_B}{\beta} = \frac{[B][RB]}{[BRB]}$$

The total concentration of receptor species is given by the sum:

$$[R_T] = [R] + [AR] + [RB] + [ARB] + [BR] + [BRB]$$

Fraction of receptors occupied by tracer A:

$$\frac{[AR]+[ARB]}{[R_T]} = \frac{[AR]+[ARB]}{[R]+[AR]+[RB]+[ARB]+[BR]+[BRB]}$$

Multiplying the numerator and denominator of the fraction on the right side by 1/[A][R] gives:

$$\frac{[AR]+[ARB]}{[R_T]} = \frac{\frac{[AR]}{[A][R]}+\frac{[ARB]}{[A][R]}}{\frac{[R]}{[A][R]}+\frac{[AR]}{[A][R]}+\frac{[RB]}{[A][R]}+\frac{[ARB]}{[A][R]}+\frac{[BR]}{[A][R]}+\frac{[BRB]}{[A][R]}}$$

After substitution:

$$\frac{[AR]+[ARB]}{[R_T]} = \frac{\frac{1}{K_A}+\frac{\alpha[B]}{K_A K_B}}{\frac{1}{K_A}+\frac{1}{[A]}+\frac{[B]}{[A]K_B}+\frac{\alpha[B]}{K_A K_B}+\frac{[B]}{[A]K_C}+\frac{\beta[B]}{K_C K_B}}$$

After multiplication and rearrangement:

$$\frac{[AR]+[ARB]}{[R_T]} = \frac{[A]K_C(\alpha[B]+K_B)}{\alpha K_C[A][B]+\beta K_A[A][B]+K_A K_B K_C+K_A K_B[B]+K_A K_C[B]+K_B K_C[A]}$$

$$\frac{[AR]+[ARB]}{[R_T]} = \frac{[A]K_C(\alpha[B]+K_B)}{K_C[A](\alpha[B]+K_B)+K_A([B](\beta[A]+K_C)+K_B([B]+K_C))}$$

After division by $K_C(\alpha[B]+K_B)$ and rearrangement:

$$\frac{[AR]+[ARB]}{[R_T]} = \frac{[A]}{[A]+ K_A\frac{[B](\beta[A]+K_C)+K_B([B]+K_C)}{K_C(\alpha[B]+K_B)}}$$

After division by binding in the absence of B $\{Y=[A]/([A]+K_A)\}$ we get normalised binding:

$$\frac{Y\prime}{Y} = \frac{[A]+ K_A}{[A]+ K_A\frac{[B](\beta[A]+K_C)+K_B([B]+K_C)}{K_C(\alpha[B]+K_B)}}$$

### Operational model of competitive inhibition

#### *Binding:*

$$K_A = \frac{[A][R]}{[AR]} \Rightarrow [R] = \frac{K_A[AR]}{[A]}$$

$$[R] \Rightarrow \frac{[R]}{[A][R]} = \frac{1}{[A]}$$

$$[AR] \Rightarrow \frac{[AR]}{[A][R]} = \frac{1}{K_A}$$

$$K_B = \frac{[R][B]}{[BR]} \Rightarrow [R] = \frac{K_B[BR]}{[B]}$$

$$[BR] \Rightarrow \frac{[BR]}{[A][R]} = \frac{[BR][B]}{[A]K_B[BR]} = \frac{[B]}{[A]K_B}$$

$$R_T = [R] + [AR] + [BR]$$

$$\frac{[AR]}{[R_T]} = \frac{\frac{1}{K_A}}{\frac{1}{[A]}+\frac{1}{K_A}+\frac{[B]}{[A]K_B}} = \frac{1}{\frac{K_A}{[A]}+\frac{K_A}{K_A}+\frac{K_A[B]}{[A]K_B}} = \frac{1}{\frac{K_A}{[A]}+1+\frac{K_A[B]}{[A]K_B}} = \frac{[A]}{K_A+[A]+\frac{[B]K_A}{K_B}}$$

$$[AR] = [R_T]\frac{[A]K_B}{[A]K_B+K_A([B]+K_B)}$$

$$\frac{[BR]}{[R_T]} = \frac{\frac{[B]}{[A]K_B}}{\frac{1}{[A]}+\frac{1}{K_A}+\frac{[B]}{[A]K_B}} = \frac{1}{\frac{[A]K_B}{[A][B]}+\frac{[A]K_B}{[B]K_A}+\frac{[A][B]K_B}{[A][B]K_B}} = \frac{1}{\frac{K_B}{[B]}+\frac{[A]K_B}{[B]K_A}+1} = \frac{[B]}{K_B+\frac{[A]K_B}{K_A}+[B]}$$

$$[BR] = R_T\frac{[B]K_A}{[A]K_B+K_A([B]+K_B)}$$

$$\frac{[R]}{[R_T]} = \frac{\frac{1}{[A]}}{\frac{1}{[A]}+\frac{1}{K_A}+\frac{[B]}{[A]K_B}} = \frac{1}{\frac{[A]}{[A]}+\frac{[A]}{K_A}+\frac{[A][B]}{[A]K_B}} = \frac{1}{1+\frac{[A]}{K_A}+\frac{[B]}{K_B}}$$

$$[R] = [R_T]\frac{K_AK_B}{[A]K_B+K_A([B]+K_B)}$$

*Functional response:*

$$E = E_{MAX}\frac{[AR]}{K_E+[AR]} = E_{MAX}\frac{\frac{[R_T][A]K_B}{[A]K_B+K_A([B]+K_B)}}{K_E+\frac{[R_T][A]K_B}{[A]K_B+K_A([B]+K_B)}}$$

$$E = E_{MAX}\frac{[R_T][A]K_B}{[R_T][A]K_B+K_E[A]K_B+K_E[B]K_A+K_EK_AK_B}$$

$$E = E_{MAX}\frac{\tau[A]K_B}{\tau[A]K_B+[A]K_B+[B]K_A+K_AK_B} = E_{MAX}\frac{\tau[A]K_B}{[A]K_B(\tau+1)+K_A([B]+K_B)}$$

$$EC_{50} = \frac{K_A}{\tau+1}\times\frac{[B]+K_B}{K_B}$$

$$EC_{50} = \frac{K_A}{\tau+1}\times(1+\frac{[B]}{K_B})$$

### Operational model of dualsteric modulation

*Binding:*

[AR] as fraction of receptors:

$$\frac{[AR]}{[R_T]} = \frac{[AR]}{[R]+[AR]+[RB]+[ARB]+[BR]+[BRB]}$$

Multiplying the numerator and denominator of the fraction on the right side by 1/[A][R] gives:

$$\frac{[AR]}{[R_T]} = \frac{\frac{[AR]}{[A][R]}}{\frac{[R]}{[A][R]}+\frac{[AR]}{[A][R]}+\frac{[RB]}{[A][R]}+\frac{[ARB]}{[A][R]}+\frac{[BR]}{[A][R]}+\frac{[BRB]}{[A][R]}}$$

After substitution:

$$\frac{[AR]}{[R_T]} = \frac{\frac{1}{K_A}}{\frac{1}{[A]}+\frac{1}{K_A}+\frac{[B]}{[A]K_B}+\frac{\alpha[B]}{K_A K_B}+\frac{[B]}{[A]K_C}+\frac{\beta[B]}{K_C K_B}}$$

$$\frac{[AR]}{[R_T]} = \frac{1}{\frac{K_A}{[A]}+1+\frac{[B]K_A}{[A]K_B}+\frac{\alpha[B]}{K_B}+\frac{[B]K_A}{[A]K_C}+\frac{\beta[B]K_A}{K_C K_B}}$$

$$\frac{[AR]}{[R_T]} = \frac{K_B K_C[A]}{K_A K_B K_C+K_B K_C[A]+K_A K_C[B]+\alpha K_C[A][B]+K_A K_B[B]+\beta K_A[A][B]}$$

[ARB] as fraction of receptors:

$$\frac{[ARB]}{[R_T]} = \frac{[ARB]}{[R]+[AR]+[RB]+[ARB]+[BR]+[BRB]}$$

Multiplying the numerator and denominator of the fraction on the right side by 1/[A][R] gives:

$$\frac{[ARB]}{[R_T]} = \frac{\frac{[ARB]}{[A][R]}}{\frac{[R]}{[A][R]}+\frac{[AR]}{[A][R]}+\frac{[RB]}{[A][R]}+\frac{[ARB]}{[A][R]}+\frac{[BR]}{[A][R]}+\frac{[BRB]}{[A][R]}}$$

After substitution:

$$\frac{[ARB]}{[R_T]} = \frac{\frac{\alpha[B]}{K_A K_B}}{\frac{1}{[A]}+\frac{1}{K_A}+\frac{[B]}{[A]K_B}+\frac{\alpha[B]}{K_A K_B}+\frac{[B]}{[A]K_C}+\frac{\beta[B]}{K_C K_B}}$$

$$\frac{[ARB]}{[R_T]} = \frac{\alpha[B]}{\frac{K_A K_B}{[A]}+K_B+\frac{[B]K_A}{[A]}+\alpha[B]+\frac{[B]K_A K_B}{[A]K_C}+\frac{\beta K_A[B]}{K_C}}$$

$$\frac{[ARB]}{[R_T]} = \frac{\alpha K_C[A][B]}{K_A K_B K_C+K_B K_C[A]+K_A K_C[B]+\alpha K_C[A][B]+K_A K_B[B]+\beta K_A[A][B]}$$

*Functional response:*

Functional response to [AR]

$$E = E_{MAX}\frac{[AR]}{K_E+[AR]}$$

For

$$\varphi = K_A K_B K_C + K_B K_C[A] + K_A K_C[B] + \alpha K_C[A][B] + K_A K_B[B] + \beta K_A[A][B]$$

$$E = E_{MAX}\frac{\frac{[R_T]K_B K_C[A]}{\varphi}}{K_E+\frac{[R_T K_B K_C[A]}{\varphi}}$$

$$E = E_{MAX}\frac{[R_T]K_B K_C[A]}{K_E\varphi+[R_T]K_B K_C[A]}$$

After division by $K_E$ and substitution $\tau=[R_T]/K_E$:

$$E = E_{MAX}\frac{\tau K_B K_C[A]}{\varphi+\tau K_B K_C[A]}$$

$$E = E_{MAX}\frac{\tau K_B K_C[A]}{K_A K_B K_C+K_B K_C[A]+K_A K_C[B]+\alpha K_C[A][B]+K_A K_B[B]+\beta K_A[A][B]+\tau K_B K_C[A]}$$

Functional response to [AR]+[ARB]

$$E = E_{MAX}\frac{[AR]+\gamma[ARB]}{K_E+[AR]+\gamma[ARB]}$$

For

$$\varphi = K_A K_B K_C + K_B K_C[A] + K_A K_C[B] + \alpha K_C[A][B] + K_A K_B[B] + \beta K_A[A][B]$$

$$E = E_{MAX}\frac{\frac{[R_T]K_B K_C[A]}{\varphi}+\frac{[R_T]\gamma\alpha K_C[A][B]}{\varphi}}{K_E+\frac{[R_T K_B K_C[A]}{\varphi}+\frac{[R_T]\gamma\alpha K_C[A][B]}{\varphi}}$$

After division by $K_E$ and substitution $\tau=[R_T]/K_E$:

$$E = E_{MAX}\frac{\tau K_B K_C[A]+\tau\gamma\alpha K_C[A][B]}{\varphi+\tau K_B K_C[A]+\tau\gamma K_C[A][B]}$$

$$E = E_{MAX}\frac{\tau K_C[A](K_B+\gamma\alpha[B])}{K_A K_B K_C+K_B K_C[A]+K_A K_C[B]+\alpha K_C[A][B]+K_A K_B[B]+\beta K_A[A][B]+\tau K_C[A](K_B+\gamma\alpha[B])}$$

### Implementation of Python functions and classes

#### Single curve fitting

Example of implementation of Equation 3

```
#get estimates & bounds
IC501_estim = x_min + (x_max - x_min) * 0.25
IC501_min = x_min + 0.5
IC501_max = x_min + (x_max - x_min) * 0.6
IC502_estim = x_min + (x_max - x_min) * 0.75
IC502_min = x_min + (x_max - x_min) * 0.4
IC502_max = x_max - 0.5
F2_estim = 50
F2_min = 0
F2_max = 100
#fit data
p0 = np.array([IC501_estim, IC502_estim, F2_estim])
def func(x, IC501, IC502, F2):
 return 100 - (100 - F2)*10**x / (10**IC501 + 10**x) - F2*10**x / (10**IC502 + 10**x)
popt, pcov = opt.curve_fit(func, x_data, y_data, p0, bounds=([IC501_min,IC502_min,
F2_min],[IC501_max,IC502_max, F2_max]))
perr = np.sqrt(np.diag(pcov))
IC501_calc = (popt[0])
IC502_calc = (popt[1])
F2_calc = (popt[2])
IC501_err = (perr[0])
IC502_err = (perr[1])
F2_err = (perr[2])
```

#### Global 3D-fit

Example of implementation of Equation 13

```
#Enter data files
selected_0 = 'ACh_000.dat'
selected_1 = 'ACh_001.dat'
```

```
selected_2 = 'ACh_003.dat'
selected_3 = 'ACh_010.dat'
selected_4 = 'ACh_030.dat'
selected_5 = 'ACh_100.dat'

#Enter parameters
Emax = 1
basal = 0
KA = 2.04e-7
tauA = 2.18
B0 = 0.000
B1 = 1e-7
B2 = 3e-7
B3 = 1e-6
B4 = 3e-6
B5 = 1e-5

#Set estimates and bounds
KB_ini = 1e-6
KB_min = 1e-9
KB_max = 1e-3
KC_ini = 1e-4
KC_min = 1e-9
KC_max = 1e-3
alpha_ini = 2
alpha_min = 0.01
alpha_max = 100
beta_ini = 0.15
beta_min =0.01
beta_max = 100
delta_ini = 3
delta_min = 0.01
delta_max = 100

#Load data
data_0 = np.loadtxt(selected_0)
data_1 = np.loadtxt(selected_1)
data_2 = np.loadtxt(selected_2)
data_3 = np.loadtxt(selected_3)
data_4 = np.loadtxt(selected_4)
data_5 = np.loadtxt(selected_5)

#Create 3D data set
data_x_0 = data_0[:,0]
data_y_0 = np.linspace(B0,B0,len(data_0))
data_z_0 = data_0[:,1]
data_zerr_0 = data_0[:,2]
data_x_1 = data_1[:,0]
```

```
data_y_1 = np.linspace(B1,B1,len(data_1))
data_z_1 = data_1[:,1]
data_zerr_1 = data_1[:,2]
data_x_2 = data_2[:,0]
data_y_2 = np.linspace(B2,B2,len(data_2))
data_z_2 = data_2[:,1]
data_zerr_2 = data_2[:,2]
data_x_3 = data_3[:,0]
data_y_3 = np.linspace(B3,B3,len(data_3))
data_z_3 = data_3[:,1]
data_zerr_3 = data_3[:,2]
data_x_4 = data_4[:,0]
data_y_4 = np.linspace(B4,B4,len(data_4))
data_z_4 = data_4[:,1]
data_zerr_4 = data_4[:,2]
data_x_5 = data_5[:,0]
data_y_5 = np.linspace(B5,B5,len(data_5))
data_z_5 = data_5[:,1]
data_zerr_5 = data_5[:,2]
x_data = np.concatenate((data_x_0,data_x_1,data_x_2,data_x_3,data_x_4,data_x_5),axis=None)
y_data = np.concatenate((data_y_0,data_y_1,data_y_2,data_y_3,data_y_4,data_y_5),axis=None)
z_data = np.concatenate((data_z_0,data_z_1,data_z_2,data_z_3,data_z_4,data_z_5),axis=None)
z_data_err =
np.concatenate((data_zerr_0,data_zerr_1,data_zerr_2,data_zerr_3,data_zerr_4,data_zerr_5),axis=No
ne)

#fit data
#initial parameter estimates
p1 = [KB_ini, KC_ini, alpha_ini, beta_ini, delta_ini]
#parameter bounds
p1_bounds =
([KB_min,KC_min,alpha_min,beta_min,delta_min],[KB_max,KC_max,alpha_max,beta_max,delta_max
])
#Global fit function
def func_glob(x, y, p):
    KB, KC, alpha, beta, delta = p
    return basal + Emax *
tauA*KC*(10**x)*(KB+delta*alpha*y)/(KA*KB*KC+KB*KC*(10**x)+KA*KC*y+alpha*KC*(10**x)*y+K
A*KB*y+beta*KA*(10**x)*y+tauA*KC*(10**x)*(KB+delta*alpha*y))
#Error function
def err(p, x, y, z):
    return func_glob(x, y, p) - z
pg_opt = opt.least_squares(fun=err, x0=p1, bounds=p1_bounds, args=(x_data, y_data, z_data))
#Calculate the residuals and the variance (mse)
residuals = pg_opt.fun
dof = len(residuals) - len(pg_opt)  # Degrees of freedom
mse = np.sum(residuals**2) / dof  # Mean squared error
#Extract the Jacobian from the result object
```

```
jacobian = pg_opt.jac
#Calculate the Covariance Matrix
try:
    cov = np.linalg.inv(jacobian.T @ jacobian) * mse
except np.linalg.LinAlgError:
    # Use pseudoinverse if the matrix is singular
    cov = np.linalg.pinv(jacobian.T @ jacobian) * mse
#Extract Standard Errors (square root of the diagonal elements)
st_errs = np.sqrt(np.diag(cov))
KB_calc = (pg_opt.x[0])
KB_err = (st_errs[0])
KC_calc = (pg_opt.x[1])
KC_err = (st_errs[1])
alpha_calc = (pg_opt.x[2])
alpha_err = (st_errs[2])
beta_calc = (pg_opt.x[3])
beta_err = (st_errs[3])
delta_calc = (pg_opt.x[4])
delta_err = (st_errs[4])
```

### Model comparison by F-test and AICc

Example of comparison of Equations 2 and 3

```
class ModelComparison:
    """
    Compare up to 3 models using the F-test and AICc criteria.

    Models are compared based on:
    1. Extra Sum-of-Squares F-test (for nested models)
    2. Akaike Information Criterion corrected (AICc)
    3. Residual diagnostics

    Attributes:
        data_x (np.ndarray): Independent variable (ligand concentration, log scale)
        data_y (np.ndarray): Dependent variable (response/occupancy)
        models (dict): Dictionary of model names and their callable functions
        fits (dict): Fitted parameters for each model
        residuals (dict): Residuals for each model
        sum_of_squares (dict): Sum of squared residuals (SS_res)
        statistics (pd.DataFrame): Comparison statistics (AIC, AICc, F-test)
    """

    def __init__(self, data_x, data_y):
        """
        Initialise comparison object.

        Parameters:
```

```
        data_x (array-like): Independent variable values
        data_y (array-like): Dependent variable values (observations)
    """
    self.data_x = np.asarray(data_x)
    self.data_y = np.asarray(data_y)
    self.n_points = len(self.data_y)

    self.models = {}
    self.fits = {}
    self.residuals = {}
    self.sum_of_squares = {}
    self.log_likelihood = {}
    self.statistics = None

def add_model(self, name, model_func):
    """
    Add a model to compare.

    Parameters:
        name (str): Model name (e.g., 'One-Site', 'Two-Site', OMA)
        model_func (callable): Function f(x, *params) -> y
    """
    if len(self.models) >= 3:
        raise ValueError("Maximum 3 models allowed")
    self.models[name] = model_func

def fit_model(self, name, p0, bounds=(-np.inf, np.inf), max_nfev=10000):
    """
    Fit a specific model to data using curve_fit.

    Parameters:
        name (str): Model name (must be added via add_model())
        p0 (array-like): Initial parameter guesses
        bounds (tuple): (lower, upper) bounds for parameters
        max_nfev (int): Maximum number of function evaluations

    Returns:
        popt (np.ndarray): Optimised parameters
        pcov (np.ndarray): Covariance matrix
        residuals (np.ndarray): Residuals (data_y - fitted_y)
    """
    if name not in self.models:
        raise ValueError(f"Model '{name}' not found. Add it with add_model().")

    model = self.models[name]

    try:
        popt, pcov = curve_fit(
```

```
            model,
            self.data_x,
            self.data_y,
            p0=p0,
            bounds=bounds,
            maxfev=max_nfev
        )
    except RuntimeError as e:
        warnings.warn(f"Fitting failed for {name}: {e}")
        return None, None, None

    # Calculate residuals
    y_pred = model(self.data_x, *popt)
    residuals_arr = self.data_y - y_pred

    # Store results
    self.fits[name] = {
        'popt': popt,
        'pcov': pcov,
        'n_params': len(popt)
    }
    self.residuals[name] = residuals_arr

    # Calculate sum of squared residuals
    ss_res = np.sum(residuals_arr ** 2)
    self.sum_of_squares[name] = ss_res

    # Calculate log-likelihood (assuming normal errors)
    mse = ss_res / self.n_points
    self.log_likelihood[name] = -0.5 * (
        self.n_points * np.log(2 * np.pi * mse) + ss_res / mse
    )
     return popt, pcov, residuals_arr

def fit_all_models(self, initial_guesses, bounds=None):
    """
    Fit all added models with the provided initial guesses.

    Parameters:
        initial_guesses (dict): {model_name: p0_array}
        bounds (dict): {model_name: (lower, upper)} or None

    Returns:
        success (bool): True if all models fitted successfully
    """
    if bounds is None:
        bounds = {}
```

```
        success = True
        for name in self.models:
            if name not in initial_guesses:
                raise ValueError(f"Initial guess missing for {name}")

            bnds = bounds.get(name, (-np.inf, np.inf))
            result = self.fit_model(name, initial_guesses[name], bounds=bnds)

            if result[0] is None:
                success = False

        if success:
            self._calculate_statistics()
        return success

    def _calculate_statistics(self):
        """
        Calculate AIC, AICc, and prepare F-test comparisons.
        """
        stats_list = []

        for name in self.models:
            fit_info = self.fits[name]
            k = fit_info['n_params']
            ss_res = self.sum_of_squares[name]
            ll = self.log_likelihood[name]

            # AIC = -2*LL + 2*k
            aic = -2 * ll + 2 * k

            # AICc = AIC + 2*k*(k+1)/(n-k-1)
            if self.n_points - k - 1 > 0:
                aic_c = aic + 2 * k * (k + 1) / (self.n_points - k - 1)
            else:
                aic_c = np.inf
                warnings.warn(
                    f"Model '{name}': n < k+1. Cannot calculate AICc. "
                    "Need more data points."
                )

            stats_list.append({
                'Model': name,
                'n_params': k,
                'SS_res': ss_res,
                'AIC': aic,
                'AICc': aic_c,
                'ΔAICc': np.nan, # Will update after comparing
                'log_likelihood': ll
```

```
        })

    self.statistics = pd.DataFrame(stats_list)

    # Calculate ΔAICc relative to minimum AICc
    min_aicc = self.statistics['AICc'].min()
    self.statistics['ΔAICc'] = self.statistics['AICc'] - min_aicc

    # Sort by AICc
    self.statistics = self.statistics.sort_values('AICc').reset_index(drop=True)

def f_test(self, model_simple, model_complex):
    """
    Perform an extra sum-of-squares F-test for nested models.

    Use this when model_complex is a constrained version of model_simple.

    Parameters:
        model_simple (str): Name of simpler (constrained) model
        model_complex (str): Name of complex (full) model

    Returns:
        f_stat (float): F-statistic
        p_value (float): Two-tailed p-value
        df_num (int): Numerator degrees of freedom
        df_denom (int): Denominator degrees of freedom
        interpretation (str): Text interpretation
    """
    if model_simple not in self.fits or model_complex not in self.fits:
        raise ValueError("Both models must be fitted first.")

    ss_simple = self.sum_of_squares[model_simple]
    ss_complex = self.sum_of_squares[model_complex]

    k_simple = self.fits[model_simple]['n_params']
    k_complex = self.fits[model_complex]['n_params']

    if k_complex <= k_simple:
        raise ValueError(
            f"'{model_complex}' must have more parameters than '{model_simple}'"
        )

    if ss_complex > ss_simple:
        warnings.warn(
            "The complex model has a worse fit than the simple model. "
            "Check that models are nested correctly."
        )
```

```
        # F-test: F = [(SS_simple - SS_complex) / Δdf] / (SS_complex / df_complex)
        df_num = k_complex - k_simple
        df_denom = self.n_points - k_complex

        if df_denom <= 0:
            raise ValueError(
                f"Insufficient degrees of freedom: {df_denom}. "
                "Need n > k_complex."
            )

        f_stat = ((ss_simple - ss_complex) / df_num) / (ss_complex / df_denom)
        p_value = 1 - f_dist.cdf(f_stat, df_num, df_denom)

        # Interpretation
        if p_value < 0.05:
            interp = f"Complex model significantly better (P={p_value:.4f})"
        elif p_value < 0.157:
            interp = f"Weak evidence for complex model (P={p_value:.4f}, AIC threshold)"
        else:
            interp = f"Simple model adequate (P={p_value:.4f})"

        return f_stat, p_value, df_num, df_denom, interp

    def summary(self, include_fits=False):
        """
        Print a summary of model comparison.

        Parameters:
            include_fits (bool): Include fitted parameters in output
        """
        print("\n" + "="*80)
        print("MODEL COMPARISON SUMMARY")
        print("="*80)
        print(f"Data points: {self.n_points}")
        print(f"Models: {list(self.models.keys())}")
        print("\n" + "-"*80)

        # AICc table
        print("\nINFORMATION CRITERIA:")
        print(self.statistics[['Model', 'n_params', 'SS_res', 'AICc', 'ΔAICc']].to_string(index=False))

        # AICc interpretation
        print("\nAICc INTERPRETATION:")
        print("  ΔAICc = 0–2:   Weak support for simpler model; models similarly supported")
        print("  ΔAICc = 4–7:   Substantial evidence for lower AICc model")
        print("  ΔAICc > 10:    Very strong support for lower AICc model")

        # Residuals diagnostics
```

```
        print("\n" + "-"*80)
        print("\nRESIDUAL DIAGNOSTICS:")
        for name in self.models:
            residuals_arr = self.residuals[name]
            print(f"\n  {name}:")
            print(f"    Mean (should be ~0):          {np.mean(residuals_arr):8.2e}")
            print(f"    Std Dev:                      {np.std(residuals_arr):8.2e}")
            print(f"    Min / Max:                    {np.min(residuals_arr):8.2e} / {np.max(residuals_arr):8.2e}")
            print(f"    Shapiro-Wilk p-value:         {self._shapiro_wilk(residuals_arr):8.4f}")

        # Fitted parameters
        if include_fits:
            print("\n" + "-"*80)
            print("\nFITTED PARAMETERS:")
            for name in self.models:
                popt = self.fits[name]['popt']
                pcov = self.fits[name]['pcov']
                stderr = np.sqrt(np.diag(pcov))
                print(f"\n  {name}:")
                for i, (p, se) in enumerate(zip(popt, stderr)):
                    print(f"    param[{i}]:  {p:12.6f} ± {se:12.6e}")

        print("\n" + "="*80 + "\n")

    def _shapiro_wilk(self, residuals):
        """
        Shapiro-Wilk test for normality of residuals.
        (Simple approximation; use scipy.stats.shapiro for full version)
        """
        try:
            from scipy.stats import shapiro
            _, p_val = shapiro(residuals)
            return p_val
        except:
            return np.nan

    def plot_fits(self, ax=None, conc_log_range=None):
        """
        Plot data and fitted curves.

        Parameters:
            ax (matplotlib.axes.Axes): Axes object. If None, creates new figure
            conc_log_range (tuple): (log_min, log_max) for smooth curve plotting

        Returns:
            ax (matplotlib.axes.Axes): Axes with plots
        """
        try:
```

```
            import matplotlib.pyplot as plt
        except ImportError:
            raise ImportError("matplotlib required for plotting. Install: pip install matplotlib")

        if ax is None:
            fig, ax = plt.subplots(figsize=(10, 6))

        # Plot data
        ax.scatter(self.data_x, self.data_y, s=50, color='black',
                   alpha=0.6, label='Data', zorder=3)

        # Determine x range for smooth curves
        if conc_log_range is None:
            log_min, log_max = np.log10(self.data_x[self.data_x > 0]).min() - 0.5, \
                               np.log10(self.data_x[self.data_x > 0]).max() + 0.5
        else:
            log_min, log_max = conc_log_range

        x_smooth_log = np.linspace(log_min, log_max, 200)
        x_smooth = 10 ** x_smooth_log

        colors = ['red', 'blue', 'green']
        for (name, model), color in zip(self.models.items(), colors):
            if name in self.fits:
                popt = self.fits[name]['popt']
                y_smooth = model(x_smooth, *popt)
                ax.plot(x_smooth, y_smooth, color=color, linewidth=2,
                        label=f"{name}
(AICc={self.statistics[self.statistics['Model']==name]['AICc'].values[0]:.1f})")

        ax.set_xscale('log')
        ax.set_xlabel('Ligand Concentration (log scale)', fontsize=12)
        ax.set_ylabel('Response / Occupancy', fontsize=12)
        ax.set_title('Ligand-Receptor Binding Models', fontsize=13, fontweight='bold')
        ax.legend(fontsize=10)
        ax.grid(True, alpha=0.3)

        return ax

    def plot_residuals(self, ax=None):
        """
        Plot residuals for each model.

        Parameters:
            ax (matplotlib.axes.Axes): Axes object. If None, creates new figure

        Returns:
            ax (matplotlib.axes.Axes): Axes with residual plots
```

```
"""
try:
    import matplotlib.pyplot as plt
except ImportError:
    raise ImportError("matplotlib required for plotting. Install: pip install matplotlib")

if ax is None:
    n_models = len(self.models)
    fig, axes = plt.subplots(1, n_models, figsize=(5*n_models, 4))
    if n_models == 1:
        axes = [axes]
else:
    axes = [ax]

colors = ['red', 'blue', 'green']
for (name, residuals_arr), color, ax_i in zip(self.residuals.items(), colors, axes):
    ax_i.scatter(self.data_x, residuals_arr, s=50, color=color, alpha=0.6)
    ax_i.axhline(y=0, color='black', linestyle='--', linewidth=1)
    ax_i.set_xscale('log')
    ax_i.set_xlabel('Ligand Concentration (log scale)', fontsize=11)
    ax_i.set_ylabel('Residuals', fontsize=11)
    ax_i.set_title(f'{name}', fontsize=12, fontweight='bold')
    ax_i.grid(True, alpha=0.3)

return ax_i if ax is not None else axes
```
